# Pleiotropy and Disease Interactors: The Dual Nature of Genes Linking Ageing and Ageing-related Diseases

**DOI:** 10.1101/2024.05.20.594975

**Authors:** Gustavo Daniel Vega Magdaleno, Joao Pedro de Magalhaes

## Abstract

Ageing-related diseases (ARDs) exhibit a broad spectrum of phenotypes yet consistently increase in incidence with advancing age. This suggests that, despite their diversity, ARDs could potentially share common biological processes deeply rooted in the mechanisms of ageing, presenting opportunities for unified therapeutic strategies. Using a network approach, we analysed gene proximity to 52 ARDs from the UK Biobank, integrating with protein-protein interaction (PPI), gene coexpression, KEGG pathways, and ageing-related genes. Interestingly, while most ageing-related genes did not associate with ARDs, they were closer to multiple ARDs than random genes. This was mainly due to indirect connections to diverse Communities of ARDs (ARCs), what we call *iARC-Interactor*s, implying indirect association to multiple ARDs through interaction with ARD-related genes, primarily via PPI and KEGG. Genes that are associated with multiple ARCs, *i.e*., *Pleiotropic* genes, were predominantly related to immunological disorders. We found a polarizing effect. When compared to multiple ageing- and disease-related genes, high *Pleiotropic* genes showed the highest tissue specificity and lowest coexpression with themselves and other diseases. In contrast, high i*ARC-interactive* genes (as those of ageing) significantly displayed the exact opposite effects, suggesting two mechanisms for genes to affect multiple ARDs, one operating through modulatory genes that simultaneously affect numerous tissues and processes; and another rather specialised, affecting single tissues that are widespread across the body, as potentially occurring in autoimmune diseases. Lastly, we used Machine Learning (ML) to predict potentially novel ageing-related genes based on each network’s *iARC-Interactions* and genes’ proximity to ARDs. PPI and KEGG showed the best performance with their top candidate genes enriched for regulation of protein metabolic process, protein stabilization, positive regulation of developmental process, and cellular response to chemical stimulus. This work paints a deeper picture of the multiple types of interactions between ageing-related processes and ARDs.

## Introduction

Ageing-related diseases (ARDs) like stroke (Yousufuddin *et al*., 2019), cancer (Chatsirisupachai et al., 2021), diabetes (Munshi et al., 2020), and dementia (Brayne *et al*., 2017) may seem independent on first sight. Still, they are all strongly associated with advancing age (Nikkoli *et al*., 2012), suggesting potential shared mechanisms rooted in the ageing process. Recent research highlights potential commonalities in underlying ageing-related mechanisms such as telomere shortening (Gruber *et al*., 2021), cellular senescence (Childs *et al*., 2015), DNA repair (Yousefzadeh *et al*., 2021), epigenetic changes (Saul & Kosinsky, 2021; Horvath & Raj, 2018), mitochondrial issues (Lane *et al*., 2015; Sun *et al*., 2016), autophagy (Cheon *et al*., 2019), and other debated hallmarks of ageing (López-Otín *et al*., 2013; Guo *et al*., 2022; Gems & de Magalhães, 2021; Keshavarz *et al*., 2023; de Magalhaes, 2024). Furthermore, ARDs interactions with these ageing hallmarks could be better understood through network propagation (Fraser et *al.*, 2022), emphasizing the interconnectedness of ageing-related processes and disease susceptibility.

Previous studies have delved into the relationship of ageing with ARDs, highlighting the role of signalling proteins in linking ARDs to lifespan (Wolfson et *al.*, 2009), and shared genes and pathways in the interplay between cellular senescence, ageing, and ARDs (Tacutu *et al*., 2011). Research also highlights the distinct properties of ageing-related genes in protein-protein interaction (PPI) networks and their connections to ARDs (Wang *et al*., 2009), along with the notable yet evolutionarily reducing overlap between human ageing and ARDs-associated genes (Fernandes *et al*., 2016). Moreover, different associations have been observed between pro-longevity genes and nervous, musculoskeletal diseases, and anti-longevity genes with cardiovascular diseases, with significant linkages in specific subnetworks suggesting potential predictors for ARDs (Yang *et al*. 2016). Building on these insights, a recent study (Donertas *et al*., 2021) analysed the age-of-onset for 116 diseases using UK Biobank (Bycroft *et al*., 2018), identifying four disease clusters, two of which increase across the population with age. They found that diseases with similar onset ages are genetically more related than expected, even when compared to diseases with similar tissue of occurrence. Comparing GenAge-associated genes with those in ageing-related clusters showed limited overlap, indicating that, while some ageing-related genes are linked to multiple ARDs, more understanding could come from examining indirect connections like protein interactions, gene co-expression, and pathway networks.

The concept of *Pleiotropy*, where a single gene affects multiple phenotypic traits, is important in understanding ARDs (He *et al*., 2016; Donertas *et al*., 2021). An example of this is genes influencing processes like cellular senescence and insulin signalling, which are linked to various ARDs (Tacutu *et al*., 2011; Sun *et al*., 2016). However, *Pleiotropy* in ARDs does not always imply a direct association with ageing, particularly if the ARDs affected are phenotypically similar or share joint tissues or biological processes. For instance, a *Pleiotropic* gene affecting both “Stroke” and “High Cholesterol” might indicate a cardiovascular, rather than an ageing-related, association. To address this, ARDs can be categorised based on the tissue of occurrence, creating what we call Ageing-Related Communities (ARCs). For example, “Stroke” and “High Cholesterol” fall under a “Cardiovascular Diseases” ARC, while “Arthritis” and “Osteoporosis” are in a “Musculoskeletal/Trauma” ARC. This study uses “Communities” and “ARCs” synonymously and defines Disease-related genes as those associated with at least one ARD/ARC.

Some genes indirectly affect multiple ARCs by interacting with ARC-related genes without the need for direct GWAS associations with ARDs. These genes, which we termed *iARC-Interactors* (with “i” for “indirect”), affect others via interactions, such as influencing gene expression or function. Acknowledging the role of indirect interactions can provide a richer understanding of disease pathogenesis and potential therapeutic interventions. However, this perspective has been poorly explored (Yang et *al.*, 2016), lacking analysis that compares it with *Pleiotropy* in ARD research.

The specificity of gene expression in various tissues has profound implications for understanding ageing and ARDs. Evidence suggests that specific genes play differential roles across tissue types and contribute uniquely to ageing. For instance, the FOXO3 gene, a known longevity-associated gene, demonstrates tissue-specific activity, with its function varying between cardiovascular, muscular, and neuronal tissues, thus impacting diverse aspects of ageing (Martins *et al*., 2016). Another study (Yang *et al*., 2015) highlighted that genes associated with ageing are not randomly distributed across tissues but are particularly enriched in specific tissues, such as the brain, skeletal muscle, and immune system. Therefore, further research is needed to understand better how this measure interacts with ageing-related genes.

In this study, we examined how genes relate to 58 ARDs using UK Biobank, combining data from GenAge (Tacutu *et al*., 2018), BioGRID’s PPI (Stark *et al*., 2006), GeneFriends (Raina *et al*., 2023) coexpression networks (at 90% and 95% thresholds), KEGG pathways (Kanehisa *et al*., 2016) (considering varied directionality) and tissue specificity from the Human Ageing Genomic Resources (HAGR) (Tacutu *et al*., 2018; Palmer *et al*., 2021). Intriguingly, although most human and model ageing-related genes depicted little overlap with human ARDs, their indirect interactions spanned multiple ARCs, reaching closer topological proximity to ARCs than random and even ARC-neighbouring genes. Such a trend was attributed to *iARC-Interactors*, predominantly through PPI and KEGG. Genes that are associated with multiple diseases through GWAS *(i.e.*, *Pleiotopic* genes) were rather tied to immunological disorders. We found that high *Pleiotropic* genes tended to exhibit higher tissue specificity and displayed reduced coexpression with themselves as a group and other ARC-related genes. Conversely, the patterns for high *iARD-Interactive* genes were opposite relative to other diseases. This proposes dual mechanisms by which genes influence several ARDs: one via universally impacting genes and another, more targeted, influencing unique yet common tissues akin to some autoimmune conditions. Finally, we used Machine Learning (ML) to identify new ageing-associated genes, considering each network’s *iARD_Interactions* and gene-to-diseases proximity. PPI and KEGG with null directionality performed the best, with coexpression networks being the least effective predictors. The predicted ageing-related genes from PPI and KEGG were enriched for protein metabolic regulation, protein safeguarding, developmental progression boosts, and cellular reactions to specific triggers.

## Results

We explored the direct and indirect interactions between ARDs- and ageing-related genes. The ageing-related genes were splitted in two groups: 307 Human Ageing-Related genes from GenAge and 1157 human homologs of ageing-related genes from model organisms. For ease of notation, we call *GenAge*_*Hum*_ and *GenAge*_*Mod*_ to the human and model organisms databases of GenAge, respectively, while their associated genes are *Ageing*_*Hum*_- and *Ageing*_*Mod*_-related genes. The intersection between the two GenAge groups is 121 genes, which represents 39.4% of human ageing-related genes and 10.4% of ageing-related genes in model organisms. Supplementary Table S1 depicts an enrichment analysis of genes associated with these sets, highlighting *apoptotic process* and *cellular response to chemical stimulus* for human ageing-related genes; *organonitrogen compound metabolic process*, *generation of precursor metabolites and energy*, *macroautophagy* and *organic anion transport* for ageing-related genes at model organisms; and cellular response to stress and response to oxygen-containing compound at the intersection of the two groups.

The ARDs were based on 58 previously reported non-cancer self-reported ARDs in UK Biobank and one phenotype of ageing-related cancer computed by us using UK Biobank as well (Methods – Selection of ARDs and ARCs). All the ARDs (58 in total) and their relationship to ARCs (9 in total) were based on the UK Biobank hierarchical guidelines of diseases (Supplementary Figures S1-S3). Throughout this work, we use the terms “Communities” and “ARCs” interchangeably. We also use the term *Disease*-related genes as any gene associated with at least 1 ARD/ARC.

In its general meaning, the word *Pleiotropy* refers to the association of one gene to multiple phenotypes. In the context of our work, *ARC-Pleiotropy* is the association of a gene to multiple ARCs, and *ARD-Pleiotropy* is the association of a gene to multiple ARDs (Methods - Pleiotropy and Indirect Interactor definitions). When further referring to *Pleiotropy* in this chapter, lacking a label “ARD” or “ARC”, we will allude to *ARC-Pleiotropy.* We make this distinction because instances of *Pleiotropy* involving multiple ARCs would tend to indicate that the gene in question impacts numerous tissues, mirroring the widespread effects of ageing. On the other hand, focusing on ARD *Pleiotropy* could more often result in correlated findings, especially when the ARDs occur within the same tissue or share functional similarities.

Figure 1 and its complement Supplementary Figure S4 show an already well-known fact (Yen *et al*., 2022; Hahn *et al*., 2022; O’Sullivan *et al*., 2022; Donertas *et al*., 2021), Cardiovascular and endocrine/diabetes disorders are notably polygenic with over 1,000 associated genes each and concurrent overlaps with other ARCs. Subsequent categories include musculoskeletal, gastrointestinal, cancer, and neurology, which have between 205 and 410 associated genes, showing varied overlaps. In contrast, haematology/dermatology and renal/urology only showed 16 genes each, mainly overlapping with cardiovascular and endocrine diseases. Immunological systemic disorders lie in between with 128 associated genes.

**Figure 1.**
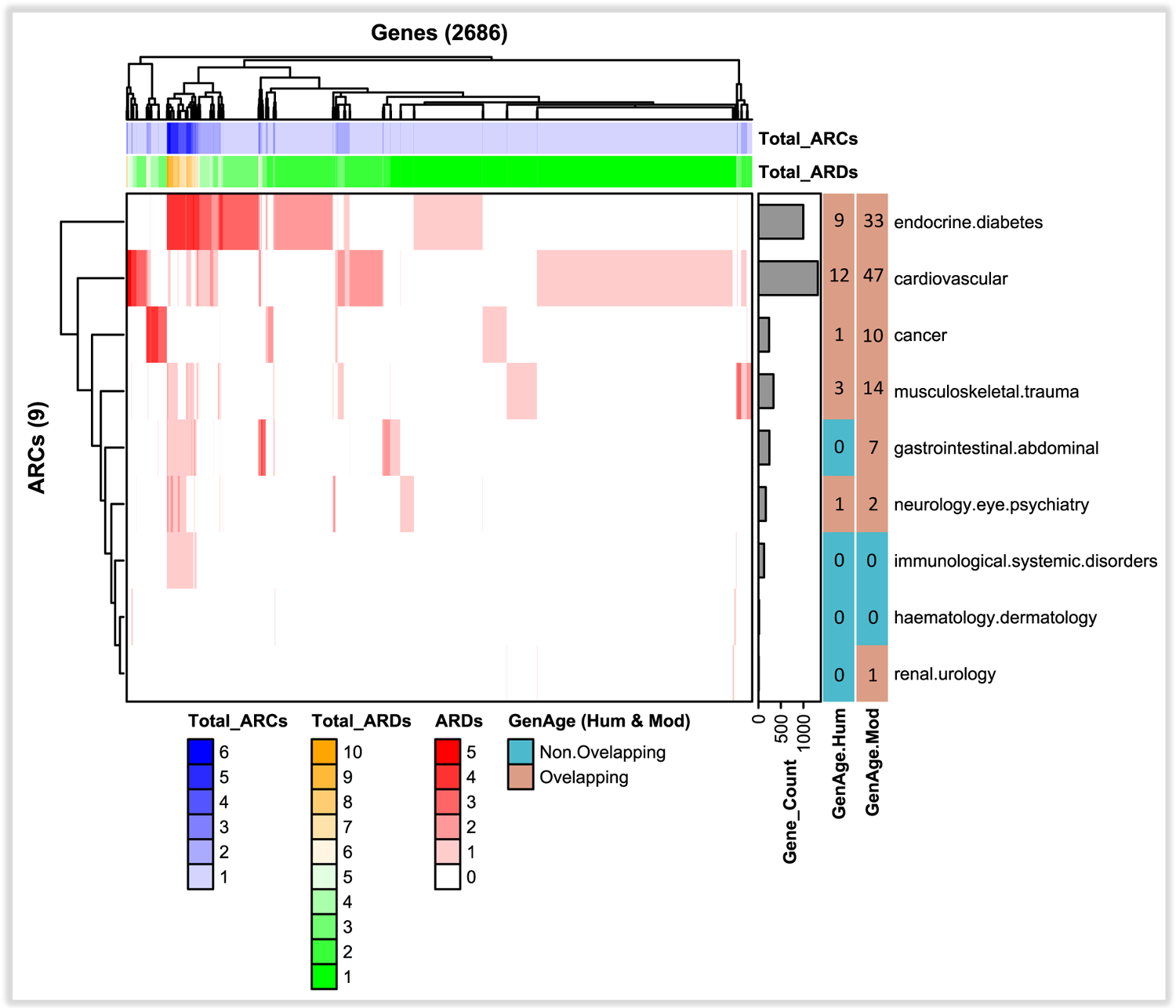
Genetic profiles of ARCs. Rows are ARCs *(i.e.*, a set of ARDs clustered by tissue) and columns are genes associated with ARDs (*i.e*., ageing-related diseases). A gradient from white to red is displayed, with white indicating that a gene does not interact with any diseases, and a stronger red colour representing a higher number of interacting diseases. Endocrine, cardiovascular, cancer, and gastrointestinal communities have a higher red colouration, indicating a greater density of diseases affected by the same gene. On the right side of the graph, a histogram shows the number of genes per region, with cardiovascular having the most genes, followed by endocrine. Additionally, two extra bars indicate whether the genes within that community overlap with any of the two GenAge-associated groups or not, showing that gastrointestinal, immunological, haematological, and renal communities do not overlap with *GenAge*_*Hum*_, whereas *GenAge*_*Mod*_overlapped with more communities except immunological and haematological. At the top of the figure, several heatmaps and bars are displayed. The first bar represents the total number of ARDs for a particular gene, summing across all rows for the same column. The next bar indicates the total number of ARCs affected by the same gene, with a gradient from light blue to dark blue, with darker blue representing more ARCs.

Most *Disease*-related genes tend to associate with one single ARC, except for those associated with immunological diseases, which often overlap with other categories. Genes that GWAS-associated with 2 ARCs, what we call second-level *Pleiotropy* or, simply put, *Pleiotropy_2* genes, were observed across all the communities but primarily cardiovascular, endocrine, and musculoskeletal diseases (Figure 1 and Supplementary Figure S4). All ARCs exhibited this form of gene interaction. *Pleiotropy_3* genes generally included the aforementioned groups but excluded others like immunological diseases.

Remarkably, higher levels of *Pleiotropy* (four to six) were mostly observed in immunological diseases, indicating major ARC-interconnectedness. Such high *Pleiotropic* genes were enriched for both non-strictly (*nucleosome assembly* and *urate transport*) and strictly (*innate immune response in mucosa, T cell receptor signalling pathway* and *antibacterial humoral response*) immune-related biological processes (Supplementary Table S2).

A modest percentage of roughly 12% of the genes at both *GenAge*_*Hum*_ and *GenAge*_*Mod*_ overlapped with ARCs (Figure 2). Most of these overlapping genes correspond to cardiovascular, endocrine, and musculoskeletal diseases (Supplementary Figure S4) and showed different levels of *Pleiotropy*, with the majority at *Pleiotropy_1*. Nevertheless, *GenAge*_*Mod*_ displayed a larger proportion of cardiovascular-, endocrine-diabetes-, and musculoskeletal-trauma-related genes compared to *GenAge*_*Hum*_. The overlap in cancer-related genes was notably higher in *GenAge*_*Mod*_ than *GenAge*_*Hum*_. Lastly. There was minimal overlap for higher *Pleiotropy* levels and no overlap with immunological disorders in both databases.

**Figure 2.**
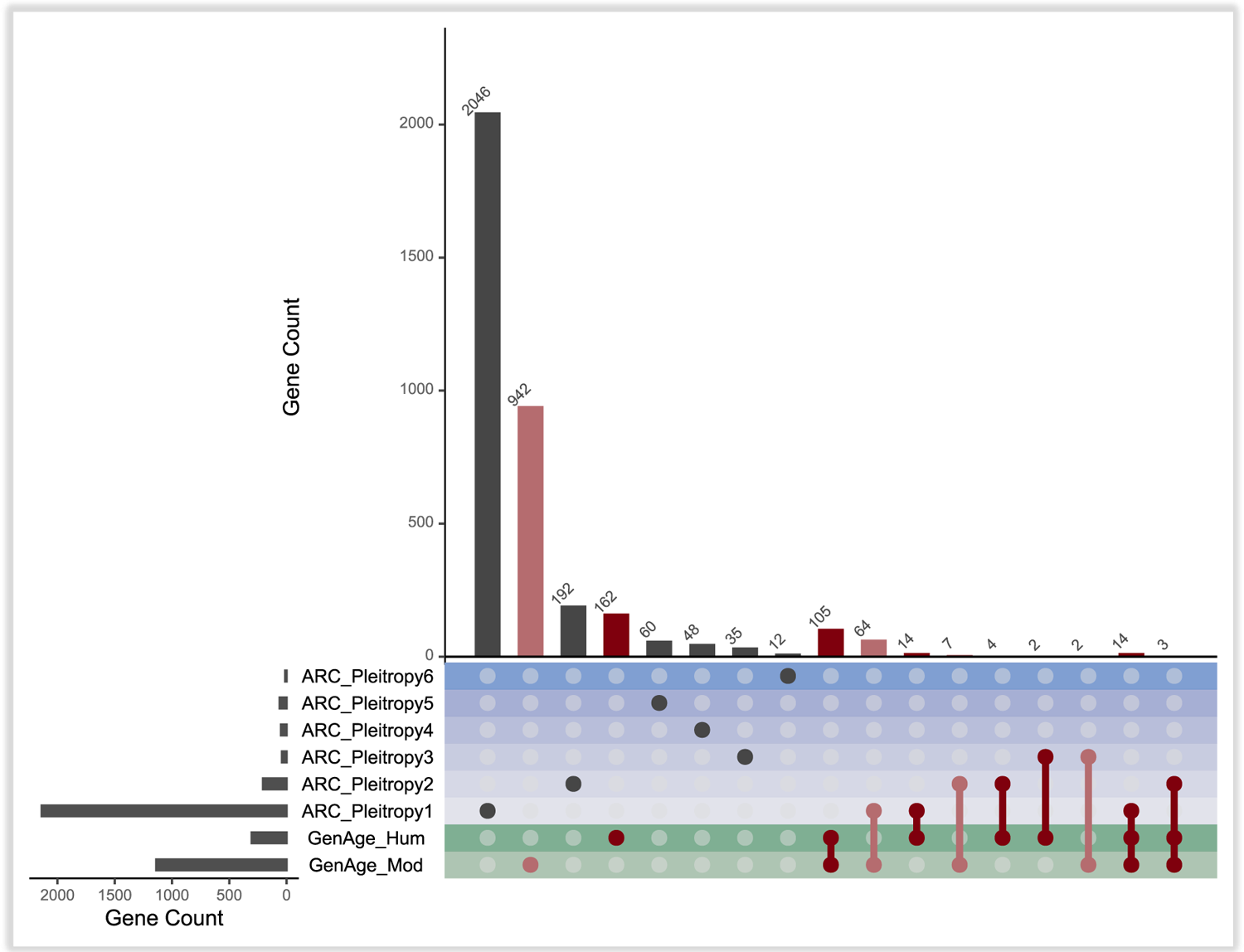
Overlapping between GenAge- and ARC-related genes expanded by their degree of *Pleiotropy*. At row names, we explicitly stated *ARC_Pleiotropy* and assigned a number indicating the count of ARCs with which the genes in the row are associated, as per GWAS data. This ARC association is based on the requirement that each gene must be linked to at least one ARD within the corresponding ARC. For instance, if a gene is associated with ARDs across three different ARCs, then it will be referred to as *ARC_Pleiotropy3*. Red colour was used for highlighting GenAge-related sets. Two green colours were used to highlight the two *GenAge* rows, *GenAge*_*Hum*_and *GenAge*_*Mod*_. An increasing palette of blues was used to highlight the increasing *Pleiotropy* levels. It can be observed that most *GenAge* genes do not overlap with any ARC-related gene at all. The major overlap with ARC-related genes occurs at genes with *Pleiotropy* of 1, followed by decreasing levels of genes with *Pleiotropies* 2 and 3. No higher *Pleiotropies* overlapped with *GenAge*-associated genes.

### ARC Genetic Networks

To better understand the relationships between ageing- and ARDs-related genes, we integrated gene-gene relationships from three databases: BioGRID’s PPI, GeneFriends’ gene coexpression, and the integration of KEGG pathways. As better explained in Methods (Methods ARC networks), the BioGRID’s PPI was straightforwardly integrated; the GeneFriends database was used to create two coexpression networks, referred to as *COX*_90_ and *COX*_95_, based on coexpression thresholds of 90% and 95%, respectively; all the pathways at the KEGG database were integrated as well, disregarding their directionality. The inclusion of genetic interactions to the GWAS-established Gen-ARC’s associations allowed generating what we refer to as *ARC*-networks, which encompass a broader range of genes and indirect associations to the ARCs. There is one *ARC*-network per type of Gen-Gen interaction, summing up to four networks: *ARC.PPI, ARC.COX*_90_, *ARC.COX*_95_ and *ARCC.KEGG*. Supplementary Figures S5, S7, S9 and S11 depict such networks, and Supplementary Figures S6, S8, S10 and S12 illustrate heatmap representations of genetic distances between genes and ARCs, as determined by their position within the *ARC.PPI, ARC.COX*_90_, *ARC.COX*_95_ and *ARC.KEGG* networks, respectively.

### Disease Interactors

Genes were categorized into five groups for *iARC_Interactor* analysis: *GenAge*_*Hum*_, *GenAge*_*Mod*_, *Diseases* (ARCs), *Neighbours* (*i.e.*, neighbour genes of *Disease*-related genes), and *Others* (*i.e*., the remaining genes in the network).Figure 3 illustrates the degree of *iARC-Interactors* associated with genes in each group, *i.e.*, the number of ARCs that can be indirectly reached by the gene through interaction with ARC-related genes (Methods – *Pleiotropy* and Indirect Interactor definitions). This quantity ranges from zero, when no ARC is reached, to nine, when nine ARCs are indirectly reached.

**Figure 3.**
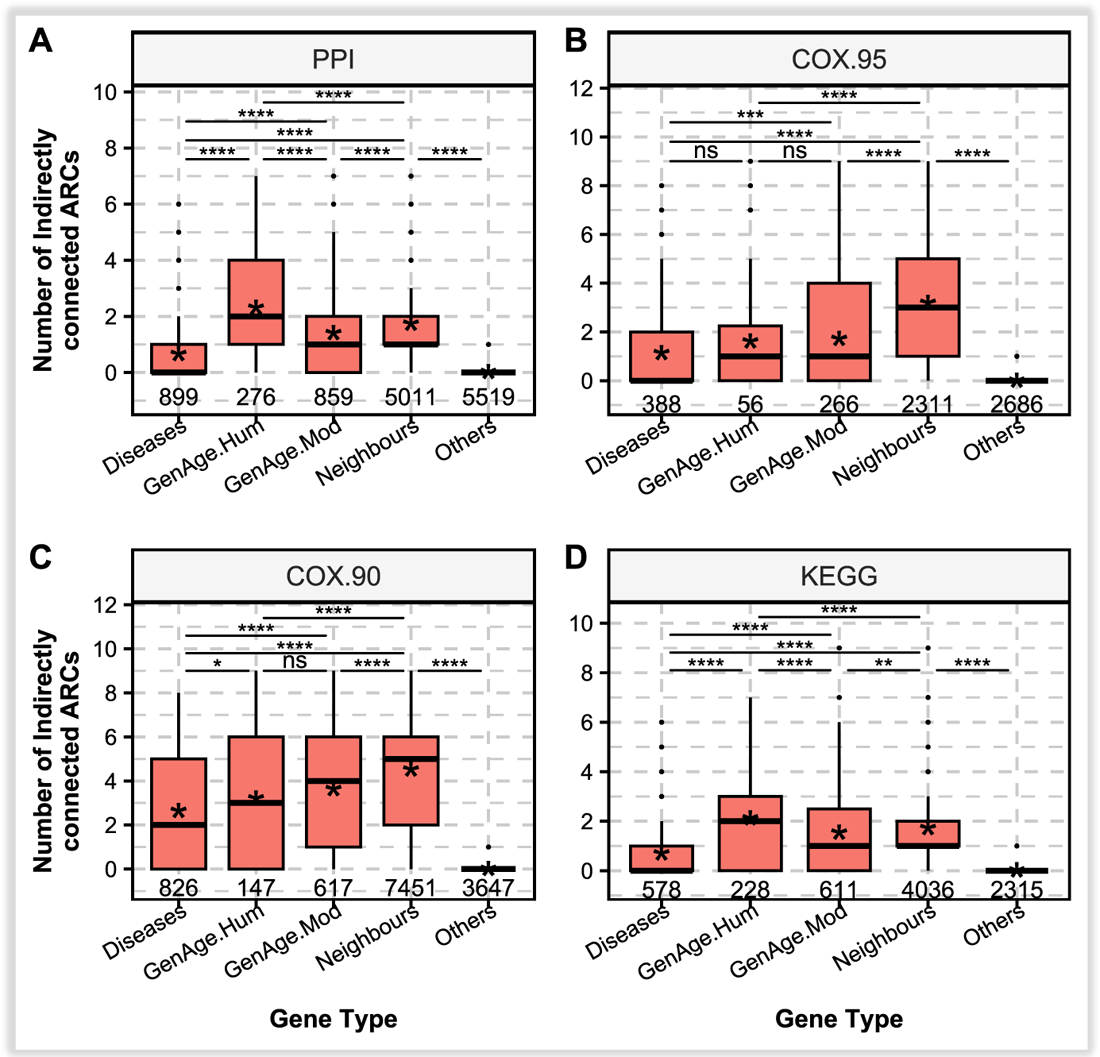
*iARC-Interaction* levels across the following sets of genes: *Diseases, GenAge*_*Hum*_, *GenAge*_*Mod*_, *Neighbours* of diseases, within each corresponding interaction network. Asterisks represent the mean values to help noticing differences in distributions with equal medians. Of note is that human ageing-related genes are more studied, thus inducing bias. This same bias is partially inherited by Neighbours as they contain. **A**. *ARC.PPI* network. **B.** *ARC.COX*_95_ network. **C.** *ARC.COX*_90_ network. **D**. *ARC.KEGG* network. Statistical differences were computed using Wilcoxon Signed-Rank and adjusted for multiple tests using Bonferroni. The statistical significance with respect to others is not shown as it is always significant since, by definition, Others do not indirectly interact with any ARC-related genes, thus all its genes having a value of zero, which is significantly lower than any of the remaining groups.

In the *ARC.PPI* network (Figure 3A), *GenAge*_*Hum*_-related genes stood out as the most pronounced *iARC_interactors*, reaching a median interaction count of two and an average of 2.32. They were followed by *Neighbouring* genes with a median of one and a mean of 1.40, and *GenAge*_*Mod*_ genes took the third position with similar median but a slightly lower mean of 1.38. *Disease*-related genes stayed behind with median at zero and mean of 0.67, while the *Others* category consistently recorded zero interactions. The variations were statistically significant across all groups within these networks.

*GenAge*_*Hum*_ genes also exhibited the broader distribution in their *iARC-Interaction* levels, spanning from zero to seven before outliers. In comparison, *GenAge*_*Mod*_ genes showed a distribution mostly between zero to five, whereas *Disease*-related genes ranged from zero to two. *Neighbours* covered a range from zero to three, reaching an intermediate wide between *GenAge*_*Mod*_ and *Diseases*.

In the *ARC.COX*_95_ and *ARC.COX*_90_ networks (Figure 3B-C), significant mean differences were present in most cases, with the exception of the *GenAge*_*Hum*_-*Diseases* comparison, where the significance was either absent (*ARC.COX*_95_) or minimal (*ARC.COX*_90_). This contrasted with the *GenAge*_*Mod*_-*Diseases* comparison, where *GenAge*_*Mod*_ showed a significant elevation, likely due to its higher gene count compared to *GenAge*_*Hum*_. In the *ARC.COX*_95_ network, both *GenAge*_*Hum*_ and *GenAge*_*Mod*_ genes exhibited a median of one and an average near 1.62, contrasting with *Disease*-related genes, which had a median of zero and a mean of 1.15. In the *ARC.COX*_90_ network, the median values for *GenAge*_*Hum*_-, *GenAge*_*Mod*_-, and *Disease*-related genes approximated three, four, and two, respectively, with corresponding means of 3.22, 3.8, and 2.66. *Neighbouring* genes surpassed all the remaining groups in these networks. In *ARC.COX*_95_, *Neighbours* achieved a median of three and an average of 3.48, and, in *ARC.COX*_90_, they predominantly reached a median of four and an outstanding mean of 4.66. This outcome highlights coexpression networks as the only ones where *Neighbours* exceeded even *GenAge*_*Hum*_-associated genes in the number of indirectly connected ARCs. The *Others* category remained consistently at zero in both network types. Moreover, the distribution range in *ARC.COX*_90_ was remarkably broad, spanning from zero to nine for most groups. In contrast, *ARC.COX*_95_ depicted a wide distribution primarily in *GenAge*_*Mod*_ and *Neighbours*, with *Diseases* and *GenAge*_*Mod*_ generally ranging from zero to five. These variances, however, did not account for a significant difference between *GenAge*_*Hum*_ and *GenAge*_*Mod*_ in *ARC.COX*_95_, as mentioned earlier.

In the *ARC.KEGG* network (Figure 3D), significant differences existed between gene groups as well. *GenAge*_*Hum*_-associated genes held their steady distribution from zero to six/seven, with median at two; and mean at 2.13 (*ARC.KEGG*). *GenAge*_*Mod*_-associated genes held a slightly lower-centered and narrower distribution from zero to five/six, with medians at one (*ARC.KEGG*); and mean at 2.13 (*ARC.KEGG*). *Disease*-related genes at *ARC.KEGG* were 0.73. *Neighbours* genes at all these networks had median 1 with an almost overlapping mean at 1 excepting for the *ARC.KEGG* network whose mean was closer to 2. In this direction, *Neighbouring* genes tended to have similar level of indirectly connected ARCs With *GenAge*_*Mod*_, thus lacking significant difference in all KEGG networks excepting *ARC.KEGG*, where neighbours were slightly higher. Most *GenAge-associated* values in *ARC.KEGG*, ranged from zero to seven. The *Others* consistently scored zero as, by definition, they are not associated with any ARC.

### Intra- and Inter-Community Coexpression

We investigated the self-coexpression (*i.e*., coexpression of genes within each specific ARC, or intra-community coexpression) and cross-coexpression (*i.e*., coexpression of genes associated with two different ARCs, or inter-community coexpression) of various gene groups, either related to specific ARCs, multiple levels of *Pleiotropy*, and the two groups of *GenAge*-associated genes. In the heatmap depicted in Figure 4, the main diagonal values of each square sub-matrix show self-coexpression within gene groups, while the remaining values reflect cross-coexpression between groups. Notably, self-coexpression in ARCs was moderately low (0.18-0.27). However, the *GenAge*_*Mod*_ group displayed a self-coexpression of 0.44, higher than any other group by far. The difference is highly significant (Supplementary Figure S16). *GenAge*_*Hum*_ was the second group with highest self-coexpression, reaching a score of 0.34. Additionally, ageing-related genes consistently showed the highest cross-coexpression with other groups led by model organisms’ genes (∼0.29) and followed by humans (∼0.27). This cross-coexpression with *GenAge*_*Mod*_-associated genes was always higher than the self-coexpression of ARCs, whereas *GenAge*_*Hum*_ was mostly higher as well, although less often and to a lesser extent. Renal and immunological disease ARCs had the lowest cross-coexpression (∼0.15), significantly (Supplementary Figure S16), but renal diseases had a high self-coexpression of 0.25, only surpassed by haematological diseases (0.27).

**Figure 4.**
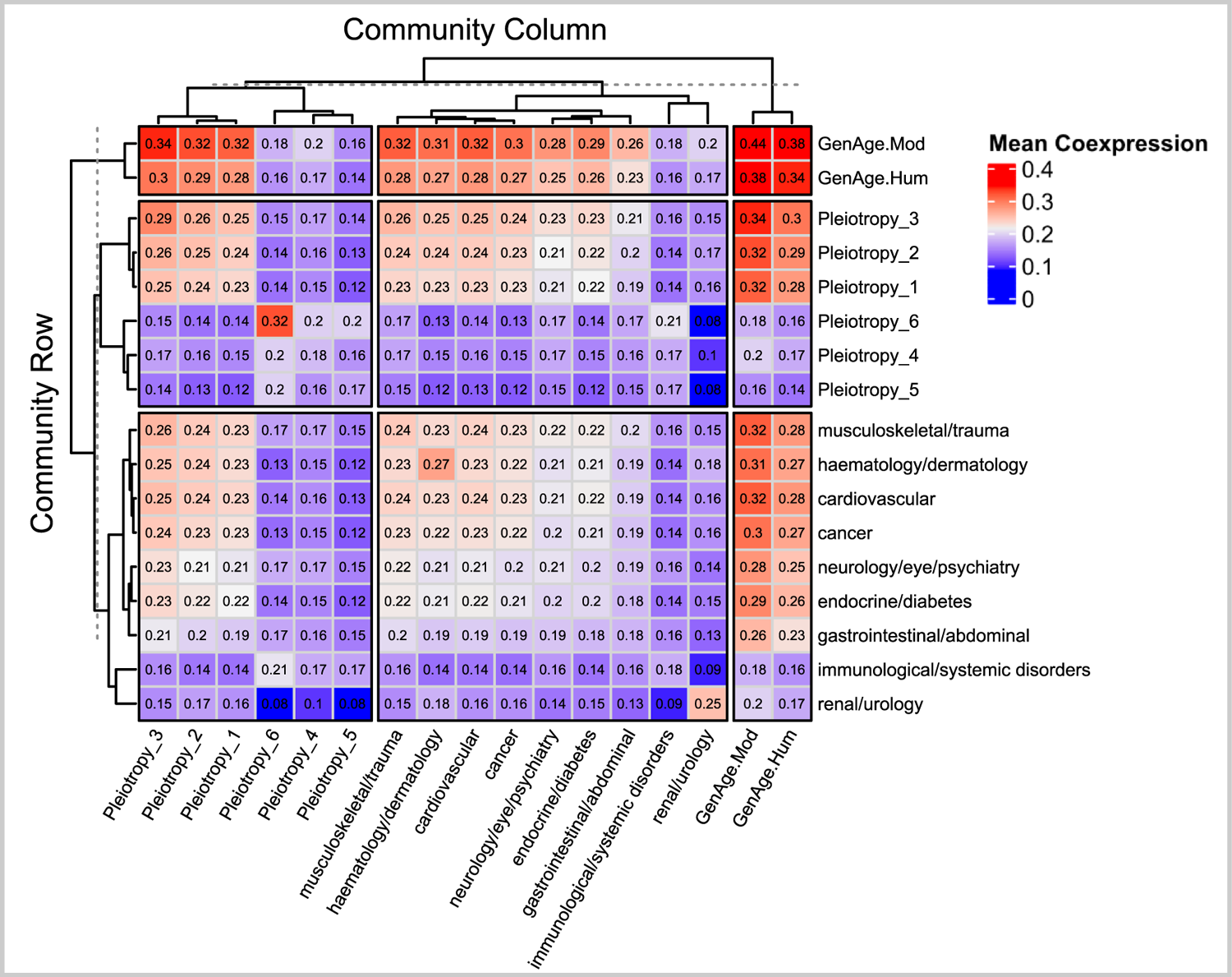
Inter- and intra-community coexpression (*i.e*., self- and cross-coexpression, respectively) among ARCs, GenAge genes, and various levels of *Pleiotropy*. It is organized into three primary groups of rows and columns related to such categories, each encompassing sub-columns related to the corresponding groups. Each cell’s value represents the average co-expression between the row and column communities. Elements along the main diagonal indicate intra-community co-expressions (*i.e*., self-coexpression), as they correlate each community with itself, while elements outside this diagonal represent inter-community coexpressions. The statistical significance of self-coexpression differences can be found in Supplementary Figure S16.

The coexpression profiles tended to be higher in genes with medium to low (1-3) *Pleiotropy*. These genes had higher self-coexpression than *Pleiotropies* four to six and exhibited somewhat higher coexpression with most diseases and both *GenAge* groups. Conversely, these higher-order *Pleiotropy groups*, displayed a low self-coexpression level. Interestingly, some of the lowest values in Figure 4 were linked to immunological and renal diseases, aligning with our previous observation that high *Pleiotropies* are associated with immunological disorders. These high-order *Pleiotropies* did not demonstrate substantial intra- or inter-community co-expression. Even *GenAge*-associated genes, which depicted high coexpression with all groups, coexpressed poorly with high pleiotropy genes.

The full distribution of Intra-community coexpression values across ARCs and *GenAge* groups is depicted in Figure 5A. The *GenAge* groups and *Immunological systemic disorders* encompass two extremes of the ARCs-defined Intra-community coexpression spectrum, with *GenAge* groups, and especially *GenAge*_*Mod*_, being notably and significantly higher than any ARC, and Immunological systemic disorders at the lowest, although very similar to other ARCs and without a significant difference to most ARCs but still significantly lower than the *GenAge* groups.

**Figure 5.**
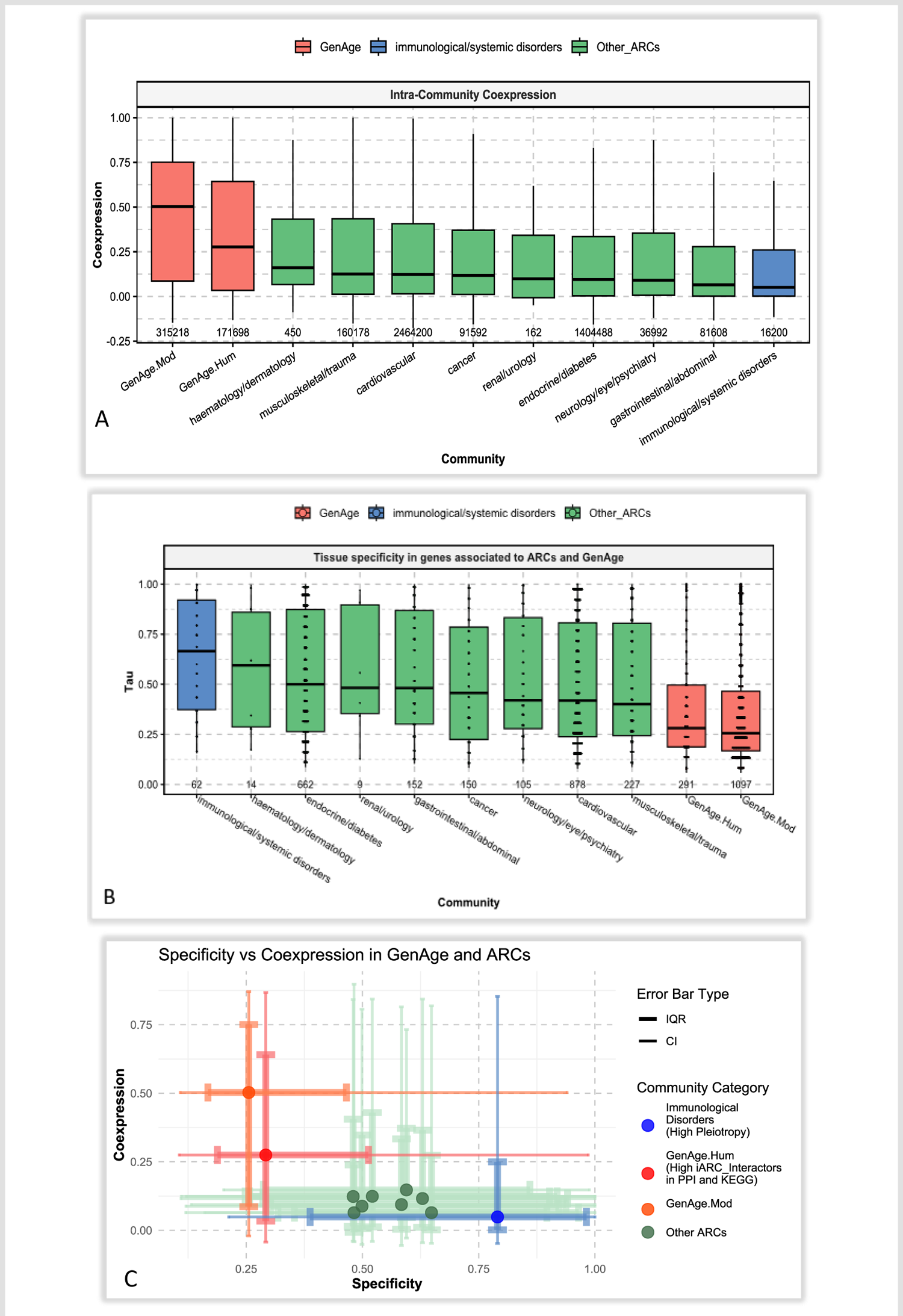
Intra-Coexpression and Tissue Specificity across ARCs and Ageing. **A**. Tissue Specificity. **B**. Coexpression. **C**. Tissue Specificity vs Self-Coexpression. Statistical tests for group differences are depicted in Supplementary Figures S16 and S17. *GenAge* groups are always significantly different from the other groups, including Immunological systemic disorders. On the other hand, immunological systemic disorders are not always significantly different from other communities in terms of Specificity. However, it tends to have a lower mean in terms of self-coexpression.

### Tissue Specificity and Coexpression

We also explored the relationship between *Pleiotropy*, different ARCs, and ageing-related genes (which in turn implies high *iARC-Interactors* on PPI and KEGG) in the context of tissue-specificity, as well as the relationship of this measure with coexpression. Consequently, we sought the Tau coefficient from the Human Ageing Genomic Resources (HAGR) database (Palmer *et al*., 2021). The Tau coefficient measures the degree of tissue-specificity for a gene’s expression based on the GTEx database (GTEx Consortium, 2013). For Tau, a value of “0” indicates a homogeneous expression across tissues, while a value of “1” indicates expression focused on specific tissues.

The distribution of Tau values across ARCs and the two *GenAge* groups is depicted in Figure 5B. Once more, the *GenAge* groups and Immunological systemic disorders encompassed two extremes of the spectrum, although in an opposite fashion than with coexpression, as here the *GenAge* groups were notably and significantly lower than any ARC, whereas Immunological systemic disorders was at the top but without significant differences with other communities excepting *GenAge* groups (Significance measures depicted in Supplementary Figure S17). Figure 5C depicts a Self-coexpression vs Specificity plot, which highlights the opposite polarities of the *GenAge* groups and Immunological systemic disorders. *GenAge*_*Hum*_, which was significantly associated with moderately to high *iARC_Interactors* in the *ARC.PPI* and *ARC.KEGG* networks (Figure 3), depicted an upper-left extreme, isolated from other ARCs by a significantly higher yet still moderate self-coexpression and significantly lower tissue specificity. *GenAge*_*Mod*_ showed even higher self-coexpression, by far, and lower tissue specificity than *GenAge*_*Hum*_. Immunological systemic disorders, which are associated to high *Pleiotropy*, accomplished the opposite role, with a lower-right extreme defined by the lowest self-coexpression (although not always significantly, Supplementary Figure S16) and the highest Tissue Specificity (significant with respect to the *GenAge* groups but not with respect to other ARCs, Supplementary Figure S17). Note how the remaining ARCs tended not to follow a linear patter across this plot.

### Tau index distribution across *iARC_interactors* and Pleiotropic genes

Figure 6 presents box plots of the distribution of Tau Scores across diverse ranges of *iARC-Interactions* and *Pleiotropies.* The *ARC.PPI* network displays strong significant differences in tissue specificity across all categories of *iARC-Interactors*, except for the minor but still significant difference between *iARC-Interaction_1* and *iARC-Interaction_4+*. Tao values decrease subtly and sequentially as *iARC_Interactions* decrease, ranging from a mean of 0.27 in *iARC-Interaction_1* to 0.20 in *iARC-Interaction_4+*. The number of associated genes also decreases as the level of *iARC_Interactions* increases. Moreover, the distributions of Tao values display median values that lie relatively at the middle of the interquartile range for all *iARC-Interaction* groups.

**Figure 6.**
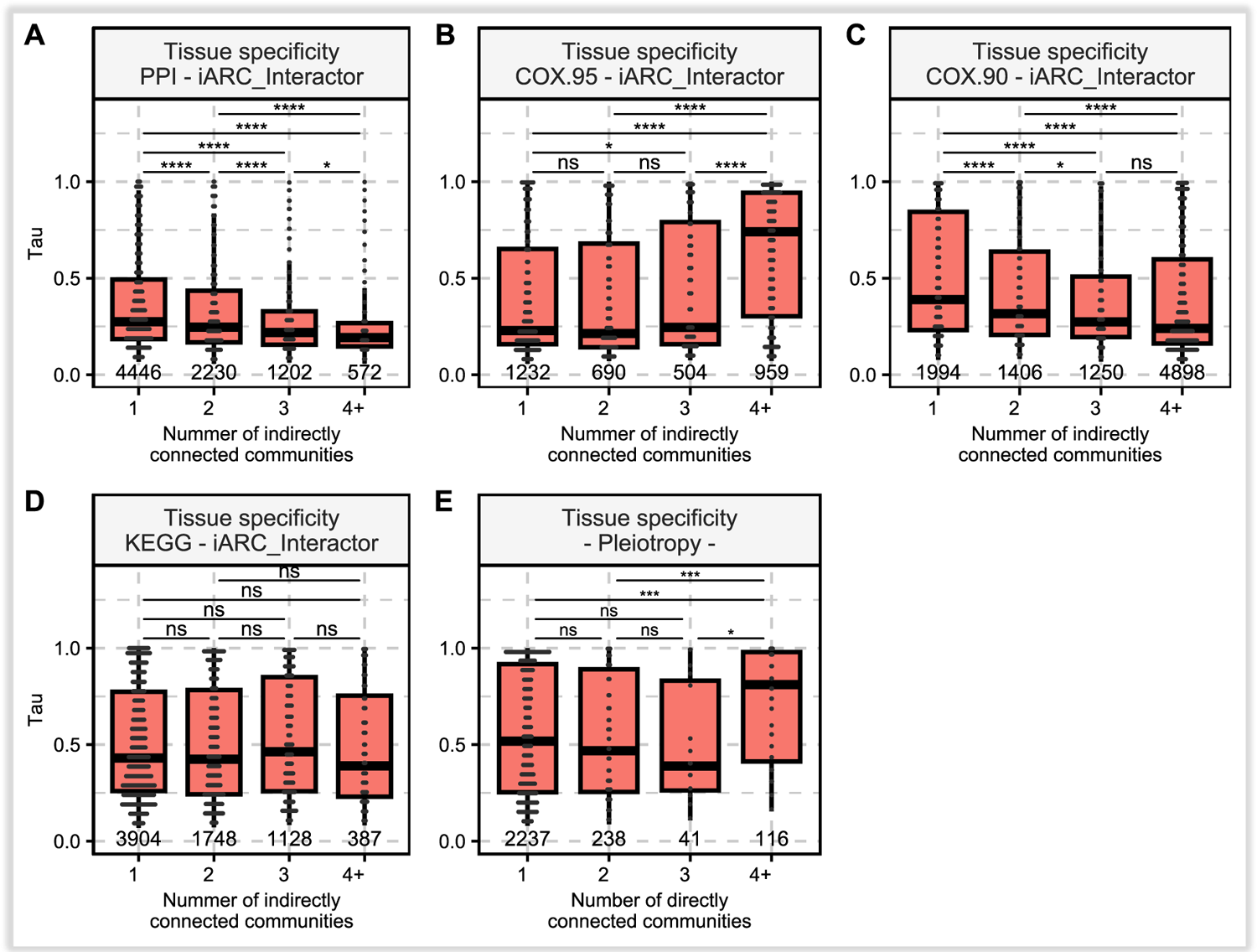
Tau score vs and *iARC interactions* and *Pleiotropy*. The X-axis represents the number of ARCs a gene can reach, either indirectly (*iARC_Interation* corresponding to Figures A-F) or directly (*Pleiotropy,* corresponding to Figure E). The Y-axis denotes the corresponding Tau value, which reflects tissue specificity. Specifically, the bar presented at the X-value “1” in each graph depicts the distribution of Tau values for the set of genes that are associated with only one ARC. The number “2” similarly represents genes associated with two ARCs, “3” for those associated with three communities, and the “4+” symbolizes the Tau values for genes that are directly or indirectly associated with four or more ARCs. **A**. *ARC.PPI* network. **B.** *ARC.COX*_95_ network. **C.** *ARC.COX*_90_ network. **D**. *ARC.KEGG* network. **E**. Tau vs Pleiotropy across all the ARDs genes. Statistical differences were computed using Wilcoxon Signed-Rank and adjusted for multiple tests with Bonferroni.

The *ARC.PPI* network displays strong significant differences in tissue specificity across all categories of *iARC-Interactors*, except for the minor but still significant difference between *iARC-Interaction_1* and *iARC-Interaction_4+*. Tao values decrease subtly and sequentially as *iARC_Interactions* decreases, ranging from a mean of 0.27 in *iARC-Interaction_1* to 0.20 in *iARC-Interaction_4+*. The number of associated genes also decreases as the level of *iARC_Interactions* increases. Moreover, the distributions of Tao values display median values that lie relatively at the middle of the interquartile range for all *iARC-Interaction* groups.

*ARC.COX*_90_ has more genes than *ARC.COX*_95_, but both display a trend of decreasing gene counts until *iARC-Interaction_3* and then a significant rise at *iARC-Interaction_4+*, more noticeable in *ARC.COX*_90_. The Tao mean values differ between the two. For *ARC.COX*_90_, Tao values decrease from 0.4 at *iARC-Interaction_1* to 0.25 at *iARC-Interaction_4+*. Meanwhile, *ARC.COX*_95_ has lower initial Tao values, with a sharp rise to 0.74 at *iARC-Interaction_4+*. Significant Tao value differences exist in *ARC.COX*_90_ except between *iARC-Interaction_2-3* and *iARC-Interaction_3-4+.ARC.COX*_95_ lacks *o*nly significant differences between *iARC-Interaction_1-2* and *iARC-Interaction_2-3*. Gene distributions in *ARC.COX*_90_ tend towards lower means, while *ARC.COX*_95_ leans towards lower medians, except *iARC-Interaction_4+* which breaks the descending pattern to go far higher (∼0.75). *ARC.KEGG*, exhibited a decline in gene counts as *iARC-Interaction* levels rose. However, the Tao values across groups did not show any significant difference nor a trend of specificity with increasing *iARC_Interactors*. The specificity values remained contained between the values of 0.4 and 0.48.

Gene counts decrease sharply with increasing levels of *Pleiotropy*, ranging from 2,237 genes in *Pleiotropy_1* to just 116 in *Pleiotropy_4+*. Tao values present a more complex pattern: slightly decrease non-significantly from 0.51 in Pleiotropy_1 to 0.48 in Pleiotropy_2 and 0.4 in *Pleiotropy_3*. However, there is a significant jump in Tau values at higher levels of *Pleiotropy*, specifically in the *Pleiotropy_4+* group, where the mean Tau value escalates to 0.8. This increase is statistically significant compared to previous pleiotropic groups, albeit at varying significance levels. Furthermore, the distribution of tau values varies across the groups—centered for *Pleiotropy_1* and *Pleiotropy_*2, skewed low for *Pleiotropy_3*, and skewed high for *Pleiotropy_4+*.

### ARC-related gene expression across tissues

Figure 7 shows gene expression levels in log-transformed reads per kilobase million (RPKM) (Palmer *et al.,* 2021) for immunological disorders- and ageing-related genes across different tissues. Immunological disorder-related genes typically have lower median expression levels in all tissues, with notable variance and lower mean expression in blood, despite their importance in immunity. This pattern is more pronounced than in other tissues, such as the pancreas, kidney, liver, and heart. The GTEx database lacks data on specialized immune tissues like bone marrow and thymus. Ageing-related genes, in contrast, exhibit more consistent and higher average expression levels across all tissues. However, the pancreas, liver, kidney, heart, and blood generally show the lowest expression for these genes. Supplementary Figure S18 displays the tissue expression plot for all the ARCs.

**Figure 7.**
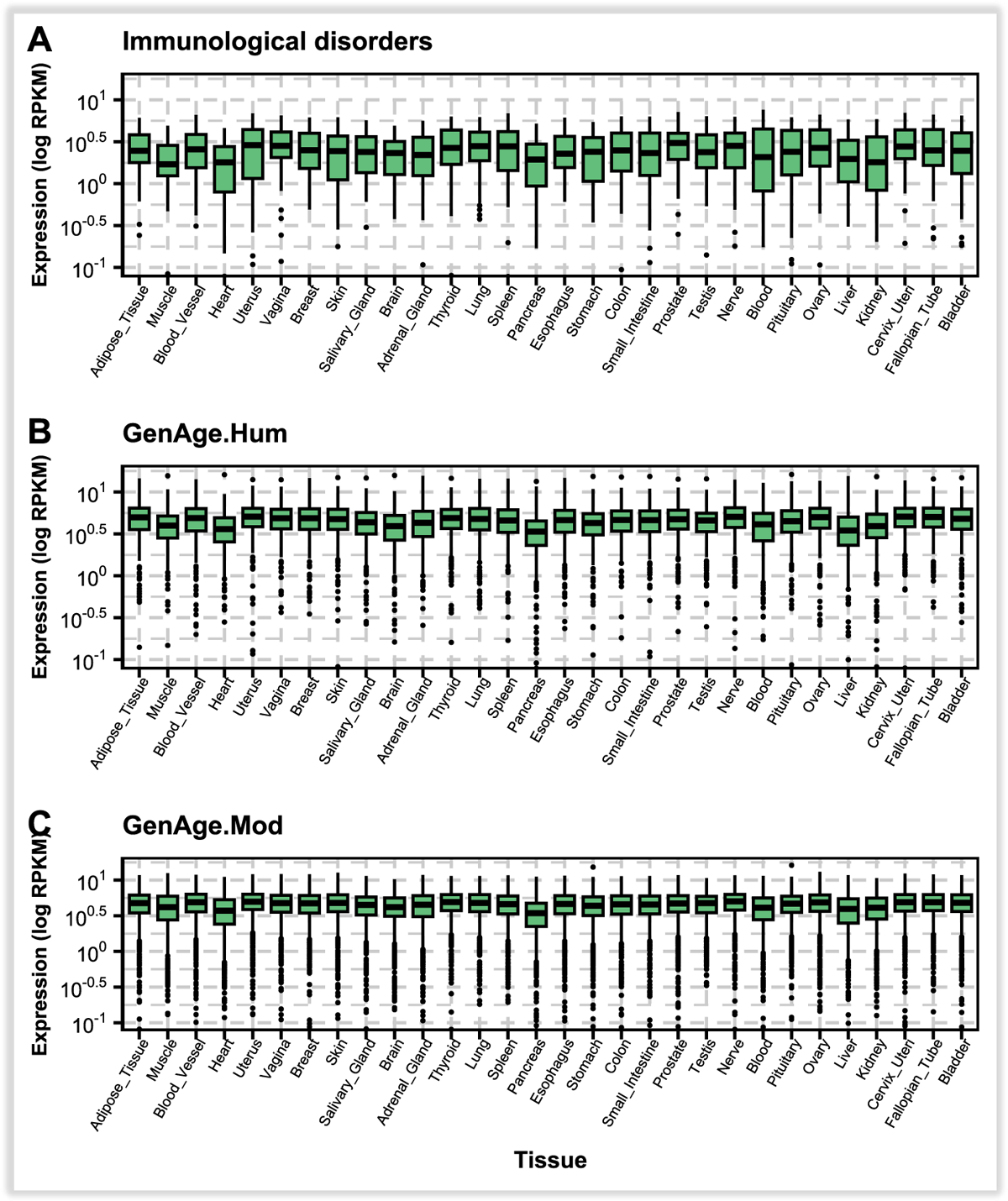
Gene expression levels across various tissues in the GTEx database for genes associated with Immunological disorders- and ageing-related genes. Expression levels (Reads per million) are shown on a logarithmic scale. Tissues commonly associated with each ARC were highlighted in red. **A)** Immunological systemic disorder-related genes. B) *GenAge*_*Hum*_-associated genes, without associated tissue, as ageing affects the entire body, C) *GenAge*_*Mod*_-associated genes, without associated tissue either.

### Prediction of ageing-related genes and pathways

We predicted ageing-associated genes based on the four previously established networks. Different methods were employed, leveraging the concept of genetic proximity to ARDs and ARCs as features which, through averaging and ML algorithms, facilitated our predictions. In-depth procedures are outlined in “Methods”, while all the performances are detailed in Supplementary Table S5. In short, using the Aurea Under the ROC Curve (AUC) as prediction metric, it was observed that *ARC.PPI* scored the highest (0.80), followed closely by *ARC.KEGG*_*both*_ (0.77). *ARC.COX*_95_ scored 0.70, whereas *ARC.COX*_90_ achieved the poorest performance (0.60).

We focused on the two top-performing networks *ARC.PPI* and *ARC.KEGG* - for gene prediction purposes. Supplementary Tables S6 and S7 display their top candidate ageing-related genes, respectively. Notably, the genes with higher prediction scores predominantly exhibit null *Pleiotropy*, meaning that almost none directly connect to any diseases. However, most of them have a high *iARC-Interaction*, mostly covering more than 4 ARCs and 10 ARDs, with only a few exceptions.

The overall correlation with the ageing-relatedness probability remained intermediate for *iARC-Interactions*, low for *Pleiotropies*; higher for ARDs than for ARCs in *iARC-Interactions*; and higher for *ARC.PPI* than for *ARC.KEGG* in *iARC-Interactions* as well. Namely, correlations with ageing-relatedness in *ARC.PPI were 0.42 (iARC-Interaction), 0.46 (iARDs-Interaction), −0.03 (ARCs-Pleiotropy) and −0.04 (ARDs-Pleiotropy);* whereas in *ARC.KEGG* results were 0.33 (i*ARCs-Interaction*), 0.40 (i*ARCs-Interaction*), −0.08 (i*ARCs-Interaction*) and −0.07 (*iARCs-Interaction*). Lastly, it is notable that the two networks do not present any genetic overlap at their top 10 candidate predictions, they both have a relatively low ageing-relatedness probability correlation (0.25, Supplementary Figure S26).

The top 5 non-GenAge genes most likely associated with ageing based on the ARC.PPI network were HSPB1, GNL3, RAF1, KAT5 and UBC; whereas for ARC.KEGG were PIK3CD, MAPK11, MAPK12, PIK3R2, and MAPK10. The top GO terms linked to the top 30 ageing-associated gene candidates from *ARC.PPI* were the *regulation of biosynthetic macromolecular processes, developmental process regulation, cellular localization, cell cycle*, and *response to abiotic cycles*, among others. For *ARC.KEGG*, the most significant terms were *intracellular signal transduction, regulation of macromolecular metabolic processes*, and *phosphorylation*. The number 30 was chosen as it covers less than 1% of genes in both networks while covering enough genes for enrichment analysis. Further information on the top genes and biological processes can be found in Supplementary Tables S6-S8.

## Discussion

Starting with a meta-analysis of a previous work on UK Biobank that identified common genes across ARDs (Dönertaş *et al*., 2021), we found that the immune systemic disorders emerged as a central player, with its associated genes showing extensive interactions across ARCs, displaying high *Pleiotropy*. Although immunosenescence or inflammaging are well-recognized players in ageing (Santoro *et al*., 2021), the identified *Pleiotropic* genes were enriched for both non-strictly (*nucleosome assemb*ly and *urate transport*) and strictly (*innate immune response in mucosa, T cell receptor signalling pathway,* and *antibacterial humoral response*) immune-related biological processes. This supports previous studies suggesting that the relationship between immunological conditions and ageing should be deeply studied (Fabian *et al*., 2021). A notable finding that motivated our further research was the limited overlap between ageing-associated genes from GenAge and ARCs-related genes, with only 10% of ageing-related genes linked to ARCs, and 0% is related to highly *Pleiotropic* genes, suggesting two distinct mechanisms to influence multiple ARCs, one mediated by ageing-related genes and another mediated by common genes across ARCs.

The fact that human ageing-related genes showed a higher number of indirect connections to ARCs than random genes and even than the neighbours of disease-related genes indicates that these genes are not just a random subset of neighbours to ARDs-related genes, but the subset of these neighbours that more extensively tends to connect with these ARCs. Homologs of ageing-related genes in model organisms also showed more connections than their neighbours, but less markedly than human ageing-related genes, suggesting that, while some genetic foundations of ageing-related genes are conserved, the specific protein and metabolic connections within the human context seem to influence ARDs stronger. However, in co-expression networks, ageing-related genes, both human and from model organisms, do not show significant differences from neighbouring disease genes, suggesting their influence on disease sets is more diluted regarding co-expression at high thresholds than in physical or metabolic interactions. This could align with recent studies showing mismatches between protein levels and gene expression in ageing tissues (Keele *et al*., 2023).

We further analysed ARCs in terms of tissue specificity and coexpression, finding opposite behaviours between highly *Pleiotropic* and highly *iARC_interactive* (*ARC.PPI* and *ARC.KEGG*) genes, which by extension associate with immunological disorders- and human ageing-related genes, respectively. Immunological disorders-related genes stood out for their high tissue specificity but were among the lowest in self- and cross-coexpression with most ARCs. In contrast, ageing-related genes, especially those associated with model organisms, displayed minimal tissue specificity but the highest levels of self- and cross-coexpression with many ARCs. The lower specificity of ageing-related genes was partially consistent with the trend that genes with the highest number of indirectly connected ARCs through PPI tended to have lower tissue specificity. The reason we mention “partially” is based upon the fact that, while human ageing-related genes tended to have some of the highest number of indirectly connected ARCs and also a low tissue-specificity, ageing-related genes in model organisms were even less tissue-specific; however, they were not as connective in terms of reachable ARCs through PPI and KEGG interactions, although they remained just slightly less connected than ARC-neighbouring genes.

It may seem paradoxical that highly *Pleiotropic* genes, such as those of the immune system disorders, whose GO terms also encompass *nucleosome assembly*; exhibited high tissue specificity and low self- and cross-coexpression given their broad impact on ARCs (Liu *et al*., 2012). A potential explanation could be that the immune system tends to have its own expression profile, but it operates ubiquitously across the body, responding to threats in all tissues, despite each tissue potentially requiring different immune gene-related needs (Domínguez *et al*., 2022; Hao *et al*., 2022). In this sense, an immunological-related tissue such as the thymus or bone marrow was absent in the GTEx-based tissue specificity dataset of HAGR (Palmer *et al*., 2021). Still, it was observed that the blood tissue (which contains, in part, immunological cells) presented a slightly lower overall expression of immune disorder-related genes than other tissues. Similarly, each tissue has distinct epigenetic requirements, influencing the expression of *nucleosome assembly* genes. In this sense, chromatin organization is also associated with inflammaging (Zhang *et al*., 2022). Ageing-related genes, on the other hand, seemed to exert their influence over multiple ARCs in a more generalized way, as low specificity is indicative of ubiquitous roles across various biological pathways (Kitsak *et al*., 2016), which could be consistent with its highest self-and inter-coexpression, potentially indicating a constant yet subtle interplay with most ARCs.

The relationship between ageing-associated genes and ARDs appears to have been most accurately characterized through PPI. This network facilitated the optimal prediction of ageing-associated genes, based primarily on their proximity to genes linked to these diseases. It is important to note that multiple outcomes of this network exhibit bias to diverse degrees, attributed to the extensive research on common ageing-related genes (Gillis *et al*., 2014). The KEGG-based metabolic interaction network also revealed a robust connection between ageing and ARDs-associated genes, albeit this association was slightly weaker. The predicted ageing-related genes at the PPI network tended to associate with cancer-related functions. This trend could reflect the network’s inclination towards well-researched genes like cancer-related genes. However, another explanation could point to an antagonist *Pleiotropy*-like relationship between cancer and ageing, where overactivation of genes like p53 could accelerate ageing, while a deficiency can lead to cancer (Campisi, 2005; Ungewitter & Scrable, 2009). In contrast, the top genes in the KEGG network were notably aligned with developmental processes and cellular reactions to chemical stimuli, complementing previous findings on KEGG pathways and ageing (Ukraintseva *et al*., 2021; Fabris & Freitas, 2016). Notably, even as the PPI and KEGG networks depicted good predictive capabilities, they highlighted different top genes, suggesting different mechanisms linking ageing-related genes with ARCs.

We noted two distinct outcomes at the level of co-expression. Defining co-expression by a threshold and considering only genes expressing above 95 or 90%, we found that ageing-associated genes exhibited no distinction. At first sight, this could give the impression that coexpression is not a means by which ageing-related genes interact with diseases. However, it was noteworthy that ageing-related genes are distinct in terms of expression when measured in smaller quantities, such as between 30 to 45%. This suggests that ageing-related genes do not necessarily affect diseases through processes that require huge co-expression with them, but instead, a more modest (yet high relative to other groups of genes) and possibly cumulative association, which is not observable under high co-expression thresholds. This was particularly true for homologs from model organisms, which express more significantly among themselves and with other diseases compared to human ageing-related genes, suggesting that the conserved component across species of model organism homologs could possibly undertake essential ARDs-related functions across cells and thereby manifesting moderate co-expression with genes associated to multiple ARCs at various tissues. Insights on this topic have been previously discussed, albeit for human genes (Yang *et al*., 2017).

## Conclusions

The research highlights two main types of genes connecting multiple ARCs: 1) genes with high *Pleiotropy, i.e.*, GWAS-associated with multiple ARC, with immunological systemic disorders being a common factor; 2) *iARC_Interactors*, i.e., genes that are indirectly associated with multiple ARCs through gene interaction, albeit without the need of a GWAS association. In the KEGG, but especially the PPI context, such *iARC_Interactors* were mostly related to human ageing-related genes. In this direction, it seemed to be an association trade-off of the *Pleiotropy* and *iARC_Interactors* groups, as most highly *Pleiotropic* genes tended to display low *iARC_Interactions* and vice versa in the PPI and KEGG networks but not the coexpression ones.

Although there was an overlap between human ageing-associated genes and those in models, human genes have a closer and more specific association with ARCs, particularly in PPI and KEGG pathways. Nevertheless, genes in model organisms exhibited higher coexpression with ARDs-related genes and lower tissue specificity, suggesting a role of gene conservation in the relationship between ARDs and ageing.

Immunological disorder-related genes exhibited high tissue specificity but low coexpression with ARCs, indicating specialized yet limited broader influence. In contrast, ageing-related genes, particularly in model organisms, displayed minimal tissue specificity and high coexpression across various ARCs, suggesting their widespread yet subtle influence on the ageing process. When analysing the tissue specificity tissue-wise, ageing-related genes exhibited a notably low expression variance compared to other ARC-related genes, including those of immunological disorders.

ML approaches, particularly those focusing on gene relationships to ARDs, consistently outperformed ARC-based methods in terms of predicting ageing-related genes, as they provide more information. Simple metrics, like the shortest distance to disease-associated genes, were more effective than complex average distance calculations. A notable finding was that a minimal distance of two intermediate interactions sufficed for accurate predictions, with the *ARC.PPI* network being the most effective for identifying human ageing-related genes, followed closely by the *ARC.KEGG* network.

## Methods

### Selection of ARDs and ageing-related diseases

#### Selection of Non-cancer ARDs

We retrieved GWAS summary statistics of non-cancer self-reported ARDs from a previous UK Biobank study (Donertas *et al*., 2021). Briefly, they extracted age-of-onset profiles of 116 diseases and clustered them into four categories based on their age. The first and second categories involved diseases that increased in frequency with age. Within these, one set of diseases (N=25) demonstrated exponential growth with age, while the other (N=51) exhibited a linear growth. The third category encompassed diseases with no regular pattern; thus, they did not appear at a fixed age but rather manifested randomly. The fourth category included diseases that emerged relatively early, around the age of thirty. For our research, we focused only on diseases with significant associated genes and displayed linear or exponential growth with age, which we refer to as non-cancer ARDs.

#### Selection of Cancer ARDs

90 self-reported cancer diseases were retrieved from UK Biobank (Application 6342) using the field 20001 and all available visits per participant. We then selected those with an occurrence of at least 2000 cases, leaving 11 Cancers to analyse. The age-of-onset of these 11 cancers was then determined using the field 20007 (Interpolated age of participant when cancer occurred). For each cancer, we divided the number of cases at a specific age by the total number of cases to get a distribution of each cancer’s occurrence throughout age. We then used a K-means clustering algorithm to group the cancers according to their age of onset patterns. Like Donertas (Donertas *et al*., 2021), there were exponential- and linear-like age-of-onset increasing trends - 2 and 6 cancers, respectively - and a third trend in which three cancers developed at middle age (Supplementary Figure S1). We then filtered for gender-independent cancers by removing those whose occurrences are biased by 90% or more to one single gender. Out of these diseases, we took the 8 cancers at the increasing age-of-onset clusters for further analysis as cancer ARDs. The resulting cancers were malignant melanoma, non-melanoma skin cancer, skin cancer, and colorectal cancer.

#### Extraction of ageing-related genes

We extracted ageing-related genes from the GenAge database (Human genes): Build 20 (09/02/2020) with 307 genes for human ageing (Tacutu *et al*., 2018). The genes from model organisms were retrieved and mapped to human homologs using the OMA database (Altenhoff, *et al*., 2015), leading to 1147 human homologues.

### GWAS analysis

The GWAS procedures presented in this section are similar to those presented by Donertas (Donertas *et al*., 2021). The main difference is that we used *Plink2* (Chang *et al*., 2015) to perform the preprocessing and the GWAS analysis. Details are provided in the next paragraphs.

### Sample Quality Control

We removed all samples from individuals who withdrew their data from the UK Biobank. Samples without genotypes were excluded, resulting in the removal of 14,248 samples. We then filtered discordant sex data by cross-referencing ‘31-0.0’ (Self-reported Sex) and ‘22001-0.0’ (Genetic Sex), leading to the exclusion of 378 samples (0.077%) with discrepancies. Furthermore, samples with sex chromosome aneuploidy (identified using ‘22019-0.0’) were excluded, encompassing 651 cases or 0.133% of the dataset.

### Heterozygosity

Genotype call rate and heterozygosity, indicators of DNA sample quality, were scrutinized using UK Biobank’s suggested exclusions. We excluded cases with ‘poor heterozygosity/missingness’ ( 6 cases, field ‘ ‘), those declaring mixed ancestry (6 cases, field ‘ 8 ‘), and those with high heterozygosity or missing rate after ancestry correction (8 cases). Additional 968 outlier cases for heterozygosity or missing rate were identified (field ‘ -. ‘).

Based on these criteria, 3,697 samples were removed. However, the specific numbers mentioned earlier might not directly add up to this total due to overlaps, where some samples met more than one exclusion criterion. To this point, 484,598 samples passed the QC.

### GWAS execution

For each disease, we performed GWAS using *plink2* (Chang *et al*., 2015). Specifically, we computed the UK Biobank’s imputed genotypic data in BGEN format. We removed participants who did not meet our QC standards and data from individuals who withdrew from the UK Biobank. SNPs deviating from Hardy-Weinberg equilibrium or with a minor allele frequency below 0.01 were excluded. Our covariates encompassed Sex, Age, BMI (derived from weight and height), assessment center, ethnicity, batch, number of self-reported non-cancer diseases and cancers; the minimum, maximum, and average ages of parental deaths, marking unavailable or inapplicable data as NA, the participant age of death (setting unavailable data as NA as well); and the first 20 PCs. Our results excluded the MHC region (chr6: 28,477,797 - 33,448,354), and positions with a p-value lower than 5×10^-8^ were identified as significantly associated (Supplementary Figure S2). We corrected for multiple tests using Bonferroni.

### SNP to Gene Mapping

We used the *biomaRt* package (Durinck *et al*., 2009) to retrieve the genomic ranges of protein-coding genes. We then retrieved the disease-associated genes for each cancer by mapping genes to significant SNPs if the SNP is either within the genomic range corresponding to the gene or as far as kb from the gene’s genomic region. We computed this using the R package *GenomicRanges* (Lawrence *et al*., 2013).

### Creation of the Gene-ARD and Gene-ARC relationships

The previously created SNP-to-gene mapping throughout the ARC-related GWAS allowed us to establish a relationship between each ARD and their respective genes. This resulted in 2588 statistically significant genes associated with at least one ARD, covering 58 diseases. Each of these relationships is what we call Gene-ARD association.

The UK Biobank organizes self-reported diseases, both cancerous and non-cancerous, hierarchically. There are main disease categories, each containing more diseases organized in subcategories associated with the main category, *e.g*., cardiovascular, haematological, etc. We used this hierarchical structure to group diseases according to the main category to which they belong. For instance, ARDs such as hypertension, heart attack, arrhythmia, etc., can be classified as cardiovascular, thus forming what we refer to as ageing-related disease communities (ARCs), as shown in Supplementary Figure S3. We established Gene-ARC associations by labelling all the genes associated with at least one ARD of the ARC as ARC-associated.

In this paper, we worked mainly with Gene-ARC associations as they provide more phenotypic heterogeneity than ARDs. This occurs because some ARDs within the same ARC may be closely related under the hierarchy, resulting in similar phenotypic and genetic expression. ARCs would not be as redundant and provide more tissue-wise insights. Nevertheless, the Gene-ARDs approach is still used for prediction purposes in the “prediction of ageing-related genes” section.

### ARC networks

We expanded the Gen-ARC relationships by integrating gene-gene connections from three databases: BioGRID for PPI (Stark *et al*., 2006), GeneFriends for gene co-expression (Raina *et al*., 2023), and KEGG Pathways (Kanehisa *et al*., 2016).

#### *ARC.PPI* network

Human PPI data was retrieved from BioGRID (Stark *et al*., 2006) – physical interactions version 4.4.221 - and used to create an unweighted network with R’s *igraph* package. The PPI network contained 10,365 nodes (genes) and 62,640 edges (gene interactions). We then created the *ARC.PPI* network upon mapping the GWAS-associated ARCs-related genes, as well as ageing-related genes, into the PPI network. We only allowed PPI-connected genes into this network, thus neglecting all the non-PPI-overlapping ARC- and ageing-related genes.

#### *ARC.COX* networks

Gene coexpression data was obtained from GeneFriends (Raina *et al*., 2023) – version 5. The dataset is a square matrix with 44,947 rows and columns representing genes, and cells are coexpression levels. The genes are coded using Ensemble; we first converted them to HGNC using *BiomRt,* leading to *32,915* genes with codes for both Ensemble and HGNC. Two binary matrices were then generated based on coexpression thresholds of 90% and 95%. If a cell is above 90% or 95%, according to each matrix, then it takes the value “ 1”; otherwise, it takes “0”. Taking the names of the genes at the rows and columns whose intersecting cell scores “1”, we created two two-column data frames (one for each coexpression threshold), with the first column containing the rows and the second the columns of the main matrix at which the value of “1” occurred. The *igraph* (Csardi *et al*., 2023) library was then used to create two coexpression networks, we called them *COX*_90_and *COX*_95_, respectively. The assignation of ARC- and ageing-related genes occurred similarly to *ARC.PPI* but in the context of these networks.

#### *ARC.KEGG* networks

Gene associations from KEGG pathways were retrieved using the *KEGGlinks* library (White & Medvedovic, 2016) to extract pathway names from the KEGG database. We then employed the *KEGGgraph* library (Zhang & Wiemann, 2009), which, creates data frames of gene interactions within each pathway through KEGG Markup Language representations resulting in 250 queries for human-associated pathways. We row-binded these dataframes into one, removing duplicated rows (i.e., duplicated gene interactions) We then used igraph to convert the resulting dataframe into an undirected graph, resulting in the KEGG network with 6,407 nodes and 62,007, interactions. The assignation of ARC- and *GenAge*-related genes occurs similarly to *ARC.PPI*. These new networks include genes outside ageing- and ARC-related genes to cover all the genes in each network. Table 1 provides further insights on this subject.

**Table 1.** Measures of genes and phenotypes nodes and edges at extended *ARD* networks.

| Network | ARDs genes | Gene-Gene Network genes | Disease genes<br>∪<br>Gene-Gene Network | Disease genes<br>∩<br>Gene-Gene Network genes | Gene-Gene edges | Gene-ARDs edges | ARDs Connected to Gene-Gene Network (Out of 58) | ARCs Connected to Gene-Gene Network (Out of 9) |
| --- | --- | --- | --- | --- | --- | --- | --- | --- |
| *ARC.PPI* | 2698 | 10,586 | 12,332 | 962 | 62,092 | 5308 | 51 | 9 |
| *ARC.COX<sub>90</sub>* | 2698 | 11,136 | 12,946 | 888 | 1,463,273 | 5308 | 49 | 9 |
| *ARC.COX<sub>95</sub>* | 2698 | 5,022 | 7,295 | 425 | 137,994 | 5308 | 45 | 9 |
| *ARC.KEGG* | 2698 | 6,407 | 8,407 | 608 | 62,007 | 5308 | 50 | 9 |

### Pleiotropy and Indirect Interactor definitions

*In the context of our work, Pleiotropy* measures the number of phenotypes that a gene can influence through GWAS-association. Indirect interactions measure how a gene influences multiple phenotypes through interaction with its GWAS-associated genes. They both can also be ARD- or ARC-related. Therefore, four permutations are possible. Figure 8 illustrates the concept. In this work, ARD-*Pleiotropy* refers to the number of ARDs that a gene is GWAS-associated with. *ARD-Pleiotropy* refers to the number of ARDs with at least one GWAS-associated gene interacting with the gene of interest through the Gene-Gene network. *iARD-Interaction* (where the “i” comes from indirect) indicates the number of *ARDs* the gene indirectly interacts with. ARC-*Pleiotropy* and *iARC-Interaction* apply similarly but for ARCs. Since this work focuses mostly on ARCs, we use the term *Pleiotropy* as a synonym of ARC_*Pleiotropy*.

**Figure 8.**
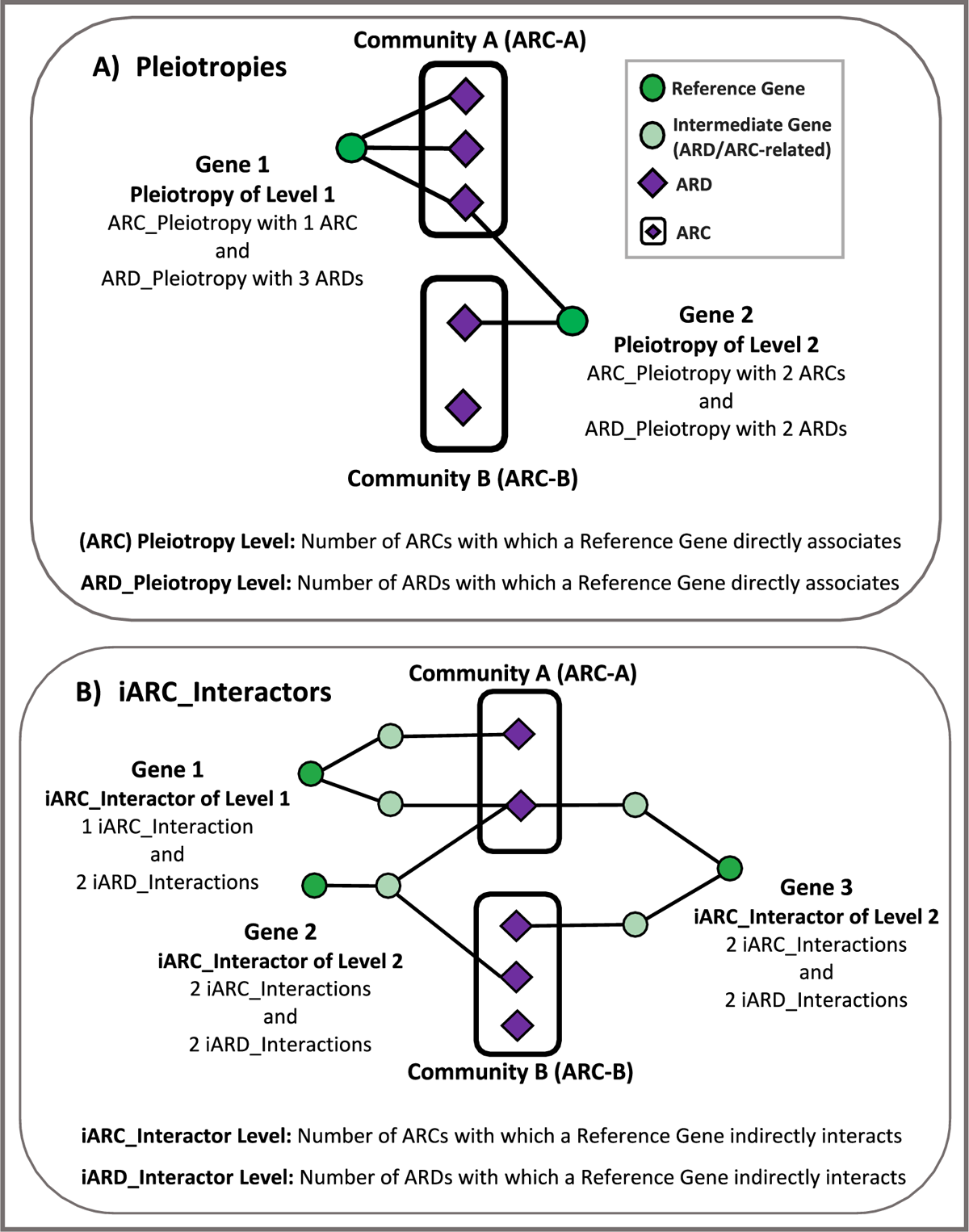
*Pleiotropy* and *iARC_Interactors*. A single gene can influence multiple ARDs or ARCs through either a direct association or indirect interaction. The intense green circle represents the reference gene, lighter green circles indicate intermediate genes, and purple diamonds symbolize diseases or phenotypes. **A)** *Pleiotropy*. The reference gene directly associates with different ARDs which in turn associates the genes with the ARCs to which the ARDs belong. *Pleiotropy* is categorized as *ARD_Pleiotropy* or *ARC_Pleiotropy,* and a level value is assigned depending on the number of associated ARDs or ARCs, respectively. Gene 1 indicates that a Gene associated with multiple ARDs does not necessarily associates with multiple ARCs. Gene 2 indicates association with 2 ARCs through association with 2 ARDs. If no context is provided, we refer to *ARC_Pleiotropy* simply as *Pleiotropy* throughout this work. **B)** *iARC_Interactor*. The reference gene indirectly interacts with different ARDs, which indirectly associates the genes with the ARCs to which the ARDs belong. The letter “i” accounts for “indirect”. The indirect interaction is mediated through Gene-Gene interaction with an ARD/ARC-related gene. Interactors are categorized as i*ARD_Interactor* or i*ARC_Interactor,* and a level value is assigned depending on the number of indirectly associated ARDs or ARCs, respectively. Gene 1 indirectly interacts with 1 ARC despite being associated with 2 ARDs. Gene 2 interacts with a *Pleiotropic* Gene of Level 2, leading to indirect interactions with 2 ARDs and ARCs.Gene 3 indirectly interacts with 2 ARDs in two different ARCs. Node designs were inspired by (Weighill *et al*., 2019).

Note that a gene interacting with multiple ARDs does not necessarily interact with different ARCs. Gene 1 in Figure 7A illustrates this point as it interacts with three ARDs belonging to the same ARC. Conversely, Gene 2 at the same sub-figure interacts with only two ARDs, each belonging to different ARCs, displaying ARC-*Pleiotropy*. It’s inferred that while ARD *Pleiotropy* does not guarantee ARC *Pleiotropy*, the latter requires the former to exist. The same logic is applied to indirect interactions (Figure 7B). Additionally, a gene can be both *Pleiotropic* and *iARC-Interactive*.

### Indirect Interactor analysis

For each gene, we computed the number of *iARC-Interactions*. This is the number of ARCs whose genes present at least one interaction with the Reference gene. Given this score, we grouped the genes by *GenAge, Diseases, Neighburs* and *Others*. Statistics were computed using the Mann-Whitney U Test since the distribution was strongly biased towards low *iARC-Interactions*, following a decreasing exponential shape.

### Coexpression

For coexpression analysis, we used the co-expression matrix for humans at GeneFriends (44,947 row and columns). We mapped all the Ensemble code values in the rows and columns to HGCN using *biomaRt* (Durinck *et al*., 2009).

#### Intra-community coexpression

For intra-community co-expression, we selected the subset of the coexpression matrix whose rows and columns correspond to the ARC-associated genes available within the GeneFriends dataset. This resulted in a square and symmetrical matrix whose lower triangular and main diagonal were transformed into a numeric array for statistical analysis where error ranges, mean, and statistical variances for each group were computed.

#### Inter-community co-expression

This analysis was conducted by selecting the genes associated with the first group as the coexpression matrix rows and those linked to in the second group as the matrix’s columns, resulting in a rectangular submatrix. Subsequently, this submatrix was transformed into a numeric array for analysis. Statistical testing was carried out using a t-test.

### Tissue Specificity

Tissue specificity was derived from Tau values in the HAGR database (Palmer *et al*., 2021). The downloaded file provides 39,366 genes across rows, 30 GTEx tissues at columns, and expression levels at cells measured by RPKM. There is another column with Tao values for gene expression specificity. A Tao value of “1” means that the gene is primarily expressed in specific tissues, whereas a value of “0” indicates a tendency towards more homogeneous expression levels. Tissue specificity differences between groups were tested using the Mann-Whitney U Test, given that the distribution of Tau values tended to display a bimodal distribution at low and high Tau values.

### Prediction of ageing-related genes

We predicted potential ageing-related genes by assessing their association with either ARCs or ARDs using algorithms based on network topology, estimating further biological insights from the inference process.

We define distance as the number of edges that must be transited from a Reference Gene to reach a Target Gene following the shortest path. In this context, we created a Gene-Gene Distance matrix using the “distances” command of the *igraph* library, where we consider genes in the rows to be the reference genes and genes in the columns Target Genes. When no genetic path between the two genes exists, the Distance is considered Infinite. If a Gene is itself the Reference and Target Gene, then the distance is zero. In the context of a specific ARC, we define Distance as the shortest path connecting the Reference Gene to the closest gene associated with such Target ARC (see Figure 9).

Some of our algorithms require computing the average distance to ARCs. However, this is impossible if a Reference Gene is totally unconnected to at least one ARC (*i.e*., lacking paths to such ARC genes) because the Distance would turn infinite for such ARC, disrupting calculations when averaging Distances to all ARCs. To cup with this, we mapped the Distance to a normalized, inversely proportional, measure that we call Proximity and is computed as:

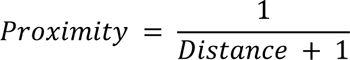

The term “+” prevents infinities at Distances of zero (*i.e*., when the gene is self-references).

We used two approaches to predict ageing-relatedness: 1. A ML approach using a balanced random forest algorithm with nested (CV) (better explained in the next section). This method identifies non-linear relationships in gene proximity to ARDs or ARCs. Genes are given an ageing-relatedness probability, and scores above a set threshold are labelled as ageing-related. 2. A direct method wherein each gene’s average association with its network targets (ARDs or ARCs) is calculated. These averages are then compared to a threshold to determine ageing-relatedness, emphasizing the significance of pure proximity averaging to multiple ageing-related diseases and communities. The effectiveness of both methods is gauged using the AUC metric.

### Datasets creation

We conducted 48 independent runs, combining eight algorithms and four networks, using the association of each gene to either ARDs or ARCs to compute ageing-relatedness. The features-retrieval algorithms were: 1 and 2) Proximity based on the shortest path between the reference gene and the closest gene of its target ARD or ARC. 3 and 4) Averaging shortest path proximities between the Reference Gene and each one of the genes of its Target ARD or ARC. 5 and 6) Counting the overlapping genes neighbouring the reference gene with the ARD-related genes for each ARD. Complementary information on Proximity, Proximity across networks, and the algorithms’ mechanisms and classification are depicted in Supplementary Figures S13, S14, S19, and Table S4, respectively. These values were used as ML features. The objective was a binary classification of genes as either ageing-related or non-ageing-related. Genes included in the ML model had at least one connection with another gene within the networks and were either disease-related, ageing-related, or their neighbours. We excluded those without a connection. We did so to focus on network topology to guide predictions and manage data imbalance. Therefore, while all the networks have either 9 or 58 features (ARCs or ARDs), the number of genes being classified and ageing-related genes varied across networks, with specific counts given for *ARC.PPI* (6721, 276, which are the number of genes under analysis and the number of ageing-related genes, respectively), ARC.COX95 (2463, 56), ARC.COX90 (7725, 147), and KEGGs (4891, 228), the number of classified genes diverges to 6721, 2463, 7725 and 4891.

### ML Algorithm & Nested CV

Balanced Random Forests from the python’s *imblearn* library (Lemaître *et al*., 2017) was chosen for ML due to the imbalanced classification challenge arising from a low count of ageing-related genes relative to non-ageing. Predictive performance was assessed using nested CV. The genes were divided into 10 stratified outer folds. Each ML algorithm underwent 10 iterations, each using one different outer fold for testing and the rest for training. Before each ML run, the hyperparameters were tuned using an inner 5-fold cross-validation, ensuring the best configuration was selected for the current iteration of the outer CV. The overall algorithm’s predictive accuracy was derived as the average of the 10 outer CV iterations. Predicted probabilities were converted into class labels using a 0.5 threshold.

### Ageing-related Genes Prediction & Enrichment Analysis

We aimed to identify novel Ageing-related genes by spotting the False Positives (FP) among those annotated as not-ageing-related but still having among the highest ML-based predicted probability of ageing-relatedness. These FP genes could be potential candidates for future experiments. Combining predictions from all outer testing splits, we focused on FP genes with a probability ≥ 0.5. We selected the top 30 FP genes from the best models as candidate ageing-related genes for further lab validation. The criterion was chosen to ensure consistent gene numbers across networks for enrichment analysis, and these genes were then used for enrichment analysis on the *gProfiler* website (Raudvere *et al*., 2019).

## Acknowledgments

GDVM thanks his sister, Laura Sujei Vega Magdaleno, for her support in enhancing the quality of some of the explanatory figures in this work.

## Authors’ contributions

GDVM and JPM conceived and planned the experiments. GDVM collected the data, which was partially generated by JPM’s team, and carried out the experiments. JPM contributed to the biological interpretation of the results. GDVM wrote the manuscript. JPM provided critical feedback and helped shape the research, analysis, and manuscript. All authors read and approved the final manuscript.

## Supplementary Materials

**Figure S1.** Age-of-onset-based Clustering of the 11 UK Biobank self-reported cancers with at least 2000 cases. **Figure S2**. GWAS analysis of the 9 filtered cancer ARDs computed from UK Biobank. **Figure S3**. Classification of 58 phenotypes of ARDs in ARCs according to the grouping hierarchy of UK Biobank’s self-reported diseases and cancers. **Figure S4**. Relationships between genes associated with ageing and various ARCs. **Figure S5**. *ARC.PPI* network. **Figure S6**. Genetic distances at the *ARC.PPI* network. **Figure S7**. *ARC.COX*_90_ Network. **Figure S8**. Genetic distances at the *ARC.COX*_90_ network. **Figure S9**. *ARC.COX*_95_ Network. **Figure S10**. Genetic distances at the *ARC.COX*_95_ network. **Figure S11**. *ARC.KEGG* Network. **Figure S12**. Genetic distances at the *ARC.KEGG* network. **Figure S13**. Representation of Proximity. **Figure S14**. Proximity Measures across *GenAge*_*Hum*_, *GenAge*_*Mod*_, Diseases, Neighbours and Others. **Figure S15**. Correlation between *Pleiotropy* and *iARC_Interactors*. **Figure S16**. Intra-community coexpression differences. **Figure S17**. Corrected Specificity Differences. **Figure S18.** Gene expression levels across of various tissues in the GTEx database for genes associated to each ARC. **Figure S19**. Proximity-based ML-features definitions. **Table S1**. Top 10 biological processes go terms associated with *GenAge*_*Hum*_, *GenAge*_*Mod*_, and their intersection. **Table S2**. Biological process GO terms associated with immunological systemic disorders. **Table S3**. Topological properties of gene groups across the ARC networks. **Table S4**. Gene-ARD/ARC proximity algorithms. **Table S5**. AUC results depicting the *GenAge*_*Hum*_-relatedness prediction for each network. **Table S6.** Top 10 ageing-related gene candidates based on *ARD*. *PPI*. **Table S7.** Top 10 ageing-related gene candidates based on *ARC.KEGG*. **Table S8.** Biological processes GO terms associated with the top 30 ageing gene candidates of the *ARC.PPI* and *ARC.KEGG* networks.

## Availability of data and materials

The datasets and code generated and/or analysed during the current study are available in the https://github.com/GusDany3691/Dual_Nature_of_Genes_Linking_ARDs/tree/main repository.

## Declarations

Not applicable.

## Ethics approval and consent to participate

Not applicable.

## Consent for publication

Not applicable.

