## Supplementary Materials for "Pleiotropy and Disease Interactors: The Dual Nature of Genes Linking Ageing and Ageing-related Diseases"

#### Cancer ARDs

Figure S1 in the study presents a classification of all cancers in the UK Biobank that exceeded the threshold of at least 2000 individuals. These cancers were categorized based on their onset patterns over time, resulting in two groups: those exhibiting exponential growth starting around the age of 50, and others showing a linear increase from approximately age 20, with some presenting in middle age.

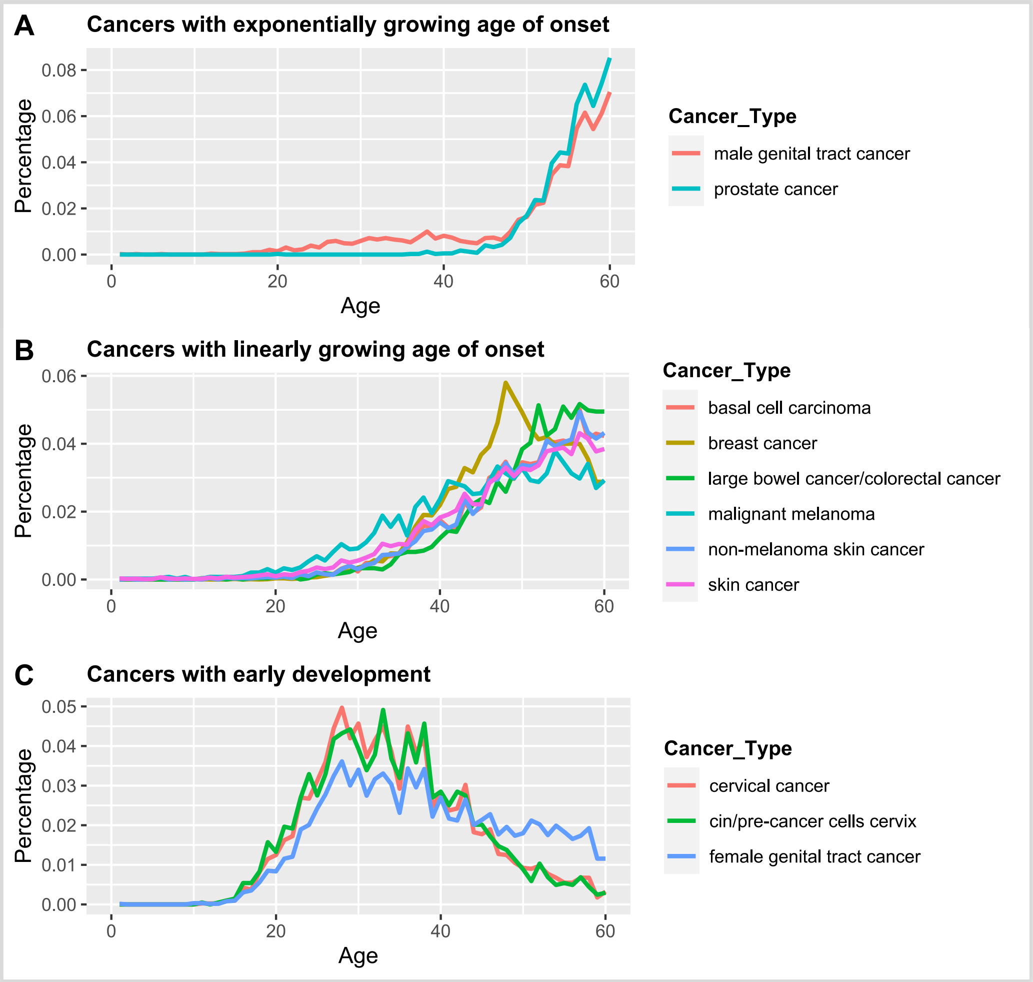


**Figure S1.** Age-of-onset-based Clustering of the 11 UK Biobank self-reported cancers with at least 2000 cases. A) Exponential growth, B) Linear growth, C) Early development. The clusters in A and B are herein considered cancer ARDs. Cancers of early development C are not considered in further analysis.

Figure S2 displays the genomic analyses of all aging-associated cancers (those increasing linearly and exponentially with age) examined in this study. It demonstrates that each of these cancers has statistically significant terms at various loci.

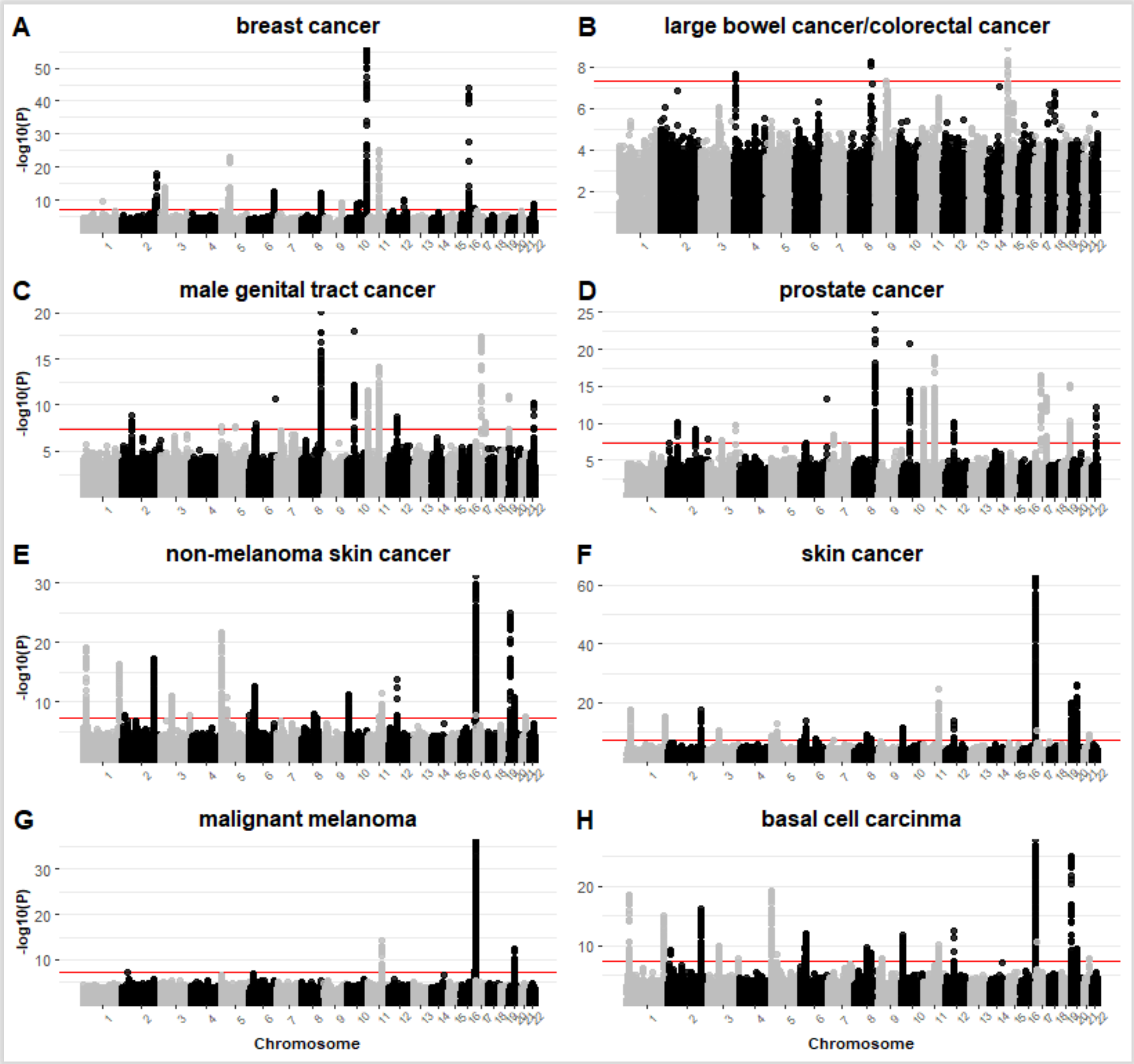


**Figure S2.** GWAS analysis of the 9 filtered cancer ARDs computed from UK Biobank. **A** Breast Cancer. **B** Large bowel cancer/colorectal cancer. **C** male genital tract cancer. **D** Prostate cancer. **E** Non-melanoma skin cancer. **F** Skin cancer. **G** Malignant melanoma. **H** Basal Cell Carcinoma.

### Communities of ARDs (ARCs)

The study delineates communities of age-associated diseases, identifying a total of 58 illnesses linked to aging. These are distributed among nine non-overlapping groups of aging-related communities. The specific communities are detailed in Figure S3 and include Cancer, Hematological-Dermatological, Non-Immunological Disorders, Musculoskeletal-Trauma, Neurobiology and Eye-Psychiatry, Gastrointestinal-Abdominal, Renal-Urology, Endocrine-Diabetes, and Cardiovascular. The Cancer community encompasses those detailed in sections 1 and 2 of the study.

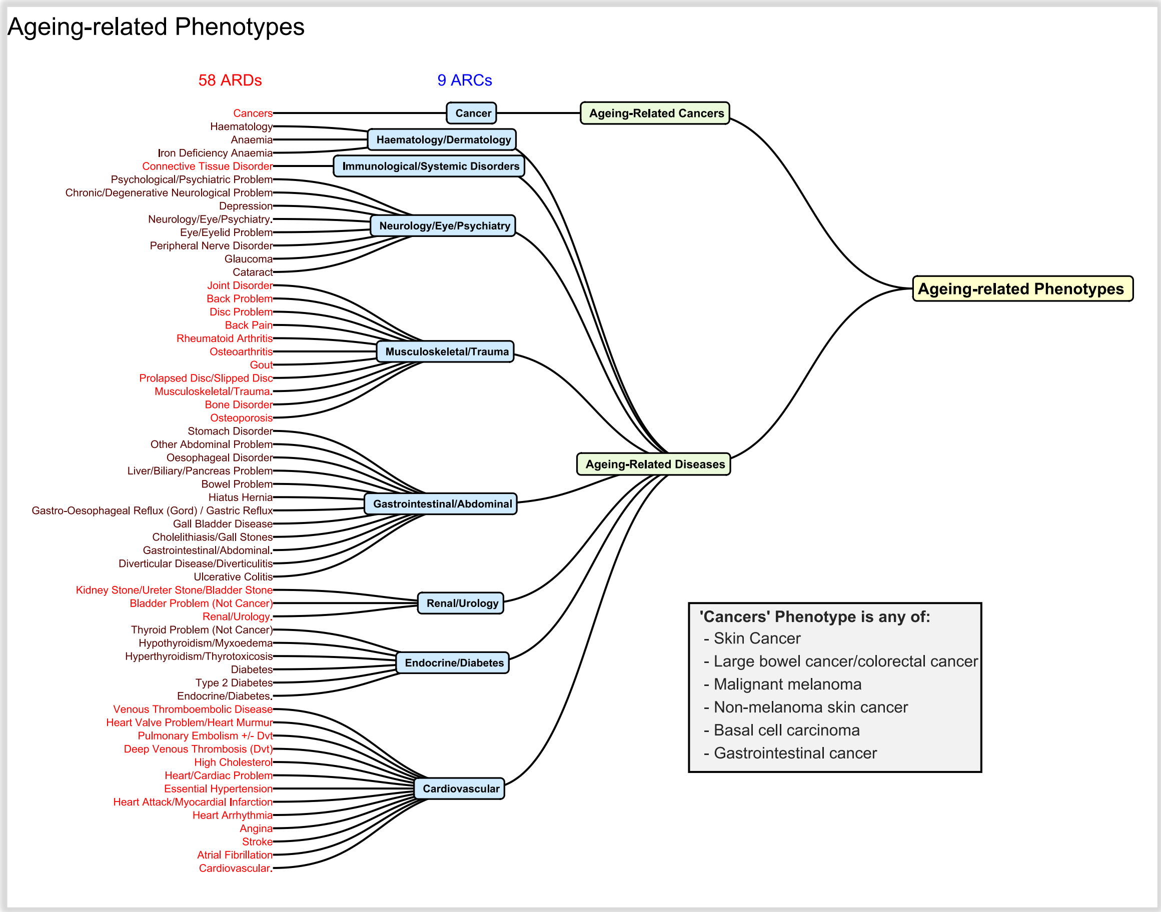


**Figure S3**. Classification of 58 phenotypes of ARDs in 9 ARCs according to the grouping hierarchy of UK Biobank's selfreported diseases and cancers. Note that we grouped all cancers on one single community and phenotype, 'cancer' and 'cancers', respectively, to avoid biasing our findings towards cancer-related genes whose genes may not only activate due to ageing but also to environmental factors.

### Relationships between genes associated with ageing and ARCs

Table S1 depicts a GO terms (Biological Processes) enrichment analysis to identify the functional differences between the set of $GenAge_{Hum}$-associated genes that excludes overlapping genes with $GenAge_{Mod}$ (i.e., $GenAge_{Hum}$-exclusive genes); the set of $GenAge_{Mod}$-associated genes that

**Table S1.** Top 10 biological process go terms associated with $GenAge_{Hum}$- and $GenAge_{Mod}$-exclusive genes as well as with the $GenAge_{Hum}$ - $GenAge_{Mod}$ intersection. Note how all these groups are mutually exclusive. The Term size is the number of genes associated with the GO term at the row. Query Size is the number of queried genes and intersection size is the number of queries genes associated with the GO term at hand.

| GenAge Set | Term Name | Term id | Adjusted p_value | Term Size | Query Size | Intersection Size |
|---|---|---|---|---|---|---|
| $GenAge_{Hum}$ Exclusive genes | apoptotic process | GO:0006915 | 3.69E-51 | 1910 | 176 | 97 |
| | cellular response to chemical stimulus | GO:0070887 | 1.07E-47 | 2691 | 176 | 107 |
| | response to UV | GO:0009411 | 6.24E-18 | 151 | 176 | 22 |
| | positive regulation of morphogenesis of an epithelium | GO:1905332 | 4.11E-05 | 37 | 176 | 7 |
| | regulation of cholesterol transport | GO:0032374 | 9.14E-05 | 62 | 176 | 8 |
| | B cell lineage commitment | GO:0002326 | 0.000136 | 6 | 176 | 4 |
| | nucleus organization | GO:0006997 | 0.00084 | 146 | 176 | 10 |
| | telomeric D-loop disassembly | GO:0061820 | 0.001116 | 9 | 176 | 4 |
| | regulation of mitochondrial membrane potential | GO:0051881 | 0.005053 | 73 | 176 | 7 |
| | acute-phase response | GO:0006953 | 0.007142 | 50 | 176 | 6 |
| $GenAge_{Mod}$ Exclusive genes | organonitrogen compound metabolic process | GO:1901564 | 9.87E-59 | 6362 | 994 | 547 |
| | generation of precursor metabolites and energy | GO:0006091 | 8.27E-30 | 514 | 994 | 98 |
| | macroautophagy | GO:0016236 | 1.99E-22 | 324 | 994 | 68 |
| | organic anion transport | GO:0015711 | 4.71E-20 | 435 | 994 | 76 |
| | membrane organization | GO:0061024 | 1.58E-09 | 813 | 994 | 87 |
| | synaptic vesicle cycle | GO:0099504 | 1.77E-09 | 200 | 994 | 37 |
| | unsaturated fatty acid metabolic process | GO:0033559 | 1.11E-07 | 111 | 994 | 25 |
| | mitochondrial transmembrane transport | GO:1990542 | 4.07E-06 | 61 | 994 | 17 |
| | ribosome biogenesis | GO:0042254 | 8.81E-06 | 327 | 994 | 42 |
| | clathrin-coated vesicle cargo loading, AP-3-mediated | GO:0035654 | 0.000155 | 7 | 994 | 6 |
| Intersection between $GenAge_{Hum}$ and $GenAge_{Mod}$ | cellular response to stress | GO:0033554 | 6.54E-37 | 1925 | 121 | 69 |
| | response to oxygen-containing compound | GO:1901700 | 1.52E-33 | 1774 | 121 | 64 |
| | endocrine system development | GO:0035270 | 9.51E-07 | 141 | 121 | 11 |
| | histone H3 deacetylation | GO:0070932 | 2.59E-05 | 6 | 121 | 4 |
| | response to epidermal growth factor | GO:0070849 | 0.000475 | 47 | 121 | 6 |
| | Ras protein signal transduction | GO:0007265 | 0.006799 | 337 | 121 | 11 |
| | embryonic cleavage | GO:0040016 | 0.017028 | 8 | 121 | 3 |
| | neuron projection organization | GO:0106027 | 0.021123 | 89 | 121 | 6 |
| | regulation of cysteine-type endopeptidase activity involved in apoptotic process | GO:0043281 | 0.021576 | 187 | 121 | 8 |
| | odontogenesis of dentin-containing tooth | GO:0042475 | 0.028912 | 94 | 121 | 6 |

excludes overlapping genes with $GenAge_{Hum}$ (i.e., $GenAge_{Mod}$-exclusive genes); and the intersection between $GenAge_{Hum}$ and $GenAge_{Mod}$. For ease of comparison and visualization, only the top ten most relevant GO terms were depicted per group. As displayed, $GenAge_{Hum}$-exclusive genes related the most (*pvalue ≤ 6.24E-18*) with *apoptotic process*, *cellular response to chemical* *stimulus* and *response to UV*. $GenAge_{Hum}$-exclusive genes, on the other hand, related the most (*pvalue ≤ 4.71E-20*) with *organonitrogen compound metabolic process*, *generation of precursor metabolites and energy*, *macroautophagy* and *organic anion transport*. The intersection between $GenAge_{Hum}$ and $GenAge_{Mod}$ was mostly inclined (1.52E-33) towards cellular response to stress and response to oxygen-containing compound. Other GO terms existed at the top 10 but no overlap was observed between these groups. At much, *apoptotic process* from $GenAge_{Hum}$-exclusive genes was related to *regulation of cysteine-type endopeptidase activity involved in apoptotic process* that corresponds to the intersection between $GenAge_{Hum}$ and $GenAge_{Mod}$, as the two GO terms refer to processes involved in programmed cell death.

Figure S4 illustrates the shared genes between ARCs and the GenAge datasets. On the other hand, Tabe S2 depicts the GO terms for the genes with the highest pleiotropy (equal or higher than 4) that were mostly associated with immunological systemic disorders.

**Table S2**. Biological process GO terms associated with immunological systemic disorders

| Term Name | Term id | Adjusted p_value | Term Size | Query Size | Intersection Size |
|---|---|---|---|---|---|
| nucleosome assembly | GO:0006334 | 5.19E-10 | 115 | 53 | 10 |
| urate transport | GO:0015747 | 1.5E-05 | 10 | 53 | 4 |
| innate immune response in mucosa | GO:0002227 | 0.001413 | 28 | 53 | 4 |
| T cell receptor signaling pathway | GO:0050852 | 0.002255 | 135 | 53 | 6 |
| antibacterial humoral response | GO:0019731 | 0.040991 | 64 | 53 | 4 |

### ARC Genetic Networks

Figures S5, S7, S9 and S11 depict the $ARC.PPI$, $ARC.COX_{90}$, $ARC.COX_{95}$, and $ARC.KEGG$ networks, respectively. Figures S6, S8, S10 and S12 illustrate heatmap representations of genetic distances between genes and ARCs, as determined by their position within the same networks, respectively. Columns represent ARCs. Rows represent genes and are split as follows: *Diseases* (ARCs-related genes), $GenAge_{Hum}$, $GenAge_{Mod}$ and *Neighbours*. *Neighbours* are genes that, within the corresponding network, are adjacent to *Disease*-related genes. Each cell represents the number of intermediate genes between the reference gene and the closest gene associated with the ARC of interest. The strongest red colour indicates a *distance=0*, meaning direct GWAS association between the gene and at least one ARD of the ARC of the corresponding column. The

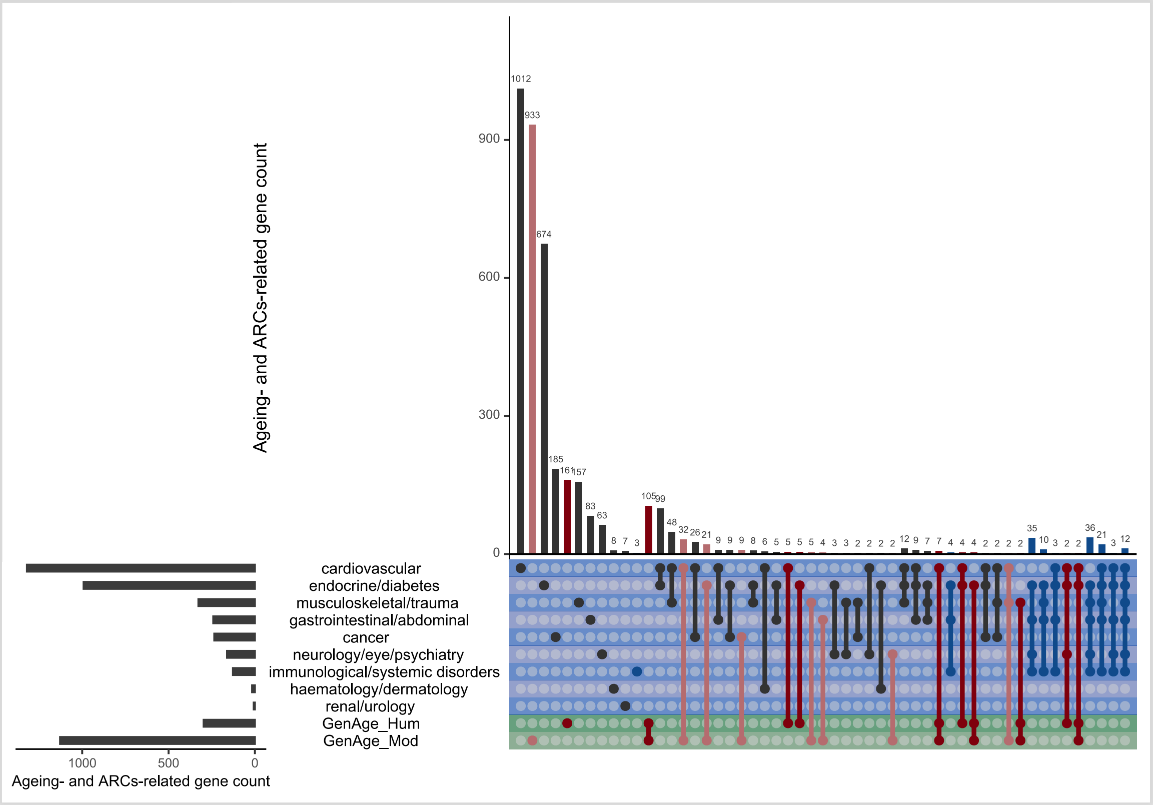


**Figure S4**. Relationships between genes associated with ageing and various ARCs. A deep red colour was employed to denote genes linked to human aging, while a lighter red indicates genes associated with aging in model organisms. A strong blue colour is utilized for lines corresponding to immunological disorders, indicating the widely extended relationships existing between them and other ARCs. Dark-coloured lines are used to represent the intersections that exist between these ARCs-related genes when neither aging- nor immune disease-related genes are involved. Green rows represent the set of genes associated with any of the two categories of ageing-related genes. The rows in blue depict all the ARCs. The gene sets comprising only a single gene, though in some instances they may be associated with more than one ARC, were excluded for the sake of visual clarity. This exclusion does not importantly impact the relationships depicted in the graph, as these single-gene sets constitute a minority.

lighter red colour indicated a *distance=1*, meaning that the gene indirectly interacts to with the ARC through interaction with a GWAS-significant gene associated to the ARC (i.e., *iARC_Interactions*). The white and further green colours indicate longest distances under the same principle. In the context of our work, we don't consider such larger distances as indirect associations or *iARC_Interactions*, since their farthest positions could dilute their effects over the distant ARCs. The dark green colour "Inf" means that the genes described in the rows cannot be connected to the ARC. To the right, there are two annotation heatmap bars. The first bar represents number of *iARC_Interactions*, which is the number of ARCs with which the gene indirectly interacts at a *distance=1* (Supplementary - Pleiotropy and Indirect Interactor definitions). The last bar on the right depicts the mean distance of the gene to all ARCs.

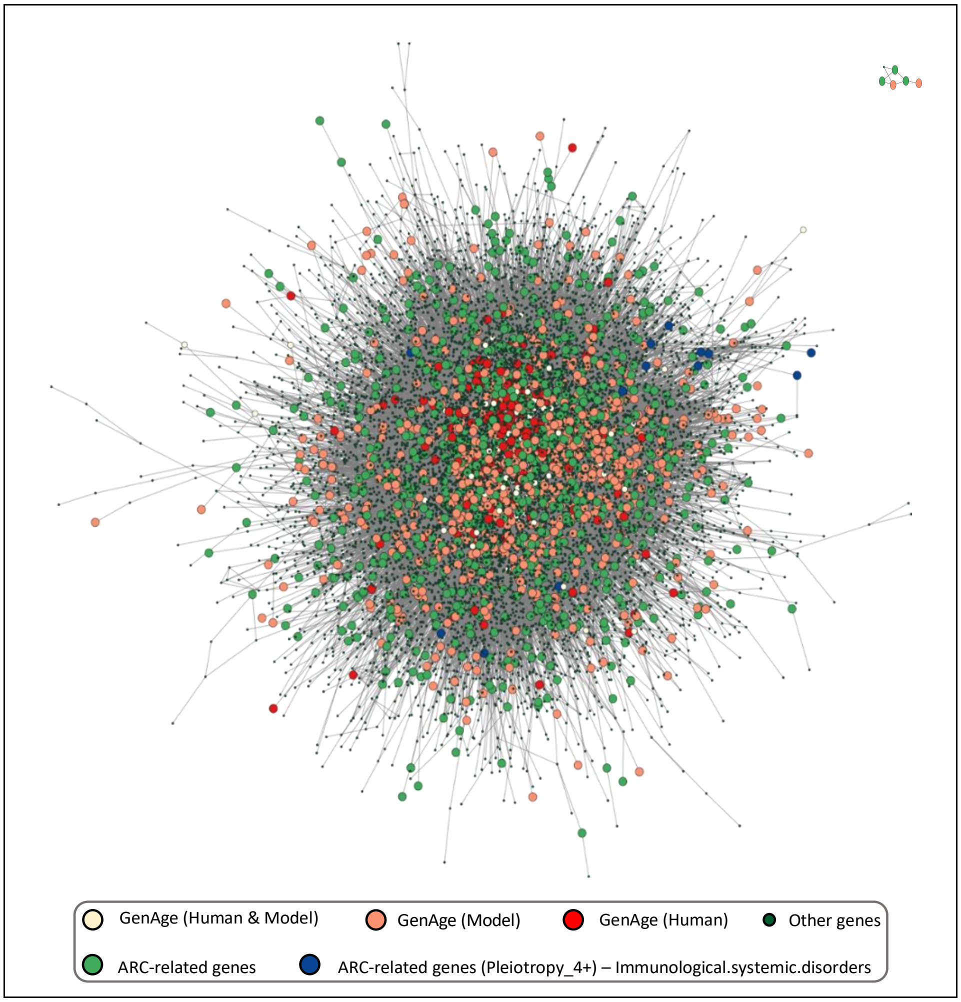


**Figure S5**. $ARC.PPI$ network. Genes are color-coded: *Disease* (green), $GenAge_{Hum}$ (red), $GenAge_{Mod}$ (orange), $GenAge_{Hum}$ & $GenAge_{Mod}$ (light yellow), and Immunological Sytemic Disorders genes (blue). Background genes have a darker green shade and smaller nodes. Only aging- and *Disease*-related genes that belong to the PPI network are displayed. Clusters of genes with less than five genes were excluded for ease of visualization.

#### $ARC.PPI$ Network

Figure S5 displays the $ARC.PPI$ network, revealing a predominant clustering of almost all genes into a large group that includes all proteins, with a minor presence of a smaller group with seven genes. Within this network, $GenAge_{Hum}$-related genes are centrally positioned with minimal dispersion, whereas $GenAge_{Mod}$-related, while also central, exhibit greater dispersion, indicated by their larger coverage area. *Disease*-associated genes are even more widely dispersed across the

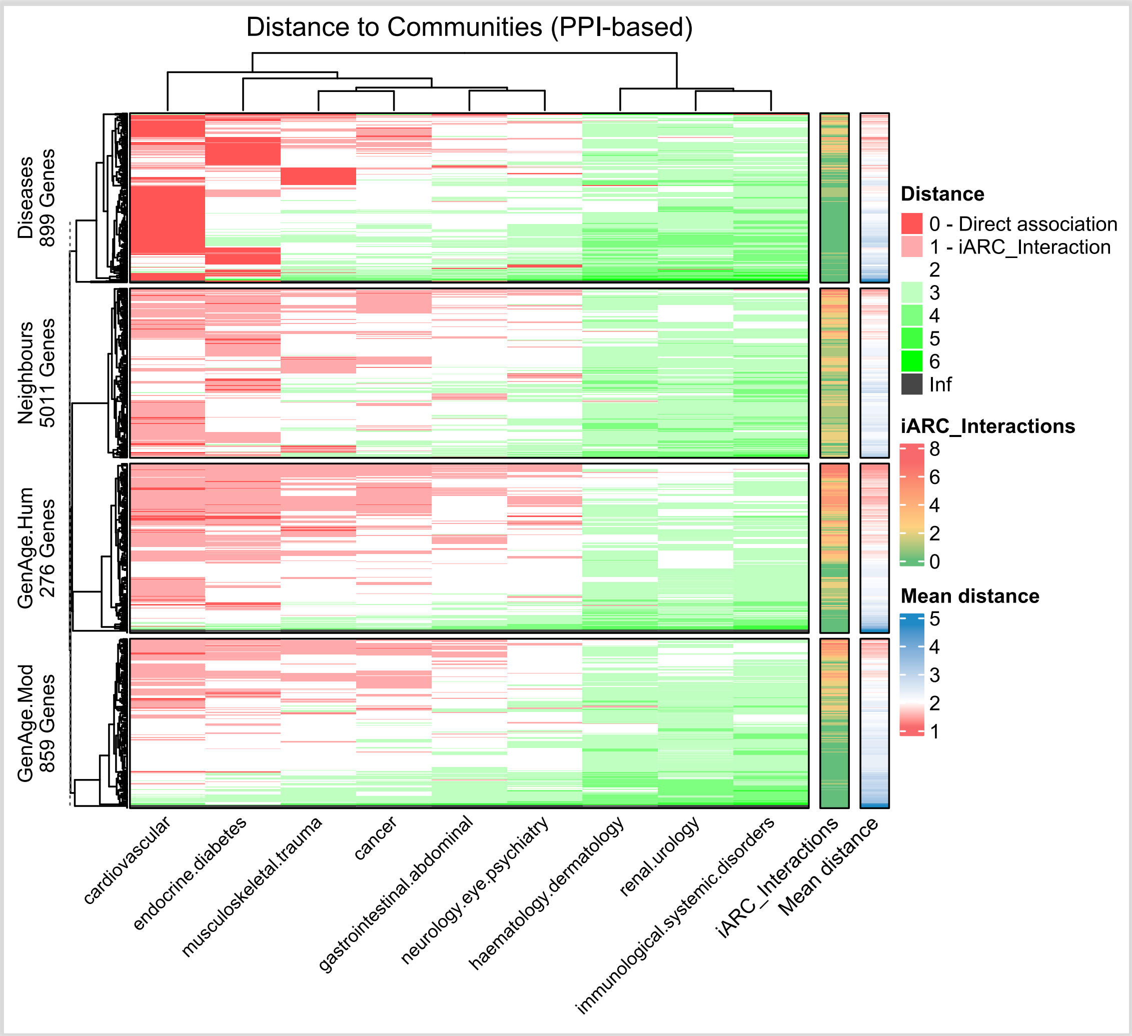


**Figure S6.** Genetic distances at the ARC.PPI network. Heatmap representation of genetic distances between a genes and ARCs, as determined by their position within the $ARC.PPI$ gene interaction network. Full explanation of this and the remaining similar figures is provided in the main text at the beginning of this section (second paragraph of section - ARC Genetic Networks).

network, spanning a larger area than $GenAge_{Mod}$-*related* genes. Genes shared between $GenAge_{Hum}$ and $GenAge_{Mod}$ occupy a central but non-concentrated location, distributed within a zone primarily occupied by $GenAge_{Hum}$-related genes. Remarkably, genes associated to immunological disorders, despite their direct connections to many ARCs, are situated at the peripheries of the network, mostly in mid-range or intermediate positions rather than central ones, with a small subset clustering in the upper right part of the figure, albeit with some scattered in various other locations.

Figure S6 reveals that the majority of genes at the $ACR.PPI$ network tended to have a relatively close distance (*distance<=2*) to most ARCs except Immunological, renal and haematological disorders, which tended to be at a remotest position with *distance>=3* across all groups of genes, with some counted exceptions, particularly at the *GenAge* groups, where more genes reached *distance=2*. Genes associated with *Diseases* predominantly associated with cardiovascular and endocrine disorders, with a smaller proportion connected to other ARCs. Apart from their own associated ARCs (i.e., strong red, *distance=0*), these *Disease*-related genes generally maintained a distance of about two intermediate genes to the next closest ARC, particularly musculoskeletal, cancer, neurological, and gastrointestinal, as can be appreciated by the predominantly white colour (*distance=2*) at these ARCs. This suggests that *Disease*-related genes tend to lack *iARC_interactions* (i.e., first order associations to ARCs) in a PPI environment, as will be statistically proven in the next sections. Moreover, there was a minority of the *Disease*-related genes that showed limited reachability despite their connection in the PPI network, confined to their own ARC or very few others, this was marked by the grey colour at the bottom of the *Diseases* window, indicating infinite distance to other ARCs.

Human ageing-associated genes displayed a clearer pattern of *iARC_interactions* when compared to the other groups (i.e., light red, *distance=1*) as roughly half of their genes tended to indirectly interact with two or more ARCs up to 6. Although most $GenAge_{Hum}$-associated genes tended to show a consistent indirect association with cardiovascular and endocrine ACRs, they also indirectly connected with other ARCs, and it also tended display relatively shortest distances to renal/urology ARCs than the other groups. On the other hand, *Neigbhours* and $GenAge_{Mod}$-associated genes mostly presented *iARC_interactions* with just one or two ARCs, while keeping the rest at a longer *distance=2*, with a minority of genes indirectly interacting with three or more ARCs. The *iARC_Interactions* and *Mean_distance* bars visually suggest the higher proportion of *iARC_Interactions* in human aging-related genes, discernible through more pronounced red and yellow coloration, this is tested statistically in the main paper's section "Disease Interactors". The mean distance is not as clearly distinguishable between groups but it is also tested statistically in Supplementary Section "Proximity Analysis".

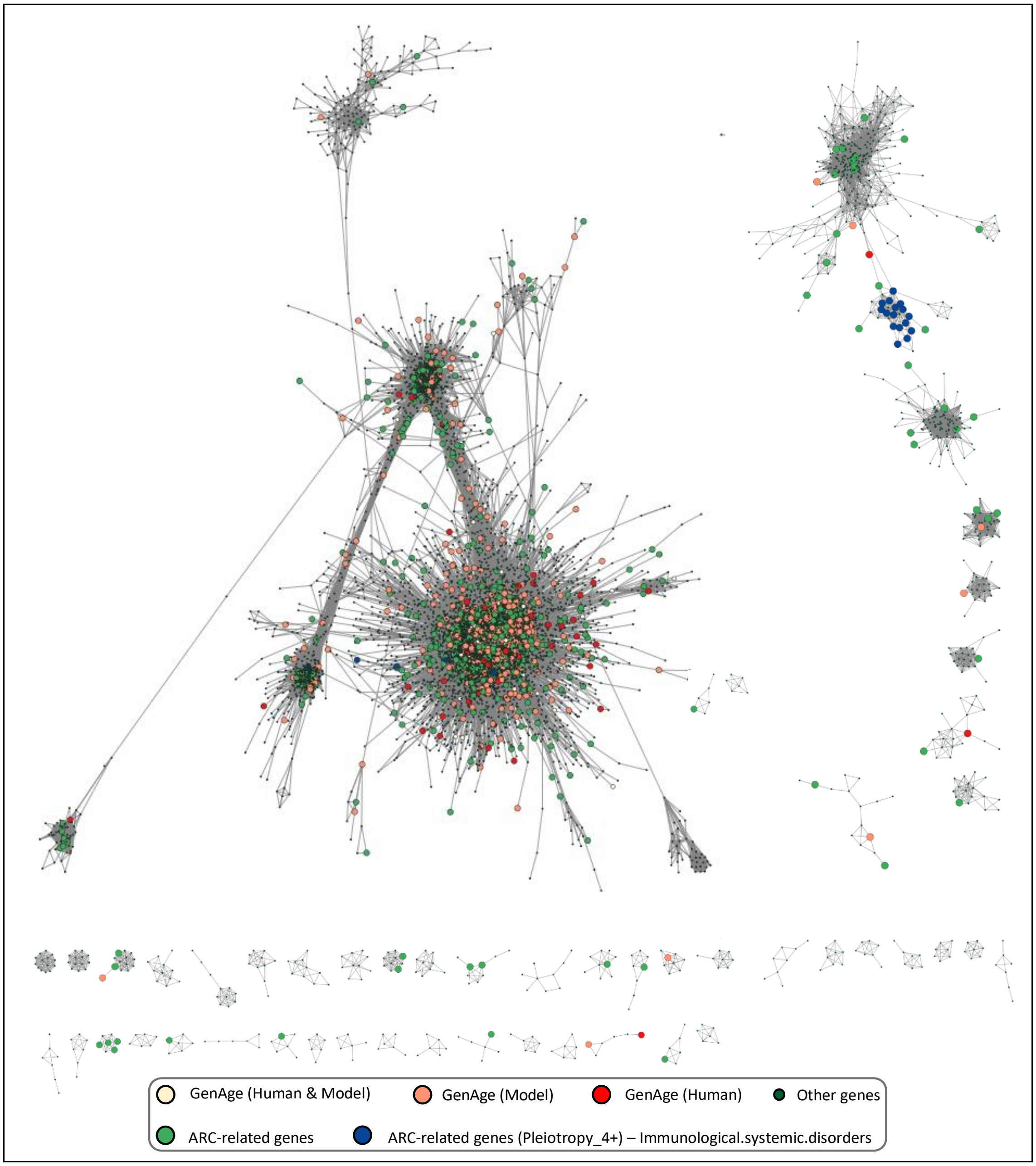


**Figure S7** $ARC.COX_{90}$ Network. Genes are color-coded: *Disease* (green), $GenAge_{Hum}$ (red), $GenAge_{Mod}$ (orange), $GenAge_{Hum}$ & $GenAge_{Mod}$ (light yellow), and Immunological Sytemic Disorders genes (blue). Background genes have a darker green shade and smaller nodes. Only aging- and *Disease*-related genes that belong to this coexpression network are displayed. Clusters of genes with less than five genes were excluded for ease of visualization.

#### $ARC.COX_{90}$ Network

Figure S7 displays the $ARC.COX_{90}$ Network. This network was characterized by its division of genes across several isolated clusters, with a dominant primary cluster containing three or four large gene groups where most human and model animal aging-related genes, alongside a majority

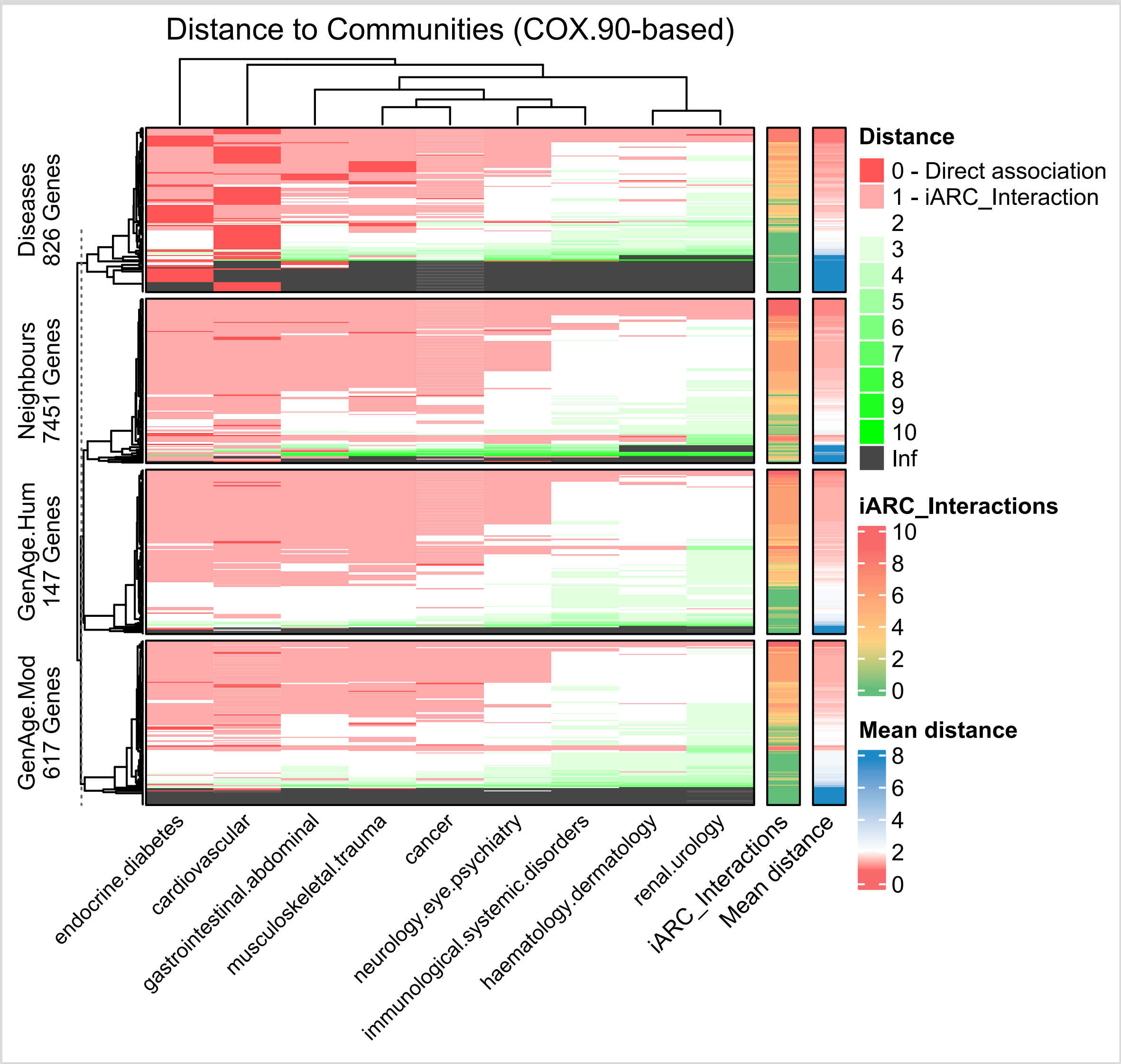


**Figure S8.** Genetic distances at the ARC.COX90 network. Heatmap representation of genetic distances between a genes and ARCs, as determined by their position within the the $ARC.COX_{90}$ gene interaction network. Full explanation of this and the remaining similar figures is provided in the main text at the bebanging of this section (second paragraph of section - ARC Genetic Networks).

of *Disease*-associated genes, are found, excluding those related to immunological diseases or high *Pleiotropy* to ARCs. Ageing-related genes, both human- and model organisms-related, were notably concentrated within the largest node of this main cluster, appearing central yet dispersed, while model organism aging-related genes, also in this module, displayed a more confined dispersion. This main cluster also included various *Disease*-associated genes near the $GenAge_{Mod}$-associated genes.

Further, a few ageing- and *Disease*-related genes, mainly from model organisms, were located within the second and third nodes of this cluster. Separately, the second-largest cluster featured a sub-module where genes associated with immunological disorders or high *Pleiotropy* were densely clustered, forming a highly coexpressed and isolated group. Meanwhile, smaller subsequent clusters contained a minority of isolated *Disease*- and Ageing-related genes. Genes shared between $GenAge_{Hum}$- and $GenAge_{Mod}$ were present in the main module but were less discernible due to the high density of genes. The cluster-based organization of this network resulted in many genes being isolated from each other when in different clusters, creating infinite distances between them.

Figure S8 illustrates that, differently to $ARC.PPI$, the *Disease*-related genes at the $ARC.COX_{90}$ demonstrated more *iARC_interactions* with a variety of ARCs, as indicated by widespread light red coloration. These indirect associations extended across most groups but are less pronounced within renal, haematological, and immunological communities. The differences in connectivity to ARCs between *Disease*-, $GenAge_{Hum}$-, $GenAge_{Mod}$-, and *Neighbouring*-associated genes were subtle, with *Neighbouring* genes showing a slightly more extensive area of *iARC_interactions*. This pattern was partially reflected in the *iARC_interactions* and *Mean_Distance* bars. Notably, even in distant communities like renal, haematological, and immunological, the shortest connectivity distances were usually around two, indicating a closer association than that observed in the $ARC.PPI$ and $ARC.KEGG$ networks. Additionally, some *Disease*-related genes were isolated within the network, existing at an infinite distance from genes of other ARCs due to their placement in distinct coexpression clusters. This phenomenon of isolation was also observed among certain *GenAge* and *Neighbouring* genes, albeit less frequently.

#### $ARC.COX_{95}$ Network

The $ARC.COX_{95}$ network, as illustrated in Figure S9, was structured into numerous clusters which, as expected, are smaller and less numerous compared to $ARC.COX_{90}$. In contrast with $ARC.COX_{90}$ where the main cluster was divided into several modules, the principal cluster at $ARC.COX_{95}$ was more uniform and remained encompassing most $GenAge_{Hum}$-, $GenAge_{Mod}$- and *Disease*-related genes, excluding those associated with immunological diseases or high pleiotropy. In this main cluster, $GenAge_{Hum}$-related genes were notably sparse and scattered, not occupying a central position, in contrast to the more homogeneously dispersed $GenAge_{Mod}$-related genes. *Disease*-associated genes, however, tended to be centrally located. The network's secondary cluster was split into two modules linked by a few genes, including some associated with

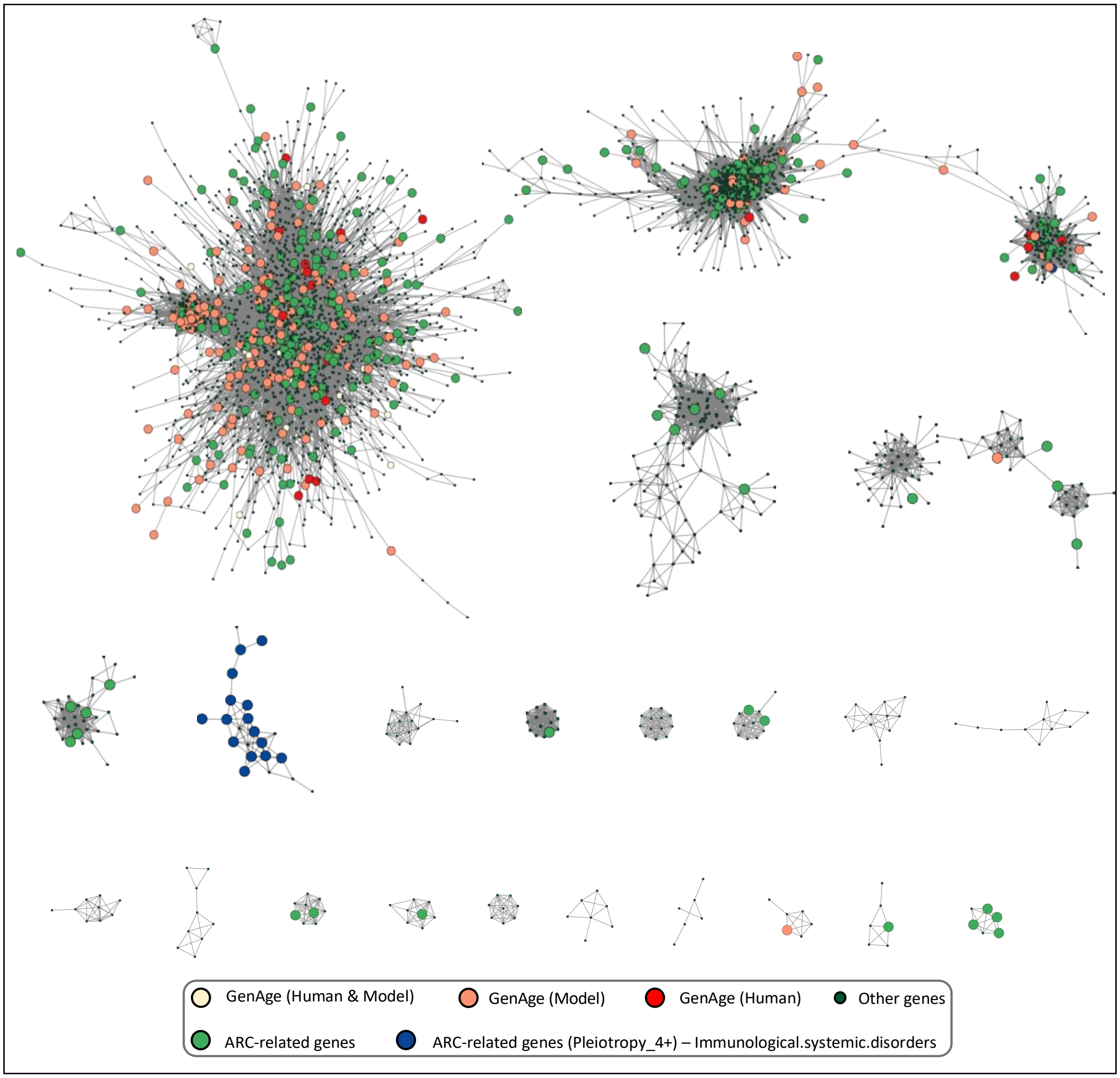


**Figure S9**. $ARC.COX_{95}$ Network**.** Genes are color-coded: *Disease* (green), $GenAge_{Hum}$ (red), $GenAge_{Mod}$ (orange), $GenAge_{Hum}$ & $GenAge_{Mod}$ (light yellow), and Immunological Sytemic Disorders genes (blue). Background genes have a darker green shade and smaller nodes. Only aging- and *Disease*-related genes that belong to this coexpression network are displayed. Clusters of genes with less than five genes were excluded for ease of visualization.

$GenAge_{Mod}$. This cluster contained a mix of both $GenAge_{Hum}$ – and $GenAge_{Mod}$-related genes and still lacked any immunological disorder-related genes. Subsequent clusters have fewer *Disease*-associated genes, but one particular cluster was heavily concentrated with immunological disorders-related genes.

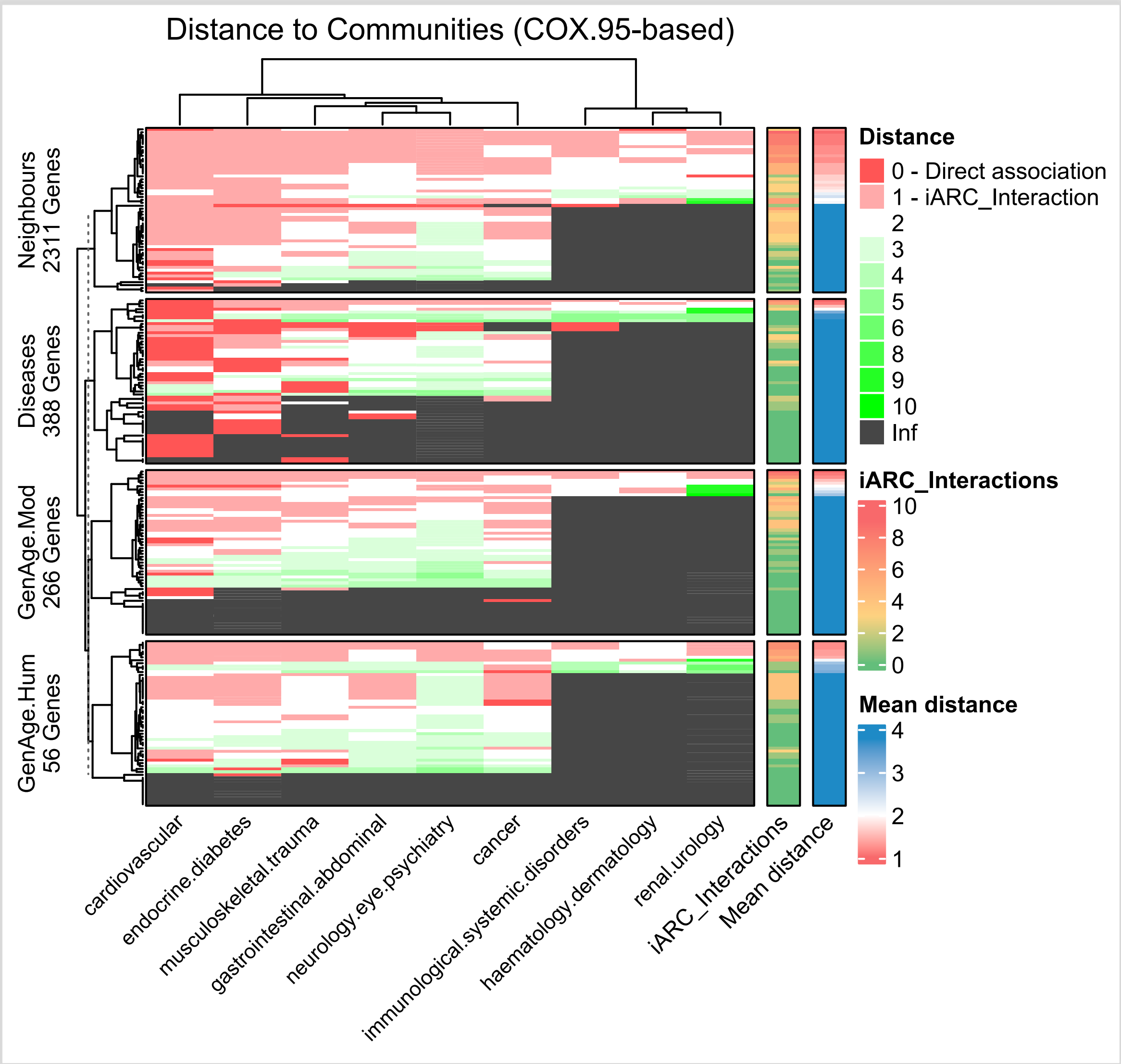


**Figure S10.** Genetic distances at the ARC.COX95 network. Heatmap representation of genetic distances between a genes and ARCs, as determined by their position within the $ARC.COX_{95}$ network. Full explanation of this and the remaining similar figures is provided in the main text at the bebanging of this section (second paragraph of section - ARC Genetic Networks).

Intriguingly, this cluster, while pleiotropically associated with a diverse range of ARCs, lacked genes exhibiting lower pleiotropy (i.e., *Diseases*-related genes not related to immunological-disorders) and did not include any $GenAge$-associated genes. This layout indicates that at least a subset of genes linked to immunological disorders, tended to coexpress in a collective yet isolated manner from other gene groups.

Figure S10 indicates that, in the $ARC.COX_{95}$ network, *GenAge*-related genes displayed a pattern similar to $ARC.COX_{90}$ in the sense that some of its genes established *iARC_Interactions* with multiple ARCs, but this connectivity was less pronounced compared to $ARC.COX_{90}$, involving only about 10% of genes. This network presented the least number of *GenAge*-associated genes, relative to the other networks. Moreover, $GenAge_{Hum}-$ and $GenAge_{Mod}$-related genes generally maintained a greater distance to ARCs with a small proportion of genes indirectly interacting with multiple ARCs at a *distance=1*, while the remaining GenAge-associated genes kept such indirect interactions with only one to zero ARCs. *Neighbours* genes exhibited less of this green coloration and maintained an average distance of around two, while *Disease*-related genes were usually one or two steps away from genes outside their specific ARC group. A notable aspect of this network was the considerable number of ARCs that remain mostly disconnected due to their genes being in isolated clusters, especially prevalent in renal, hematological, and immunological diseases. However, approximately 10-30% of each group still manageed to reach these ARCs indirectly or from a slightly greater distance. This highest level of gene disconnection is attributed to the formation of varied clusters and the limited number of genes exceeding the 95% coexpression threshold.

#### $\boldsymbol{ARC.KEGG}$ Networks

The $ARC.KEGG$ signalling network was primarily comprised of a single large principal cluster (Figure S11), similar to the $ARC.PPI$ network. It encapsulates the concatenation of all human pathways at the KEGG database. A few smaller clusters existed, mainly containing background genes with only one single *Disease*- or $GenAge_{Mod}$-related genes, though these were minor compared to the vast main cluster. Here, $GenAge_{Hum}$-related genes were predominantly positioned in a central yet dispersed manner, with some genes concentrated in the center and others scattered throughout. $GenAge_{Mod}$-related genes were similarly distributed, avoiding the network's extremes or 'leaves'. *Disease*-associated genes also tended to be centrally located, mirroring the $GenAge$-related genes' distribution. Intriguingly, genes associated with immunological disorders were positioned at the network's terminal points, clustering together in the same cascades and forming distinct groups. Despite their close grouping, there is a slight intermixture of $GenAge$- and less pleiotropic *Disease*-related genes nearby. This network maintains a hierarchical structure, ($ARC.KEGG$). Additionally, genes common to $GenAge_{Hum}$ and $GenAge_{Mod}$ are located in the main cluster, dispersed but relatively central, indicating their overlapping yet distinct roles within the signalling network's framework.

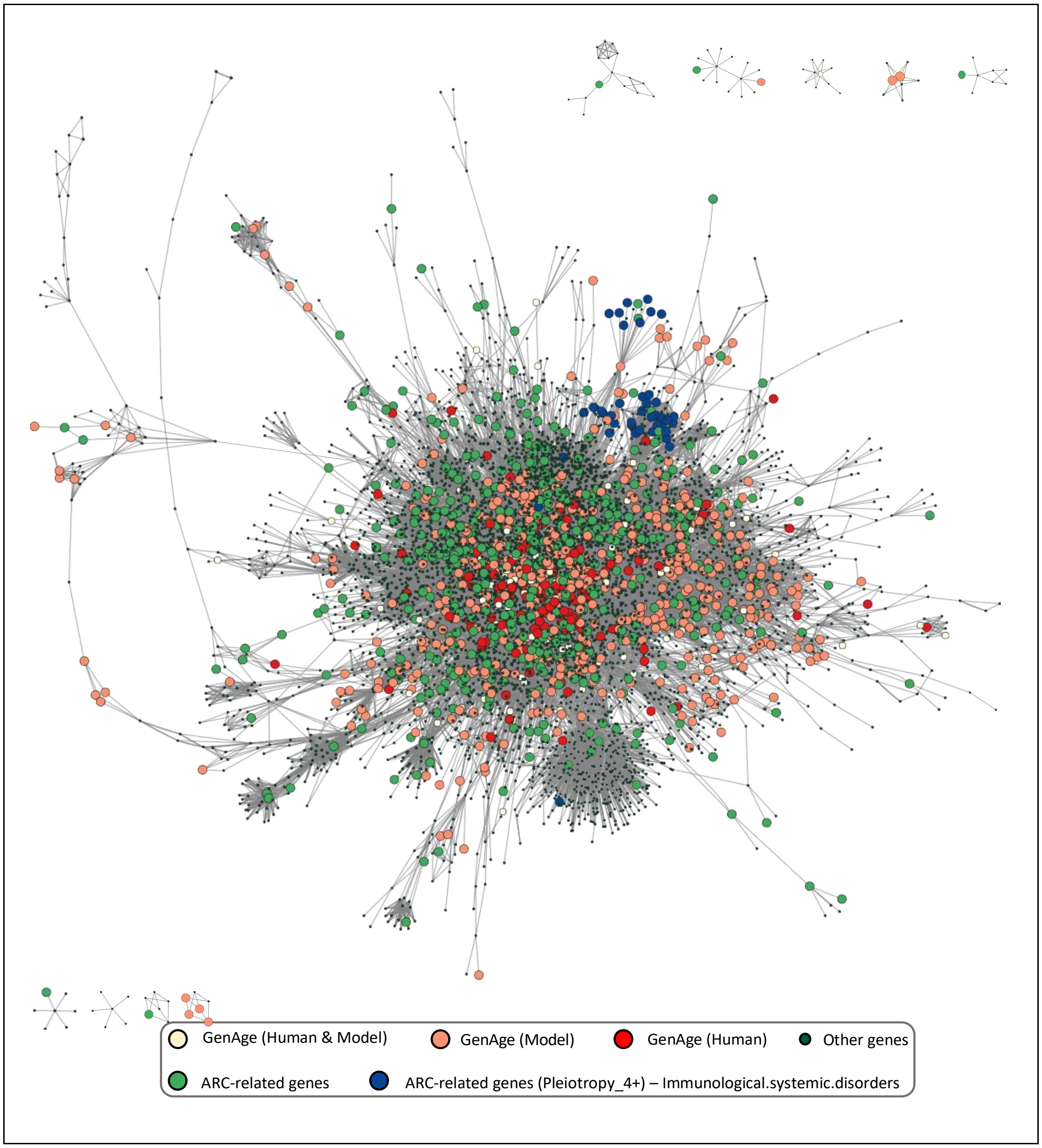


**Figure S11**. $ARC.KEGG$ Network. Genes are color-coded: *Disease* (green), $GenAge_{Hum}$ (red), $GenAge_{Mod}$ (orange), $GenAge_{Hum}$ & $GenAge_{Mod}$ (light yellow), and Immunological Sytemic Disorders genes (blue). Background genes have a darker green shade and smaller nodes. Only aging- and *Disease*-related genes that belong to this coexpression network are displayed. Clusters of genes with less than five genes were excluded for ease of visualization.

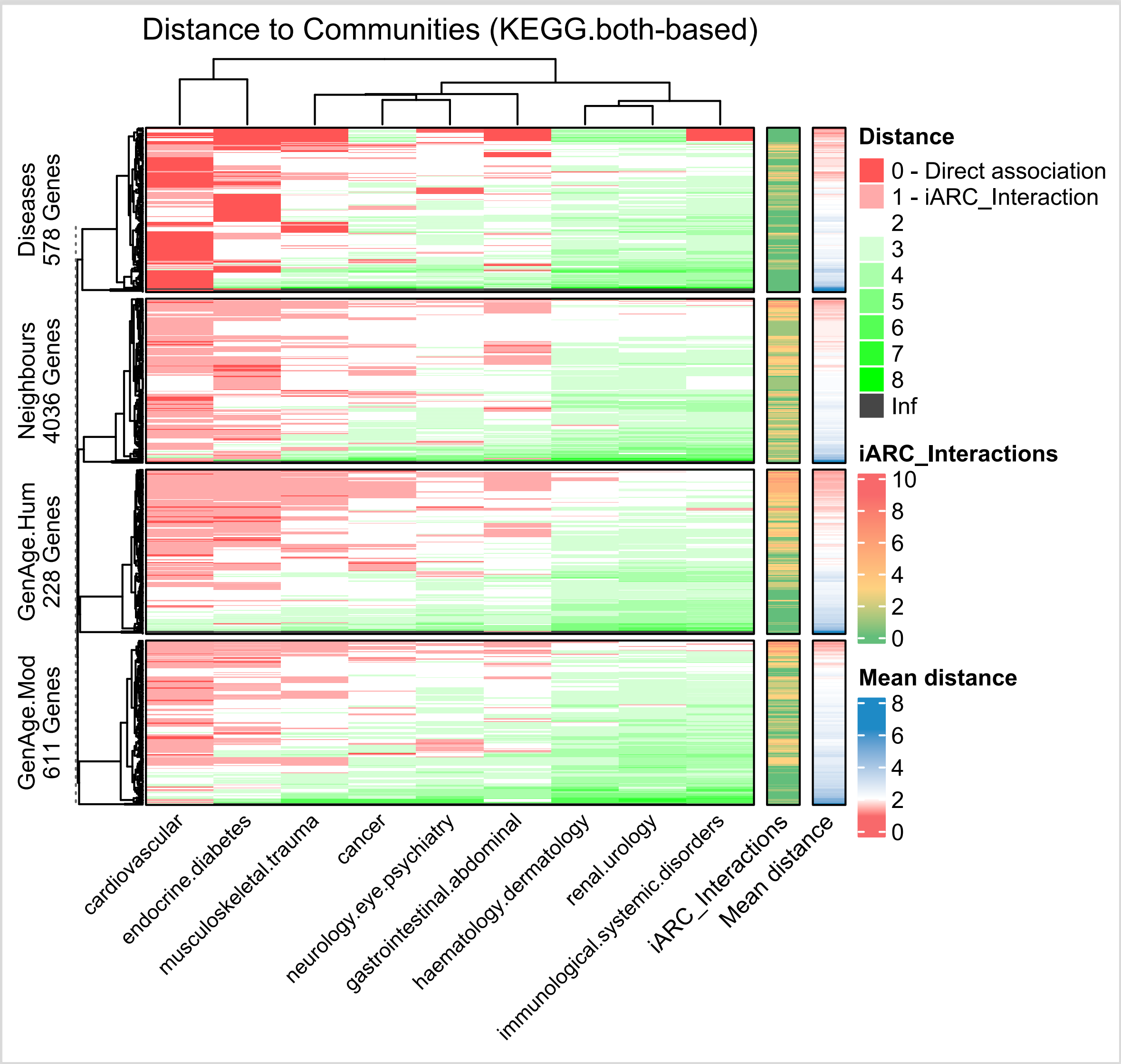


**Figure S12.** Genetic distances at the ARC.KEGG network. Heatmap representation of genetic distances between a genes and ARCs, as determined by their position within the $ARC.KEGG_{both}$ network. Full explanation of this and the remaining similar figures is provided in the main text at the bebanging of this section (second paragraph of section - ARC Genetic Networks).

*Disease*-related genes typically maintained direct connections (*distance=0*) with their respective ARCs but exhibited a *distance≥2* with other ARCs. $GenAge_{Hum}$-related genes showed most of its *iARC_Interactions* with cardiovascular and endocrine diseases, with a smaller percentage of its genes simultaneously *iARC_interacting* with gastrointestinal, musculoskeletal, and cancer communities $ARC.KEGG$ reaches a distribution of distances similar to that of $ARC.PPI$. *Neighbour* genes maintained *iARC_Interactions* with just few communities per gene, usually one or two, and mostly cardiovascular and endocrine disorders, followed by greater distances with the rest of communities.

$GenAge_{Mod}$-related genes were intermediate to $GenAge_{Hum}$ and *Neighbouring* genes in terms of proportion of *iARC_Interactions* across all KEGG networks, also keeping most of its association with cardiovascular and endocrine disorders. Immunological, renal, and haematological diseases remained the least connected.

#### Topological Properties of the Networks

Table S3 illustrates that, generally, $GenAge_{Hum}$-associated genes tended to exhibit greater centrality in terms of degree (i.e., the number of genes with which a gene interact) and betweenness (i.e., the number of times a gene acts as a bridge along the shortest path between two other genes) within the $ARC.PPI$ and $ARC.KEGG$ networks. This difference highlights the well-known bias due to more extensively investigated genes (Gillis et al., 2014). In contrast, $ARC.COX$ networks did not show *GenAge*-associated genes as highly connected compared to other groups. While $GenAge_{Mod}$-associated genes also trended towards high degree centrality, this was less pronounced than in $GenAge_{Hum}$. In the $ARC.COX$ networks, the $GenAge_{Mod}$-associated genes tended to have a higher degree centrality than $GenAge_{Hum}$, although they got a lower betweenness centrality.

*Disease*-related genes tended to display lower degree and betweenness centrality in the $ARC.PPI$ and $ARC.KEGG$ networks compared to both $GenAge_{Hum}$- and $GenAge_{Mod}$-associated genes, yet they maintained a slightly higher position than the background genes within the network (*Others* Group). Immunological disorders-related genes, characterized by high pleiotropy, showed the lowest centrality in terms of both degree and betweenness across all groups, including *Others*. This trend was consistent across all networks. The closeness centrality (i.e., how close a gene is to all other genes) of all genes in the network tended to be lower for *GenAge*-associated genes, but overall, differences between the various groups were minimal, with Immune disorders' and *Other's* genes showing slightly higher closeness centrality, with few exceptions. In the $ARC.PPI$ and $ARC.KEGG$ networks, genes exhibited lower closeness centrality than the $ARC.COX$ networks, with $ARC.COX_{95}$ presenting the highest.

**Table S3.** Topological properties of gene groups across the ARC-related networks. The sets of genes are associated with $GenAge_{Hum}$, $GenAge_{Mod}$, *Diseases (ARCs-related genes)* and Immunological Disorders. The group *Others* represent random genes outside any of these categories. For each group of genes, the mean value of Degree, Betweenness and Closeness Centralities are displayed. Similarly, the mean value of Clustering Coefficient and the mean Percentage of the Reference gene's neighbours associated with at least one Disease are presented.

| Network | Genes | Degree Centrality | Betweenness Centrality | Closeness Centrality | Clustering Coefficient | Percentage of Neighbour Genes related with *Diseases* |
|---|---|---|---|---|---|---|
| $ARC.PPI$ | $GenAge_{Hum}$ | 51 | 137325 | 2% | 13% | 10% |
| | $GenAge_{Mod}$ | 24 | 39423 | 1% | 22% | 9% |
| | *Diseases* | 13 | 16108 | 1% | 23% | 10% |
| | Immune Disorders. | 4 | 8031 | 0% | 19% | 8% |
| | Others | 10 | 10162 | 2% | 24% | 10% |
| $ARC.COX_{90}$ | $GenAge_{Hum}$ | 295 | 33768 | 8% | 68% | 9% |
| | $GenAge_{Mod}$ | 348 | 14043 | 5% | 70% | 9% |
| | *Diseases* | 287 | 15540 | 10% | 72% | 15% |
| | Immune Disorders. | 48 | 729 | 14% | 71% | 47% |
| | Others | 255 | 10557 | 14% | 74% | 8% |
| $ARC.COX_{95}$ | $GenAge_{Hum}$ | 46 | 1484 | 23% | 62% | 12% |
| | $GenAge_{Mod}$ | 57 | 2871 | 17% | 61% | 9% |
| | *Diseases* | 50 | 2032 | 24% | 63% | 25% |
| | Immune Disorders. | 21 | 25 | 16% | 63% | 67% |
| | Others | 55 | 1270 | 30% | 68% | 7% |
| $ARC.KEGG_{both}$ | $GenAge_{Hum}$ | 49 | 84796 | 2% | 17% | 10% |
| | $GenAge_{Mod}$ | 35 | 41304 | 4% | 25% | 9% |
| | *Diseases* | 22 | 16233 | 3% | 23% | 11% |
| | Immune Disorders. | 8 | 227 | 5% | 4% | 2% |
| | Others | 17 | 9356 | 5% | 29% | 11% |

Regarding the clustering coefficient, the $ARC.COX_{90}$ network presented the highest level of clustering, followed by $ARC.COX_{95}$, then $ARC.PPI$, and finally, the $ARC.KEGG$ network exhibited the lowest average clustering. The *Others* group tended to show the highest clustering coefficient across all groups of genes, while $GenAge_{Hum}$-associated genes displayed relatively low values, with $GenAge_{Mod}$ being slightly higher. Immunological disorders-related genes maintain the lowest value in this term in the $ARC.KEGG$ network, whereas they showed a relatively standard value, relative to the remaining groups of the same network, in the $ARC.PPI$, and $ARC.COX$ networks.

The 'Percentage of Disease-related Neighbours' column indicates the mean percentage of the genes interacting with the reference gene that is associated with at least one ARC. This measure is however, biased, as most *Disease*-related genes are predominantly associated with the cardiovascular and endocrine ARCs. The analysis reveals that $GenAge_{Mod}$-associated genes consistently showed about 9% of their connections linked to *Disease*-related genes, with $GenAge_{Hum}$ following a similar trend, albeit slightly higher (+1%) in some cases. However, these $GenAge$-associated genes did not importantly differ from the *Others* group or *Disease*-associated genes in terms of *Disease*-related neighbouring ratio, as they were occasionally even lower, especially in $ARC.COX_{90}$ and $ARC.COX_{95}$. In most networks, immunological disorder-associated genes displayed a lower pepercentage of connections with *Disease*-associated genes than the remaining groups. For example, in the $ARC.PPI$ network, immunological disorder-associated genes were comparable to the remaining groups with about 8% of connections linked to *Diseases*-related genes, slightly less than the 10% seen in the remaining groups of this network. In the $ARC.KEGG$ networks, however, the immunological disorders-related genes showed only 2% of network connections with *Disease*-related genes, compared to the next lowest of 9% in *Others*. The $ARC.COX$ networks, and especially $ARC.COX_{95}$, contrasted this trend by highlighting the prominence of immunological disorders-related genes at connecting with genes associated with *Diseases*-related genes, outscoring any of the other groups and networks in this matter. However, as observed in Figures S7 and S9, this high scoring was due to almost exclusively clustering with other Immunological disorder-related genes, but not other *Disease*-related genes with lower *Pleiotropy*.

#### Proximity analysis

We define Distance as the number of edges that need to be transited from a Reference Gene to reach a Target Gene following the shortest path. In this context we created a Gene-Gene Distance matrix using the “distances” command of the *igraph* library, where we consider genes at the rows as the Reference genes and genes at the columns the Target Genes. When no genetic path between the two genes exist, the Distance is considered Infinite. If a Gene is itself the Reference and Target Gene, then the distance is zero. In the context of a specific ARC, we define Distance as the shortest path connecting the Reference Gene to the closest gene associated to such Target ARC (see Figure S13). Our analysis requires computing the average distance to multiple ARCs. However, this is not possible if a Reference Gene is totally unconnected to at least one ARC (i.e., lacking paths to such ARC genes) because the Distance would turn infinite for such ARC, disrupting calculations when averaging to Distances to all ARCs. To cup with this, we mapped the mapped the Distance to a normalized measure that we call Proximity and is computed as:

$$Proximity = \frac{1}{Distance + 1}$$

Note how Proximity is inversely proportional to Distance and evaluates, from zero to one, the closeness of the Reference Gene to the Target ARC based on the shortest path to its closest gene. The term “+1” prevents infinites at Distances of zero (i.e., when the gene is self-references). Proximity of zero arises when Distance is Infinite. Proximity then increases as Distance decreases. When the Distance is zero, Proximity is 1.

In Figure S13, we illustrate the concept of Proximity with a hypothetical graph, where the Gene G4 (red node) is taken as Reference from where measuring Distance/Proximity to a given ARC or ARD of interest. The genes G0, G6 and G9 (green nodes) are genes associated to such ARC/ARD. There is an individual Distance/Proximity to each of such genes. Depending on the analysis we take either the closest one – the one with the highest Proximity (Figure S14, based on ARCs; and the *ClosestsGene* algorithms described later in this Supplementary document -.  Prediction of human ageing-related genes), based on either ARC/ARD) – or average all Proximities associated to such ARC/ARD-related genes (*AverageGene* algorithms, described later in methods).

For each gene, we computed the *Closest.Proximity* to each ARC. Subsequently, we averaged the proximity quantities towards all ARCs, yielding an average proximity per gene to all ARCs. When depicting Figure S14, we aggregated all genes associated with each of the five groups, and for each of those groups, we compiled the previously obtained average Proximity, forming a distribution of average proximities to all ARCs per gene corresponding to each group. The $GenAge_{Hum}$ and $GenAge_{Mod}$ groups corresponds to *GenAge* genes in humans and model organisms, the *Disease* group corresponds to genes associated with at least one disease and also part of the respective genetic interaction network, meaning they interact with at least one gene within the respective network. *Neighbours* are genes adjacent to disease genes without being the disease genes

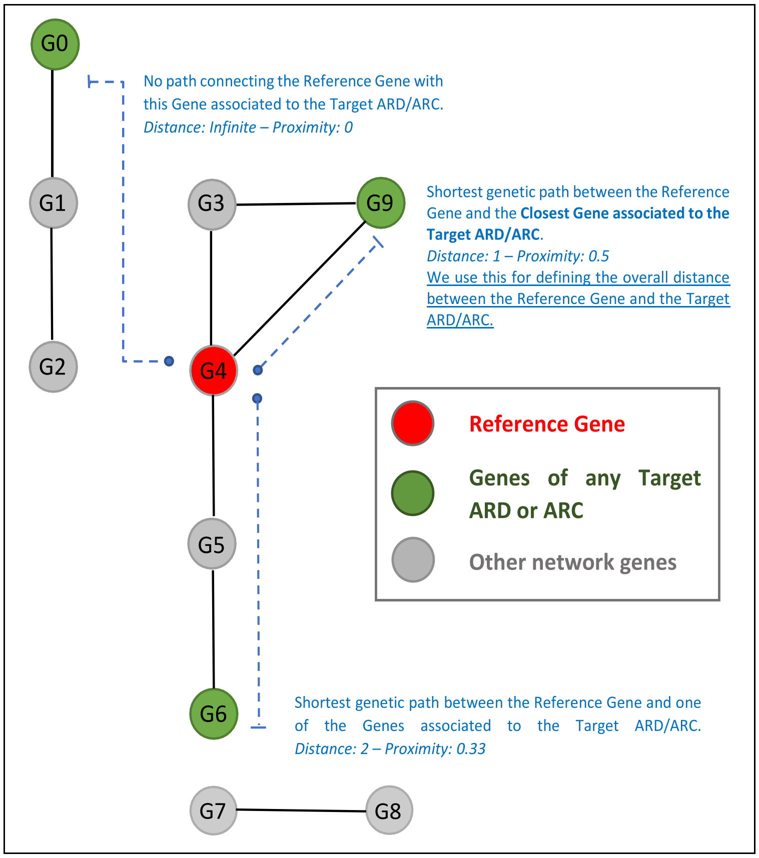


**Figure S13.** Representation of Proximity. A hypothetical PPI/COX/KEGG network with genes G0, G6 and G9 associated to one particular ARD or ARC is presented. Distance to the ARD is measured from gene G4. The shortest genetic path between the Reference Gene and the Closest gene associated with the Target ARD is chosen to define the overall Distance. As for this hypothetical network, the distance from the Reference Gene G4 to the Target ARD's genes is Infinite with respect to G0, 2 with respect to G6 and 1 with respect to G9. Of these, the Distance between the Reference Gene and G9, Distance of 1, is taken as representative of the Distance between the reference gene and the ARD. This procedure is repeated for each gene in the network and for all ARDs. Different ARDs will have different associated genes.

themselves, and the *Others* group comprises all remaining genes within the network. Statistical differences were computed using t-test given their relatively Gaussian distribution across groups.

#### Proximity Analysis

We conceived the metric Proximity that quantifies the inverse of the network distance between genes and ARCs based on shortest paths. For a specific gene, a Proximity of one to an ARCs means direct association to it. A proximity of 0.5 means one intermediate gene to the closest ARC-related gene. Increasing intermediate genes result in lower proximities and when no genetic paths exist in between, the number of intermediate genes is considered infinite and the Proximity zero. We then averaged this metric for all ARCs per gene. In the proximity analysis, genes were categorized into five groups: $GenAge_{Hum}$, $GenAge_{Mod}$,

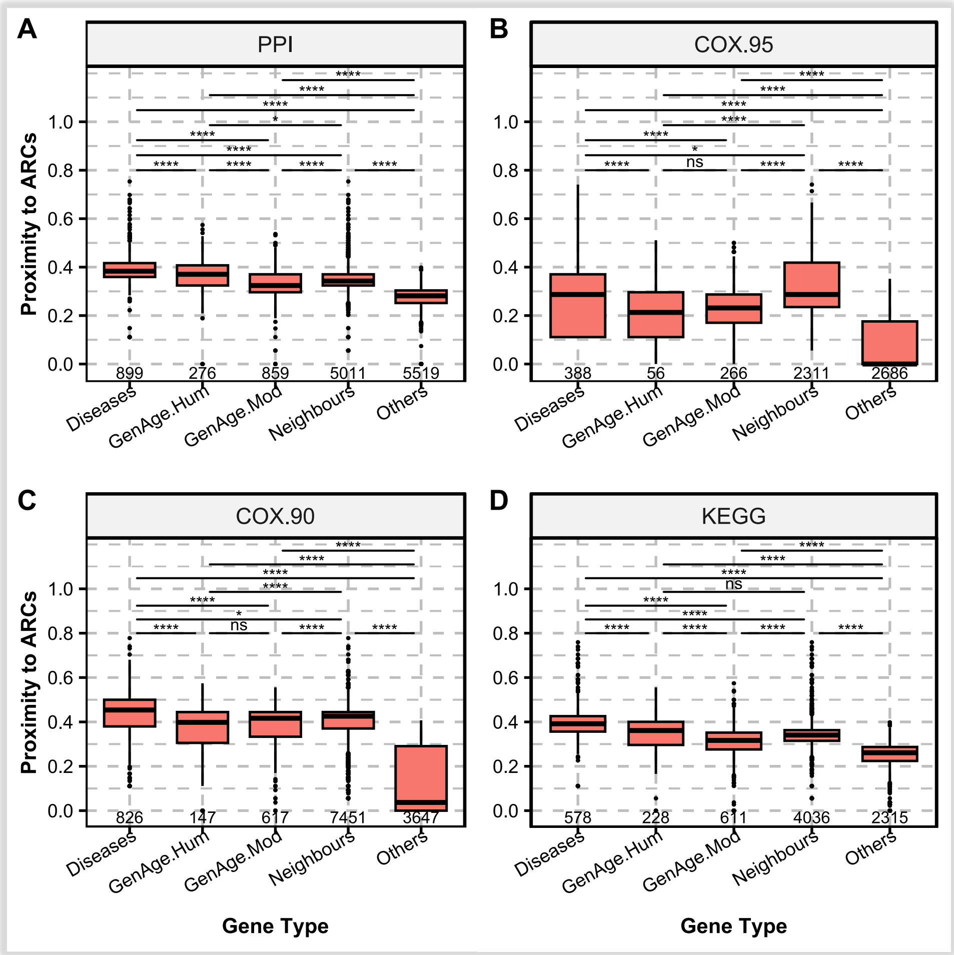


**Figure S14.** Proximity density across the following sets of genes: $GenAge_{Hum}$, $GenAge_{Mod}$, *Diseases*, *Neighbours*, and *Others* (*i.e.*, the remaining genes within each corresponding interaction network). **A**. $ARC.PPI$ network. **B.** $ARC.COX_{95}$ network. **C.** $ARC.COX_{90}$ network. **D**.$ARC.KEGG$. Statistical differences were computed using t-test and adjusted for multiple tests using Bonferroni. Note that, in *Diseases*, we measured the average proximity of each ARC-related gene to all ARCs (e.g., the distance of a cancer related gene to all ARCs, including cancer itself: cancer, cardiovascular, endocrine/diabetes, etc).

*Diseases* (ARCs), *Neighbours* (i.e., Neighbour genes of *Disease*-related genes), and *Others* (i.e., the remaining genes in the network), as depicted in Figure S14.

In the $ARC.PPI$ and $ARC.KEGG$ networks, $GenAge_{Hum}$-associated genes showed a proximity to ARCs almost equivalent to that of the *Disease* group, despite limited direct disease connections. This proximity was slightly closer than the *Neighbour* group, a trend that was echoed in the KEGG network, albeit with weak significance. However, a notable divergence appeared in the coexpression networks, where the *Disease* group was significantly closer to ARCs compared to $GenAge_{Hum}$. In contrast, $GenAge_{Mod}$ demonstrated a significantly lower proximity to ARCs in these networks, a marked difference from the higher values observed for $GenAge_{Hum}$. In the $ARC.COX$ networks, particularly $ARC.COX_{95}$, this trend continued with both $GenAge_{Hum}$ and $GenAge_{Mod}$

showing no significant differences in proximity to ARCs, and their relationship with the remaining gene groups in these networks largely mimicked each other. Thus, while $GenAge_{Hum}$ maintained a marginally closer position to ARCs compared to *Neighbours* in the $ARC.PPI$ network, $GenAge_{Mod}$ held a notably lower proximity in comparison to all groups except *Others.*

A consistent trend across all networks was that the *Others* group scored always a significantly lower proximity to ARCs than the *GenAge* groups, as well as than *Diseases* and *Neighbours*; suggesting that ageing-related genes in both humans and models are significantly closer to ARCs than random genes, regardless of the network. The *Others* group at $ARC.COX_{90}$ and $ARC.COX_{95}$ got several genes with zero proximity, especially at $ARC.COX_{95}$ where the median was biased towards zero. As previously shown in Figures S7 to S10, this isolation in $ARC.COX$s can be attributed to the segmented nature of these networks with multiple clusters, potentially isolating some genes from *Diseases* genes, whereas the number of isolated genes in $ARC.KEGG$ resulted negligeable (Figure S12).

#### Correlation between Pleiotropy and *iARC_Interactions*

Figure S16 depicts the correlation between the spectrum of both Pleiotropies and *iARC_Interactions* across all the ARC-networks. The first column of all sub-figures highlights genes lacking direct ARC affiliations (*Pleiotropy_0*), yet capable of engaging in differing degrees of *iARC_Interactions*. It can be observed that this column contained most of the genes for all networks. Moreover, there was a systematic reduction in gene count as the number of *iARC_Interactions* increases at *Pleiotropy=0* in $ARC.PPI$ and $ARC.KEGG$. An exception to this trend occurred at the first column of the $ARC.COX$ networks, which displayed a relatively consistent gene density across multiple *iARC_Interaction* levels when *Pleiotropy=0*. In particular, $ARC.COX_{90}$ presented its highest density of genes at *Pleiotropy=0*, *iARC_Interactor=6*; whereas all the other networks did it at the minimal *iARC_Interactor=1* when *Pleiotropy=0*.

In most networks, a general trend was observed where the number of genes decreases as function of an increased number of *iARC_Interactions*, regardless of the *Pleiotropy* level, with the only exception mentioned in the last paragraph. Similarly, but to more pronounced extent, there was a

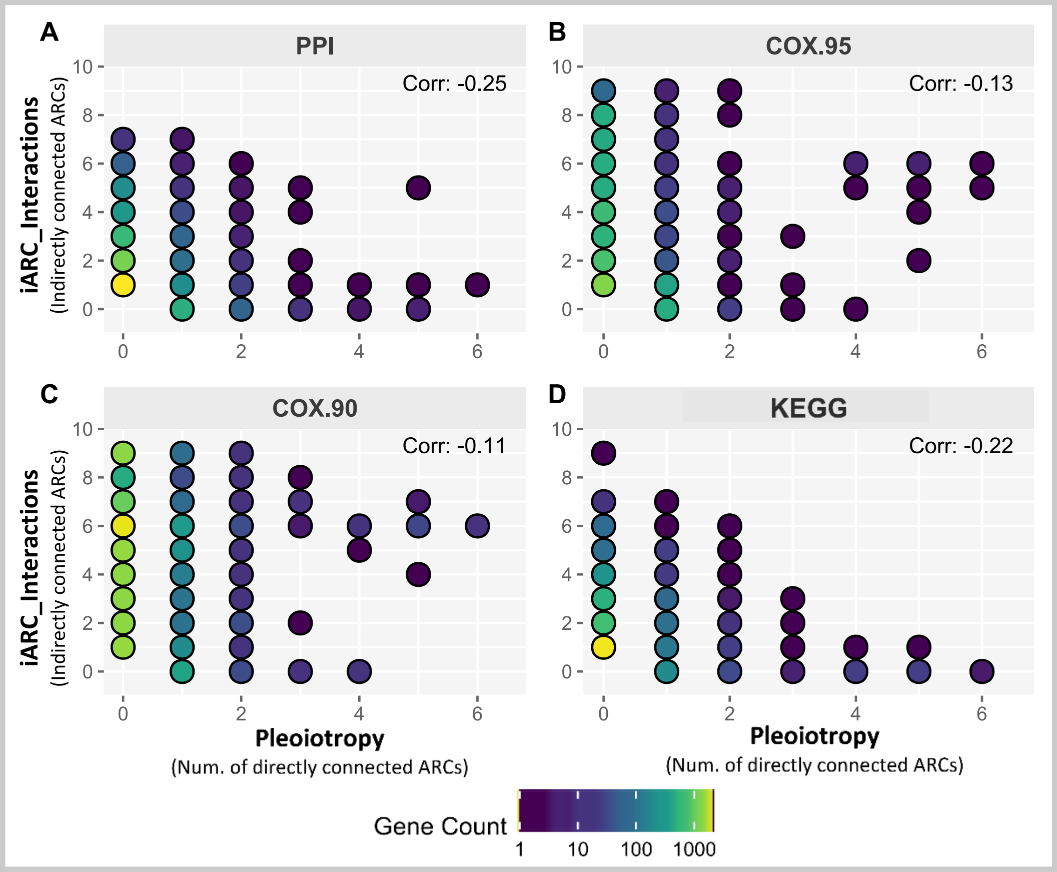


**Figure S15.** Correlation between Pleiotropy and *iARC_Interactors*. Direct and Indirect connections to ARCs (i.e., Pleiotropies and iARC_Interactions, respectively) across all genes on the six different networks used in this study. Each point in this chart, denoted by a circle, represents a unique pairing of *Pleiotropy* and *iARC_Interactions* values, with the circle's colour indicating the number of genes associated to each point. Correlation scores are written on each panel. **A.** $ARC.PPI$ network. **B.** $ARC.COX_{95}$ network. **C.** $ARC.COX_{90}$ network. **D.** $ARC.KEGG$.

decrease in the number of genes as the *Pleiotropy* level increased, regardless of the *iARC_Interaction* level. In the $ARC.PPI$ and $ARC.COX$ networks, the spectrum of *iARC_Interactions* was maintained in the range of 0-2 *Pleiotropies*, mostly allowing for six or more *iARC_Interactor* levels. The $ARC.KEGG$ network, on the other hand, showed an early reduction of *iARC_Interactions*, which became more pronounced with increased *Pleiotropies.* Beyond the 2$^{nd}$ level of *Pleiotropy*, and particularly after the 4$^{th}$, the reachability of high *iARC_Interactions* sharply declines in all networks. This effect was most evident in the $ARC.PPI$ and $ARC.KEGG$ networks, where genes that directly connected with 3 or more ARCs often got only one degree of *iARC_Interaction*. $ARC.KEGG$ networks demonstrated the most stringent trend, with *Pleiotropy_6* genes typically lacking *iARC_Interactions*. $ARC.COX$ networks, slightly deviated from this pattern, allowing genes with higher levels of *Pleiotropy* (5 to 6) to maintain a relatively high number of *iARC_Interactions* (6 to 7).

Overall, as depicted in Figure S15, the correlation between *Pleiotropies* and *iARC_Interactions* was negative in all cases. $ARC.KEGG_{both}$ had a correlation of -0.22, roughly on par with the $ARC.PPI$ network at -0.25. The $ARC.COX$ networks showed the least negative correlations, with -0.13 ($ARC.COX_{95}$), and the lowest -0.11 ($ARC.COX_{90}$). Thus, there was a relative trend where lower *Pleiotropies* were more common at higher levels of *iARC_Interactions* and vice versa, more strongly maintained the $ARC.PPI$ and $ARC.KEGG$ networks, and to a lesser extent in the $ARC.COX$s.

#### Specificity and self-coexpression

Figure S16 indicates that the higher self-coexpression of *GenAge*-related genes with respect with the other groups as observed in Figures 4 and 5A is significant, with $GenAge_{Mod}$-related being even significantly higher than $GenAge_{Hum}$. Renal diseases and *Pleiotropy_6* genes were not significantly different from the *GenAge* groups but mostly due to their small number of genes. Urology and hematology genes generally showed no significant self-coexpression differences compared to other ARCs, with some exceptions in haematology. The Figure S16 also illustrates that the lower coexpression value previously observed for Immunological disorders-related genes relative to the two *GenAge* groups and most ARCs was significant, except for renal, urological, and gastrointestinal diseases. Regarding the gene *Pleiotropy* sets, no significant self-coexpression differences were observed between immunology disorders-related genes and *Pleiotropies* at levels 4 and 5, as expected since they are the same set of genes, but there is a minor difference with *Pleiotropy_6* genes which demonstrated higher self-coexpression, deviating from the patterns of decreasing coexpression started from *Pleiotropy_4*. Generally, genes associated with Pleiotropy at levels 1, 2, and 3 showed significantly higher self-coexpression than groups of genes with higher *Pleiotropy*.

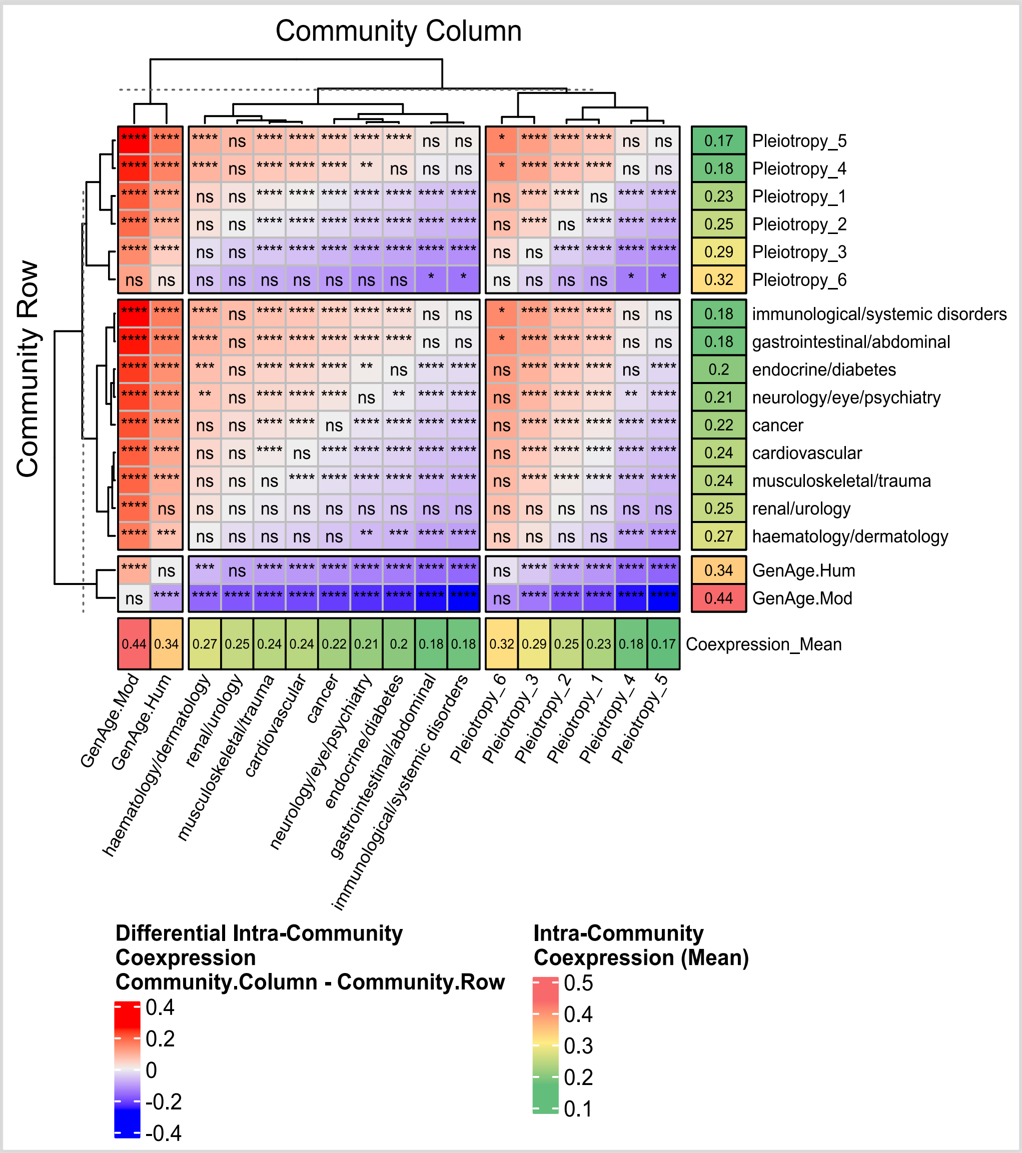


**Figure S16**. Intra-community coexpression differences. **I**ntra-community coexpression differences across various groups, whether they be *GenAge* groups, ARCs, or genes associated with different degrees of *Pleiotropy*. Additionally, outside the primary heatmap, the lowest annotation row and rightmost column each provide another heatmap, reflecting the intra-community coexpression of each community through the various columns and rows respectively. Colouring within the map indicates differential amounts, computed by subtracting the column group value from the row group value, representing a mean difference. Asterisks denote the significance level of this difference, which could also be non-significant, and these significance differences were computed utilizing a t-test with Bonferroni correction for multiple testing. For instance, to examine whether $GenAge_{Hum}$-related genes have higher intra-community coexpression compared to cancer genes, one would reference the $GenAge_{Hum}$ column and Cancer row, thereby considering the difference of $GenAge_{Hum}$ (column mean) minus Cancer (row mean). In this scenario, a reddish colour near 0.15 indicates a highly significant difference, demonstrating that $GenAge_{Hum}$-associated genes tend to significantly exceed most other groups. Notably, *Pleiotropy*_6 genes also tended to exceed some groups but not significantly, attributed to their scarce quantity (12 genes). *Pleiotropy_4_5* genes, along with immunological disorder genes associated with this *Pleiotropy*, are observably significantly lower than the majority of other groups in terms of coexpression.

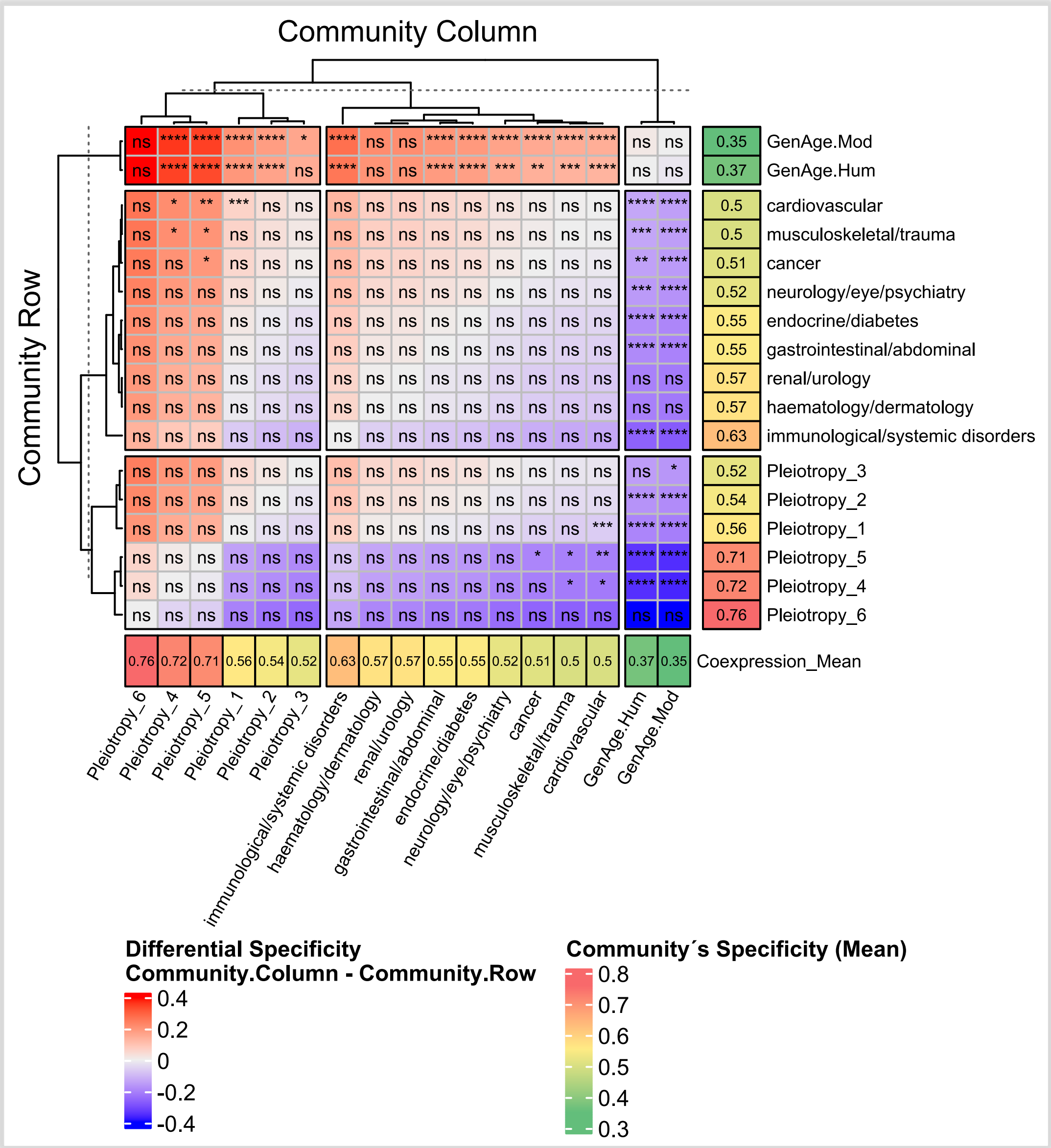


**Figure S17**. Corrected Specificity differences. Tissue specificity differences across various groups are displayed, whether they be GenAge-related genes, ARCs-related genes, or genes associated with different degrees of *Pleiotropy*. The difference is calculated by subtracting the specificity indicated in the column (noted in the bottom row) from the specificity of the community (denoted in the far-right column). This subtraction results in the colors visible in the heatmap: red signifies positive differences (where the column value exceeds the row), blue denotes negative differences, and white indicates no difference. Text within each cell represents the significance level of the observed mean differences, which was assessed using the Wilcoxon test due to the non-Gaussian distribution of the data and corrected with Bonferroni.The differences were mostly non-significant for the majority of communities after Bonferroni correction for multiple tests upon the use of the Wilcoxon test due to non-gaussian distributions. Broadly, the GenAge groups tend to have significantly lower specificity than most other groups.

Figure S17 indicates that the lower specificity observed at Figure 5B in *GenAge*-associated genes from human and model organisms was highly significant with respect to the majority of groups, with the exceptions being the renal community where there was no significant difference and the hematological community where the significant difference was weak. It is also noted that there was no significant difference in tissue specificity levels among different ARCs after correcting for multiple tests. Therefore, although the tissue specificity of immune disorders was greater, it is not necessarily significantly higher than that of other ARCs, though it was significantly higher than the two sets of *GenAge*-associated genes. Regarding the *Pleiotropies* section, no notable significant differences were observed between them either.

The $ARC.PPI$ network displays strong significant differences in tissue specificity across all categories of *iARC-Interactors*, except for the minor but still significant difference between *iARC-Interaction_1* and *iARC-Interaction_4+*. Tao values decrease subtly and sequentially as indirect pleiotropy increases ranging from a mean of 0.27 in *iARC-Interaction_1* to 0.20 in *iARC-Interaction_4+*. The number of associated genes also decreases as the level of indirect pleiotropy increases. Moreover, the distributions of Tao values display median values that lies relatively at the middle of the interquartile range for all *iARC-Interaction* groups.

#### ARC-related gene expression across tissues

Figure S18 presents a description of ARC-related gene expression across various tissues. It was found that generally, there is no dramatic difference in expression levels, although specific tissues associated with certain ARCs showed slight variations. Cancer-related genes displayed no significant expression distinctions across associated tissues like breast, skin, colon, and prostate. Cardiovascular disease genes had slightly lower expression in heart tissue, but not in blood vessels, while gastrointestinal genes showed a minor decrease in pancreas and liver. Renal disease genes exhibited a broad expression variance, with no notable differences in kidney, urethra, and bladder. Endocrine diseases saw lower expression in the pancreas, with minor differences in the liver and thyroid. Neurological disease genes had similar patterns across tissues, with nerve tissue slightly higher. Muscle tissue for musculoskeletal diseases underexpressed marginally, and hematological/dermatological diseases showed no major differences, although blood had a lower median. . Immunological disorder-related genes typically have lower median expression levels in all tissues, with notable variance and lower mean expression in blood, despite its importance in immunity. This pattern is more pronounced than in other tissues, such as the pancreas, kidney, liver, and heart. The GTEx database lacks data on specialized immune tissues like bone marrow and thymus.

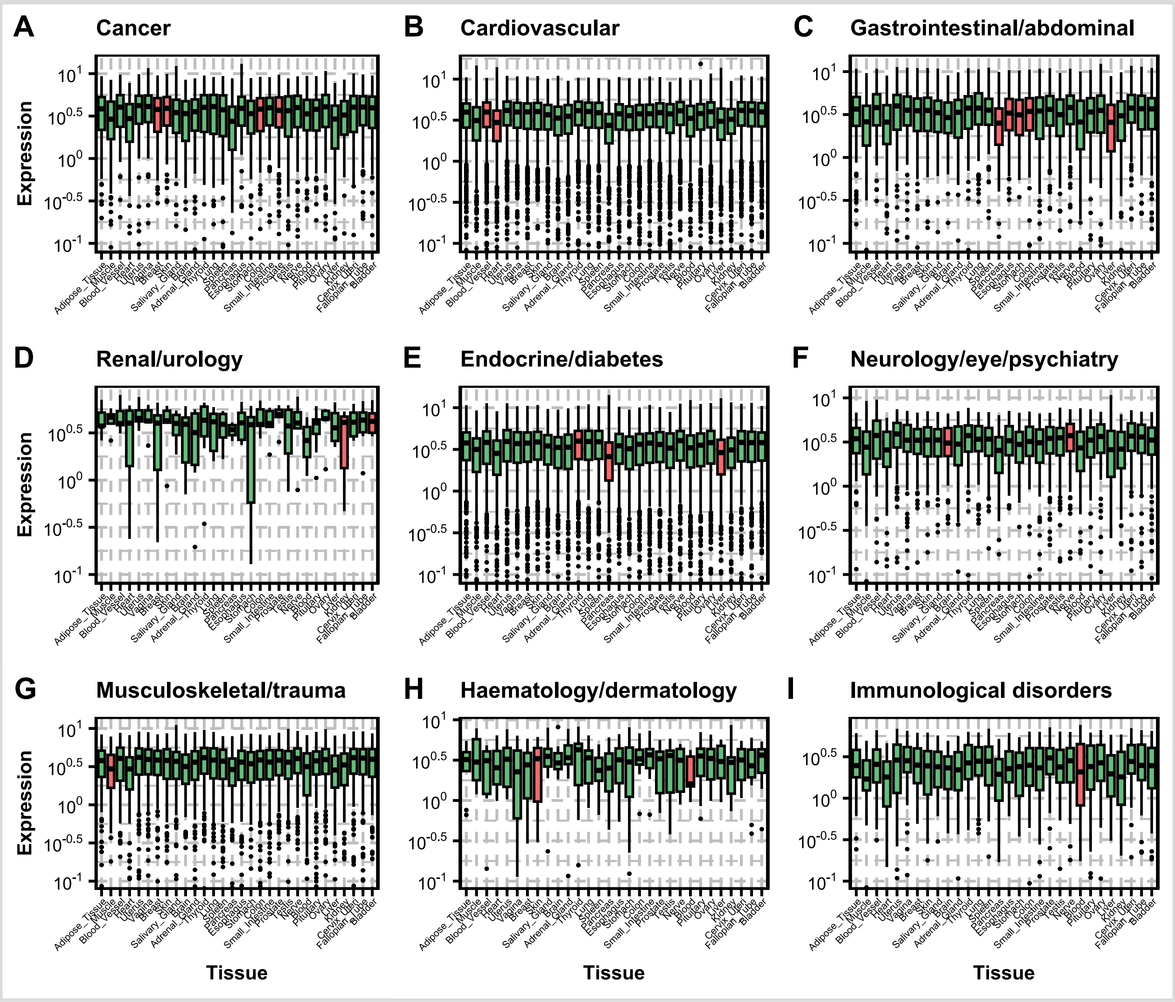


**Figure S18.** Gene expression levels across of various tissues in the GTEx database for genes associated to each ARC. Expression levels were shown on a logarithmic scale. Tissues commonly associated with each ARC were highlighted in red. The X-axis represents the tissues and the Y-axis represents expression levels measures in log-transformed reads per kilobase million (RPKM). **A)** Cancer. The selected tissues were skin, colon, breast and prostate as this ARC was created from the aggrupation of genes associated with cancer in those tissues. **B)** Cardiovascular diseases with heart and blood vessels as associated tissues; **C)** Gastrointestinal/Abdominal diseases, associated with the stomach, colon, liver, small intestine, pancreas, and esophagus. **D)** Renal/Urology diseases, associated with kidney and bladder. **E)** Endocrine/Diabetes, associated with pancreas, liver, and thyroid. **F)** Neurology/eye/psychiatry, associated with brain and nerves. **G)** musculoskeletal diseases, associated with muscle; **H)** Haematological and dermatological diseases, associated with blood and skin. **I)** Immunological diseases, associated with blood.

#### Datasets creation

We conducted 48 independent runs, combining eight algorithms and six networks to utilizes the association of each gene to either ARDs or ARCs to compute ageing-relatedness. The algorithms are as described in Table S4. These values were used as features in a ML procedure. The objective was a binary classification of genes as either $GenAge_{Hum}$- or non-$GenAge_{Hum}$-associated. Genes included in the ML model had at least one connection with another gene within the networks and were either *Disease*-related, *GenAge*-related, or their *Neighbours*. We excluded those without a connection. We did so to focus on network topology to guide predictions and manage data imbalance. Therefore, while all the networks have either 9 or 58 features (ARCs or ARDs), the number of genes being classified and ageing-related genes varied across networks, with specific counts given for $ARC.PPI$ (6721, 276, which are the number of genes under analysis and the number of human ageing-related genes, respectively), $\mathrm{ARC.COX}_{95}$ (2463, 56), $\mathrm{ARC.COX}_{90}$ (7725, 147), and $ARC.KEGG$ (4891, 228); the number of genes being classified diverges to 6721, 2463, 7725 and 4891. A visual description of ARD vs ARC analysis is at Figure S19.

**Table S4** Gene-ARD/ARC proximity algorithms (see Figures S13 and S19).

| Algorithm | Description | Target |
|---|---|---|
| *Closest.Proximity2ARD* | Proximity based on the shortest path between the reference gene and the closest gene of its target ARD. | ARD |
| *Average.Proximity2ARD* | Averaging shortest path proximities between the Reference Gene and each one of the genes of its Target ARD. | |
| *Neighbours2ARD* | Counting the overlapping genes neighbouring the reference gene with the ARD-related genes for each ARD. | |
| *Closest.Proximity2ARC* | Proximity based on the shortest path between the reference gene and the closest gene of its target ARC. | ARC |
| *Average.Proximity2ARC* | Averaging shortest path proximities between the reference gene and each one of the genes of its target ARC. | |
| *Neighbours2ARC* | Counting the overlapping genes neighbouring the reference gene with the ARC-related genes for each ARC. | |

#### Machine Learning Algorithm & Nested Cross Validation

Balanced Random Forests from the python's *imblearn* library (Lemaître, 2017) was chosen for ML due to the imbalanced classification challenge arising from a low count of ageing-related genes relative to non-ageing. Predictive performance was assessed using a nested cross-validation (CV) process. The genes were divided into 10 stratified outer folds. Each ML algorithm underwent 10

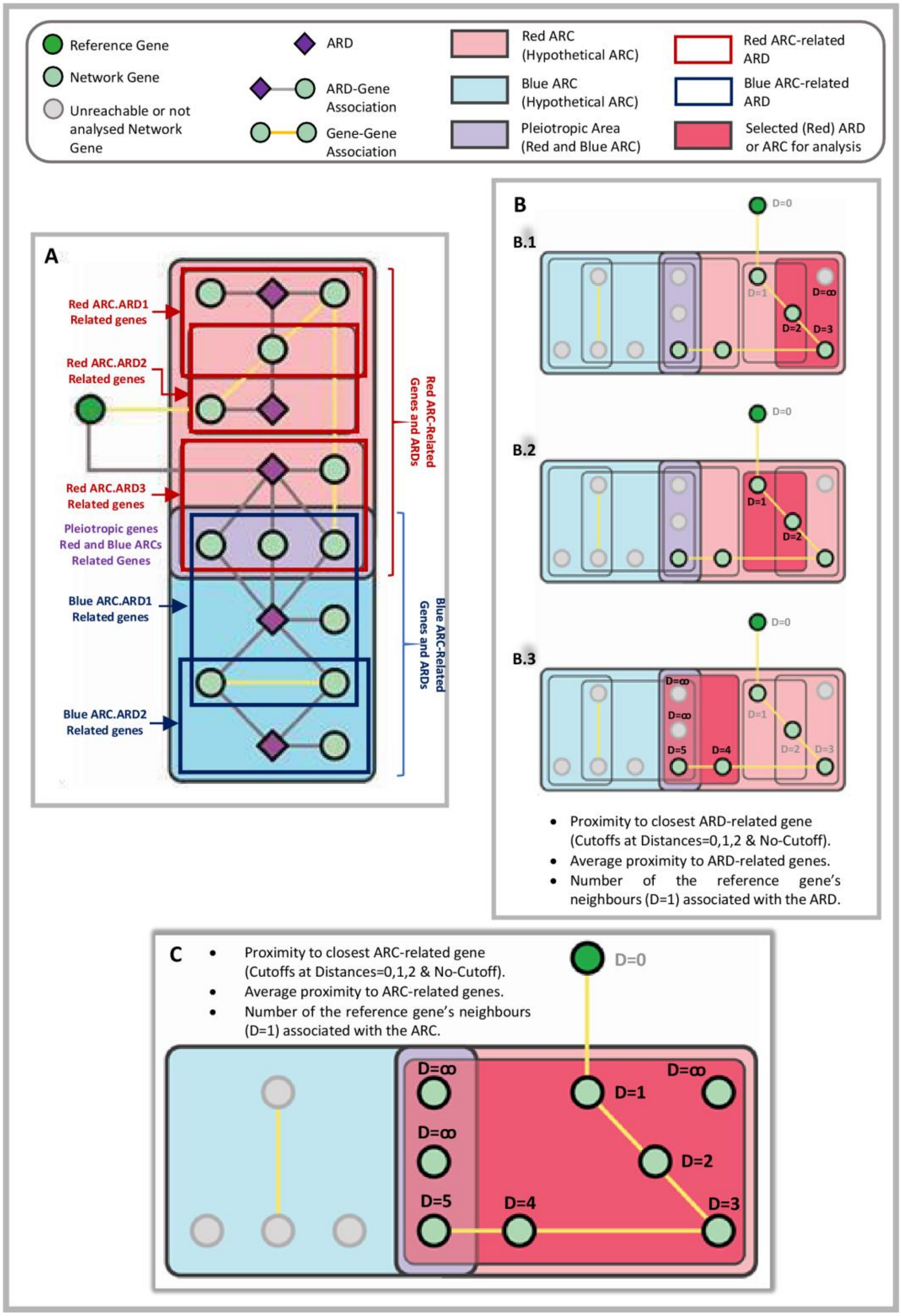


**Figure S19.**. Proximity-based ML-features definitions. Measure of distances between reference genes (green circles), phenotypes (purple diamonds), and communities: **A**. Displays a schematic of gene-to-phenotype connections (grey) and gene-to-gene connections (yellow), representing two hypothetical communities (Red and Blue) and the intersections between them (Purple). There is also an external reference gene (strongest green circle). **B**. Three illustrations indicating the distance between the reference gene and the genes associated with the phenotype of interest, highlighted by a red rectangle. The purple diamonds are omitted, and the calculation is based solely on the gene interaction network and the phenotype-associated genes. The ARD-related features values are assigned to the reference gene as explained in Table 3.8 for algorithms *ARD_ClosestGene* (considering its thresholds), *ARD_AverageGene* and *ARD_Neighbouring*. **C**. ARC-related feature value assigned to the reference gene, similar to that in Figure C, but considering the entire community instead of individual phenotypes. The ARD-related features values are assigned to the reference gene as explained in Table 3.8 for algorithms *ARC_ClosestGene* (considering its thresholds), *ARC_AverageGene* and *ARC_Neighbouring*.

iterations, each using one different outer fold for testing and the rest for training. Before each ML run, the hyperparameters were tuned using an inner 5-fold cross-validation, ensuring the best configuration was selected for the current iteration of the outer CV. The overall algorithm's predictive accuracy was derived as the average of the 10 outer CV iterations. Predicted probabilities were converted into class labels using a 0.5 threshold.

*Human Ageing-related Genes Prediction & Enrichment Analysis:* Although predictions were binarized, the Ageing-related probabilities served as an indication of the model's confidence level. For instance, two genes, A and B, with probabilities of 0.6 and 0.9, would both be classified as Ageing-related. However, gene B is perceived as more reliably ageing-related by the model. We aimed to identify novel Ageing-related genes by spotting the False Positives (FP) among those annotated as Not-Ageing-related but having a high predicted probability of being ageing-related. These FP genes could be potential candidates for future experiments. Combining predictions from all outer testing splits, we focused on FP genes with a probability ≥ 0.5. We selected the top FP genes with probability score above 0.9 from the best models of two different Networks as candidate ageing-related genes for further lab validation. The criterion was chosen to ensure consistent gene numbers across networks for enrichment analysis, and these genes were then used for enrichment analysis on the *gProfiler* website (Raudvere et al., 2019).

### Prediction of human ageing-related genes and pathways

We predicted human ageing-related genes ($GenAge_{Hum}$-associated) based on the six previously established networks. Different methods were employed, each using a metric of relationship with diseases at either the ARC or ARD levels to generate ML features (thus either 58 ARD-related features for some datasets while 9 ARC-related features for other datasets). The features were then used to predict associations to $GenAge_{Hum}$-related genes using two different approaches, one based on ML and another purely based on averaging the values of all features at the corresponding dataset (i.e., pure proximity-based prediction). We call this last approach *Avg* as a short for averaging-based.

Although in-depth procedures were outlined in the previous section, here we briefly explain the methodologies used to assign feature values to the relationships between genes and diseases at either the ARD or ARC levels. We do so as we display the performance results across networks in Table S5 but its nomenclature requires a basic understand of the algorithms.  The methodology comprises several strategies, and includes: 1. Our already stablished concept of proximity, based on the shortest path between the reference gene and the closest gene associated to its target ARD or ARC (*ARD_ClosestGene* and *ARC_ClosestGene*, respectively). 2. The same concept of (1) but introducing distance thresholds of zero, one or two (*Thr0*, *Thr1*, *Th2*) to cap path lengths. 3. Averaging the proximity between the reference gene and each one of the targets ARD- or ARC-associated genes (*ARD_AverageGene* and *ARC_AverageGene*, respectively). 4. counting the overlapping genes neighbouring the reference gene that are ARD-related or ARC-related genes (*ARD_Neighbouring* and *ARC_Neighbouring*, respectively).

#### Performances of the human ageing-estimating algorithms.

The prediction scores across all the combinations of networks and prediction algorithms is described in Table S5. The Aurea Under the ROC Curve (AUC) was used as prediction metric. In short, it was observed that, upon the use of ML, $ARC.PPI$ scored the highest (0.80), followed closely by $ARC.KEGG$ (0.77). $ARC.COX_{95}$ scored below (0.70), whereas $ARC.COX_{90}$ achieved the poorest performance (0.60). Such highest scores generally took place when defining features by the *ARD_ClosestGene* algorithm without capping thresholds.

Looking at the *Mean* column in Table S5 we find that, although individual combinations of network and algorithms can achieve relatively high AUC (e.g., 0.80) the mean values of the *Avg* and *ML* approaches were lower (at much 0.65 and 0.70, respectively). Moreover, the *ML*-based scores did generally better than the A*vg*-based by 0.02 to 0.09 AUC, with the only exception of $ARC.COX_{95}$ where the *Avg*-based scores got 0.03 greater. Looking at the Mean row at the bottom of Table S5 we found that, upon computing the Mean value of all the ML scores across the different algorithms of each network, the best-performing networks were the $ARC.PPI$ and the $ARC.KEGG$, scoring 0.71 each. The $ARC.COX$ networks generally performed worse, with $ARC.COX_{90}$ scoring the lowest of all networks (0.57), and $ARC.COX_{95}$ (0.63) the least scoring pathway-based network. On the other hand, $ARC.PPI$ and $ARC.COX_{95}$ consistently outperformed other networks from the *Avg*-scores point of view. The *Avg* scores at the $ARC.PPI$ network, in particular, even outscored or met the highest ML-based prediction value of other networks except $ARC.KEGG$, whose *Avg*-scores were low but the ML-based scores were high relative to most networks.

**Table S5.** AUC results depicting the $GenAge_{Hum}$-relatedness prediction performance of each of the networks across each of the proposed relationships with ARDs and ARCs. The 'ML' sub-columns represent the predictions performances when features are processed with ML. The 'Avg' text is short for "Average" and its associated sub-columns represent the predictions performances when the mean (i.e., average) value of all the features is used as predictor. A colour gradient from red to green was used to highlight, in red, the strongest performances, while in green the weakest ones. The last column "Mean" depicts the mean value of the "Avg" and "ML" approaches across all the networks. The last sub-column "All" of the "Mean" column, indicates the mean value of the "Avg" and "ML" approaches taken together. The last row "Mean" indicates the mean value throughout the algorithms of the associated sub-column.

| DATASETS | $ARC.PPI$ | | $ARC.COX_{90}$ | | $ARC.COX_{95}$ | | $ARC.KEGG$ | | Mean | | |
|---|---|---|---|---|---|---|---|---|---|---|---|
| ALGORITHM | Avg | ML | Avg | ML | Avg | ML | Avg | ML | Avg | ML | All |
| ***ARD_ClosestGene*** | 0.72 | 0.8 | 0.59 | 0.57 | 0.69 | 0.66 | 0.61 | 0.77 | 0.65 | 0.7 | 0.68 |
| ***ARC_ClosestGene*** | 0.66 | 0.73 | 0.6 | 0.57 | 0.68 | 0.62 | 0.58 | 0.74 | 0.62 | 0.67 | 0.64 |
| ***ARD_AverageGene*** | 0.74 | 0.79 | 0.57 | 0.59 | 0.66 | 0.63 | 0.63 | 0.74 | 0.65 | 0.69 | 0.67 |
| ***ARC_AverageGene*** | 0.74 | 0.76 | 0.57 | 0.61 | 0.68 | 0.62 | 0.64 | 0.7 | 0.65 | 0.67 | 0.66 |
| ***ARD_Neighbouring*** | 0.67 | 0.71 | 0.61 | 0.59 | 0.7 | 0.67 | 0.6 | 0.74 | 0.63 | 0.68 | 0.65 |
| ***ARC_Neighbouring*** | 0.67 | 0.69 | 0.61 | 0.62 | 0.7 | 0.67 | 0.6 | 0.73 | 0.63 | 0.67 | 0.65 |
| **Mean** | 0.66 | 0.71 | 0.57 | 0.57 | 0.66 | 0.63 | 0.58 | 0.71 | 0.65 | 0.68 | 0.66 |

Considering the *Mean-All* column, which presents the mean values of the *Avg*- and *ML*-based scores, the *ARD_ClosestGene* algorithm (0.68) and its second order capped version *ARD_ClosestGene_Thr2* (0.67) generally matched and outperformed other ARD-based algorithms with more complex mechanisms of ARD-relationship, namely *ARD_AverageGene* (0.67) and *ARD_Neighbouring* (0.65). In this direction, *ARD_Neighbouring* matched the performance of its ARC-based equivalent *ARC_Neighbouring* (0.65). In contrast, almost all the other ARD-based algorithms were respectively superior to their ARC-based parallels by a minor – *ARC_AverageGene* (0.66) vs *ARD_AverageGene* (0.67) – or slightly greater – *ARC_ClosestGene_Thr1* (0.62) vs *ARD_ClosestGene_Thr1* (0.64); *ARC_ClosestGene_Thr2* (0.64) vs *ARD_ClosestGene_Thr2* (0.67); *ARC_ClosestGene* (0.64) vs *ARD_ClosestGene* (0.68) – extent. Noteworthy is as well that, while the ARD-based algorithms found their peak performance at the *ARD_ClosestGene* algorithm, outranging more complex ARD-based measures; its ARC-equivalent, the *ARC_ClosestGene* algorithm, did not display this effect, as it achieved equal to lower scores compared to more complex ARC-based algorithms.

Continuing with the *Mean-All* column, it is noteworthy that employing thresholds in the *ARD_ClosestGene* and *ARC_ClosestGene* networks – reveals clear trends. Setting *Thr_0* yielded random predictions (*Mean* of 0.5 and 0.52, for the ARD- and ARC-based respectively). However, *Thr1* showed noticeable improvement (0.64 and 0.62), while the improvement from *Thr1* to Thr2 was subtle (*Mean* of 0.67 and 0.64, respectively). Skipping the thresholds yielded to networks performing relatively equal with respect to *Thr2* (*Mean* of 0.67 and 0.64, respectively). The only exception to this trend was $ARC.COX_{95}$ whose ML-scores at *Thr1* in both the ARD- and ARC-based versions, remarkably outperformed *Thr2* and the threshold-less versions.

The top performing scores across the PPI- and KEGG-based networks converged in the use of ML and the *ARD_ClosestGene* algorithm. Namely, $ARC.PPI$, $ARC.KEGG$, with AUC of 0.80 and 0.77, respectively. The top performing scores across the $ARC.COX$ networks also converged in the use of ML but used different algorithms and got from slightly to substantially lower scores than other networks. Specifically, $ARC.COX_{95}$ scored 0.72 when using *ARD_ClosestGene_Thr1*. On the other hand, $ARC.COX_{90}$ lied far below any other network, achieving 0.62 at *ARC_Neighbouring*. Lastly, $ARC.COX_{90}$, although notably low, was the only network reaching its maximal performance from an ARC-based approach.

#### Predicted genes and biological processes.

Tables S6 and S7 display the top 10 genes most likely to be associated with ageing based on the protein interaction network and the combined pathway network. These genes currently have no association with GenAge. Table S8 lists the Biological processes GO terms linked to the top 30 ageing-associated gene candidates from both the protein interaction and combined pathway networks. The number 30 was chosen as it covers less than 1% of genes in both networks while covering a sufficient number of genes for enrichment analysis.

**Table S6.** Top 10 ageing-related gene candidates based on $ARD.PPI$. The column "Rank" indicates the hierarchy of the genes across the candidates based on the ML-inferred Ageing-relatedness probability. Direct and indirect pleiotropy to ARCs and ARDs is displayed.

| Rank | Gene | Ageing-relatedness Probability | Pleiotropy (ARCs) | iARC_Interactors (ARCs) | Pleiotropy (ARDs) | iARD_Interactors (ARDs) |
|---|---|---|---|---|---|---|
| 1 | HSPB1 | 0.968 | 0 | 4 | 0 | 15 |
| 2 | GNL3 | 0.958 | 1 | 4 | 1 | 11 |
| 3 | RAF1 | 0.958 | 1 | 3 | 1 | 9 |
| 4 | KAT5 | 0.948 | 0 | 5 | 0 | 15 |
| 5 | UBC | 0.948 | 0 | 5 | 0 | 18 |
| 6 | TMEM106C | 0.946 | 0 | 1 | 0 | 1 |
| 7 | KPNB1 | 0.936 | 0 | 2 | 0 | 4 |
| 8 | VHL | 0.936 | 0 | 5 | 0 | 19 |
| 9 | HSP90AB1 | 0.934 | 0 | 4 | 0 | 9 |
| 10 | NPM1 | 0.932 | 0 | 4 | 0 | 14 |

**Table S7.** Top 10 ageing-related gene candidates based on $ARC.KEGG$. The column "Rank" indicates the hierarchy of the genes across the candidates based on the ML-inferred Ageing-relatedness probability. Direct and indirect pleiotropy to ARCs and ARDs is displayed.

| Rank | Gene | Ageing-relatedness Probability | Pleiotropy (ARCs) | iARC_Interactors (ARCs) | Pleiotropy (ARDs) | iARD_Interactors (ARDs) |
|---|---|---|---|---|---|---|
| 1 | PIK3CD | 0.979 | 0 | 6 | 0 | 22 |
| 2 | MAPK11 | 0.970 | 0 | 7 | 0 | 23 |
| 3 | MAPK12 | 0.970 | 0 | 7 | 0 | 23 |
| 4 | PIK3R2 | 0.961 | 0 | 6 | 0 | 20 |
| 5 | MAPK10 | 0.947 | 0 | 5 | 0 | 14 |
| 6 | AKT2 | 0.933 | 0 | 5 | 0 | 11 |
| 7 | PRKCB | 0.919 | 0 | 5 | 0 | 20 |
| 8 | FOXO6 | 0.915 | 0 | 4 | 0 | 12 |
| 9 | ATF4 | 0.915 | 0 | 6 | 0 | 15 |
| 10 | MCHR1 | 0.896 | 1 | 0 | 1 | 0 |

**Table S8.** Biological processes GO terms associated with the top 30 ageing gene candidates of the $ARC.PPI$ and $ARC.KEGG_{Both}$ networks.

| Network | GO term | Description | Number of Genes (out of 30) | Adjusted pvalue |
|---|---|---|---|---|
| $ARC.PPI$ | GO:0051246 | Regulation of protein metabolic process | 22 | 2.32E-11 |
| | GO:0050821 | Protein stabilization | 8 | 6.49E-07 |
| | GO:0071495 | Cellular response to endogenous stimulus | 13 | 4.51E-05 |
| | GO:0033044 | Regulation of chromosome organization | 7 | 8.26E-05 |
| | GO:0009628 | Response to abiotic stimulus | 11 | 0.000382 |
| | GO:0030878 | Thyroid gland development | 3 | 0.015345 |
| | GO:0051403 | Stress-activated MAPK cascade | 5 | 0.033487 |
| $ARC.KEGG_{Both}$ | GO:0051094 | Positive regulation of developmental process | 18 | 1.17E-11 |
| | GO:0070887 | Cellular response to chemical stimulus | 22 | 6.93E-11 |
| | GO:0002551 | Mast cell chemotaxis | 3 | 0.000939 |
| | GO:0043551 | Regulation of phosphatidylinositol 3-kinase activity | 4 | 0.00192 |
| | GO:0050926 | Regulation of positive chemotaxis | 3 | 0.01095 |
| | GO:0050900 | Leukocyte migration | 6 | 0.02945 |
